# Blind genomic tree scans identify loci underlying adaptive peaks in Antirrhinum

**DOI:** 10.1101/2025.02.12.637406

**Authors:** Daniel M. Richardson, Desmond Bradley, Lucy Copsey, Annabel Whibley, Monique Burrus, Christophe Andalo, Sihui Zhu, Hilde Schneeman, David L. Field, Yongbiao Xue, Enrico Coen

**Affiliations:** Department of Cell and Developmental Biology, John Innes Centre, Colney Lane, Norwich, NR4 7UH; School of Biological Sciences, University of Auckland, Auckland, New Zealand; Centre de recherches sur la biodiversité et l’environnement CRBE, CNRS, UMR 5300, Université Toulouse III Paul Sabatier, F-31062, Toulouse, France; Center for Genomics and Biotechnology, Fujian Provincial Key Laboratory of Haixia Applied Plant Systems Biology, Key Laboratory of Genetics, Breeding and Multiple Utilization of Corps, Ministry of Education, Fujian Agriculture and Forestry University, Fuzhou, 350002, China; Institute of Science and Technology Austria, Am campus 1, 3400 Klosterneuburg, Austria; Applied BioSciences, Macquarie University, NSW 2109, Australia; Institute of Genetics and Developmental Biology; Chinese Academy of Sciences, Beijing, China; Xianghu Laboratory, Hangzhou 311231, China

## Abstract

A key question in evolutionary biology is how barriers to gene flow operate and arise between populations, leading to speciation. Barriers are typically identified by comparing populations distinguished by phenotype, geographical location and/or environment. However, such prior classifications may bias recovery towards features obvious to the human eye, influencing mechanistic interpretations. Here we develop a method blind to prior classifications. We apply it to 18 Antirrhinum populations occupying habitats from coastal to alpine. By scanning the genome for regions with deeply rooted similar trees, we identify a single multi-locus partition, comprising less than 0.5% of the genome. It derives from eight genes, three newly identified here, which interact to generate complementary pollinator guides, likely corresponding to two adaptive peaks in a fitness landscape. Hybrid genotypes fall in a fitness valley, allowing the partition to be maintained through hybrid incompatibility and creating steep clines in allele frequency at a hybrid zone. Another partition derives from a single locus linked to a gene controlling plant height. The locus exhibits a shallow cline in allele frequency across an eco-geographic barrier and may be eco-adaptive. Our results suggest that two ancestral subspecies underwent extensive gene flow except at a few barrier loci, showing how both hybrid incompatibility and eco-adaptation can operate to maintain diversity.

## Introduction

A fundamental problem in evolutionary biology is how barriers to gene flow arise and act between populations, leading to speciation^1,2,3^. One limitation of previous studies is that they typically involve comparisons between pairs of populations with clear phenotypic differences, so they may bias analysis towards traits that are readily observed, influencing mechanistic interpretations. For example, populations may be subdivided in multiple ways not apparent to the human eye, challenging hypotheses based on a small number of ancestral subspecies. The bias may be considerable as less than one third of genes may give obvious phenotypes when mutated^4,5^. A further limitation is that by using relative divergence (F_ST_) as the primary metric for pairwise genome scans, barrier loci may be confounded with loci undergoing differential selective sweeps^6^.

The limitations of pairwise comparisons have been addressed by using phylogenetic methods to compare multiple taxa simultaneously^7,8,9,10,11^. However, these methods can only be applied to a small number of taxa, typically subsets of four, selected based on phenotype, ecology, or geography.

Here we develop a blind approach that allows trees of 20 or more taxa or populations to be compared, using absolute divergence (d_XY_)^12^ as the primary metric, which is insensitive to selective sweeps when calculated over large genomic regions. Once population partitions and candidate barrier loci have been identified, phenotypes, ecological relations and mechanisms underlying reduced gene flow can be assessed. We apply this method to populations of *Antirrhinum* sampled across a wide range of habitats and consider the implications of our findings for genotype, phenotype and fitness spaces.

*A. m. majus* var. *striatum* has yellow flowers with a magenta highlight that signals the pollinator entry site, whereas *A. m. m.* var. *pseudomajus* has a complementary pattern of magenta flowers with a yellow highlight^13^. These varieties form narrow hybrid zones (∼1km wide), allowing barrier loci which exhibit steep clines in allele frequencies to be identified^14^. The loci include three major flower colour genes: *ROSEA* (*ROS*), *ELUTA* (*EL*) and *SULFUREA* (*SULF*). *ROS* and *EL* encode MYB transcription factors that control the distribution of magenta pigment^15,16,17^; whereas *SULF* encodes small RNAs that destabilise transcripts of a yellow pigment biosynthetic gene^18^. More recently, two further barrier loci have been identified through F_ST_ or cline scans: *FLAVIA* (*FLA*), which affects yellow and is the target of *SULF*^19^, and *RUBIA* (*RUB*), which affects magenta and likely encodes flavonol synthase^20^.

Populations of both varieties are found in a broad range of environmental conditions, from coastal to alpine. It is unclear whether hybrid zones exist for multiple other traits and partitions, with flower colour only having been identified because of its salience to geneticists and taxonomists. It is also unclear whether flower colour clines are maintained through hybrid incompatibility, and/or through differential adaptation to non-obvious environmental differences across the hybrid zone.

Here we apply genomic tree scans to 18 populations (9 striatum and 9 pseudomajus) sampled across their geographic range. We identify forests with deep-rooted trees that highlight candidate partitions and barrier loci. Two forests stand out. One divides the populations according to flower colour. This partition is accounted for by the five previously identified flower colour loci as well three further loci *CRE, AUN* and *XAT* identified here. The partition is most likely maintained through hybrid incompatibility between complementary flower colour phenotypes, corresponding to two peaks on a fitness landscape. The other partition is accounted for by a single genomic region linked to plant height that shows a shallow cline in allele frequency across an ecogeographic barrier. Our results suggest that two ancestral subspecies underwent extensive gene flow, except at a few loci that underlie distinct adaptive peaks or are eco-adaptive.

## Results

### Whole genome trees are polyphyletic for flower colour

To explore the genomic relationship between *A. m. m.* var. *pseudomajus* and *A. m. m.* var. *striatum*, nine populations of each variety were sampled across the subspecies range (Fig. 1a). A tree based on geographic distance between populations, gave a polyphyletic grouping for flower colour, showing that phenotype was not strongly correlated with proximity (Fig. 1b).

**Figure 1:**
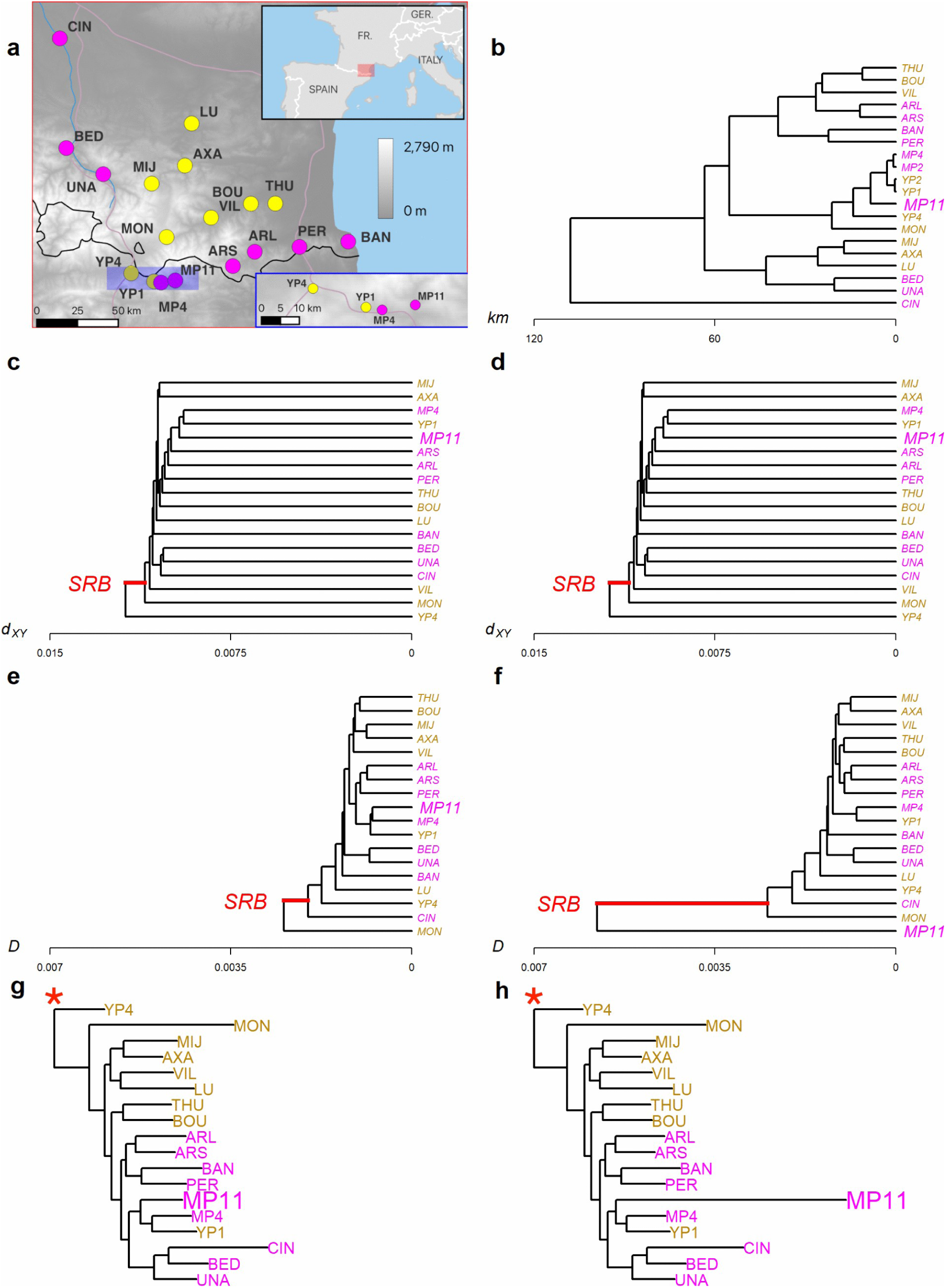
Population locations and whole-genome trees. **a** Topographic map showing the locations of populations of *A. m. m.* var. *pseudomajus* (magenta spots) and *A. m. m.* var. *striatum* (yellow spots) populations. Grey-to-white colour gradient represents altitude (metres above sea level). The hybrid zone region is shaded in blue and shown in the lower inset map. The upper inset map shows the broader study location (black box). GPS coordinates and population sizes are shown in Supplementary Table 1. **b** Tree constructed using pairwise geographic distance between populations, based on GPS coordinates of population sampling sites. Magenta labels are *A. m.* var. *pseudomajus* populations, and yellow labels are *A. m. m.* var. *striatum* populations. **c** Whole-genome d_XY_ tree based on pairwise comparison of all populations over all genomic biallelic SNPs. The shortest root branch (SRB) is highlighted in red. **d** Whole-genome d_XY_ tree after simulating a selective sweep across all polymorphic sites in the MP11 population. **e** Whole genome D tree based on pairwise comparison of all populations over all genomic biallelic SNPs. **f** Whole genome D tree after simulating a selective sweep across all polymorphic sites in the MP11 population. **g** Maximum likelihood whole-genome tree, rooted on an *Antirrhinum sempervirens* outgroup, based on consensus sequences of genomic SNPs. Red asterisk indicates the root, based on outgroup *A.sempervirens* (not shown). **h** Maximum likelihood tree generated after simulating a selective sweep in the MP11 population.

To construct genomic trees, DNA was sampled from each population, pooled within populations, and mapped to the *Antirrhinum majus* reference genome^21^ (Supplementary Table 1). Whole-genome trees based on d_XY_ (equation 2 in Methods) or Nei’s D (equation 4) were generated by taking the average of each statistic across all SNPs in all pool comparisons and generating a UPGMA hierarchical clustering tree. Both d_XY_ and D trees gave polyphyletic grouping for flower colour, but with different tree topologies (Fig. 1 c and e). We also generated a maximum likelihood tree using the consensus sequence for each population and rooting the tree with *Antirrhinum sempervirens*^22^. This tree grouped all pseudomajus populations within a single clade that also included one striatum population (YP1) (Fig. 1g).

Theoretically, d_XY_ trees should be insensitive to sweeps or inbreeding over a large genomic region^6,12,15^. To confirm this expectation, we simulated a genome-wide selective sweep in one population, MP11, by randomly sampling SNPs at each position and setting their frequency to 1. The d_XY_ tree showed no change, whereas in the D tree, MP11 was shifted to become the outgroup (Fig. 1 d and f). Topology of the maximum likelihood tree was unchanged, but the terminal branch length of the swept population, MP11, increased (Fig. 1h). Similar results were obtained with sweeps for other populations, though sometimes tree topology as well as the terminal branch length was changed in maximum likelihood trees (Supplementary Fig. 1a, b, c). Thus, in accordance with theory, d_XY_ trees offer the possibility of identifying candidate barrier loci without systematic confounding effects caused by sweeps or inbreeding.

### Genome scans identify forests with deep root divisions

Consider an ancestral population that becomes split in two through a geographical and/or ecological barrier. The d_XY_ tree for the two populations has an initial height equal to the ancestral within-population divergence, π_w_ (Fig. 2a, D *=* 0). Further assume, for simplicity, that within-population divergence does not change over time. For genomic regions that do not flow between the populations, tree height will increase with time due to the accumulation of mutations in the separated lineages (Fig. 2b). Sub-populations sampled across the species range will give bifurcating d_XY_ trees, with root branch length (RB) depending on duration of divergence (Fig. 2c). For genes that freely flow between the populations, trees have short root branches (Fig. 2d). Thus, scanning the genome for regions that give deeply divided d_XY_ trees provides a route to identifying potential barrier loci.

**Figure 2:**
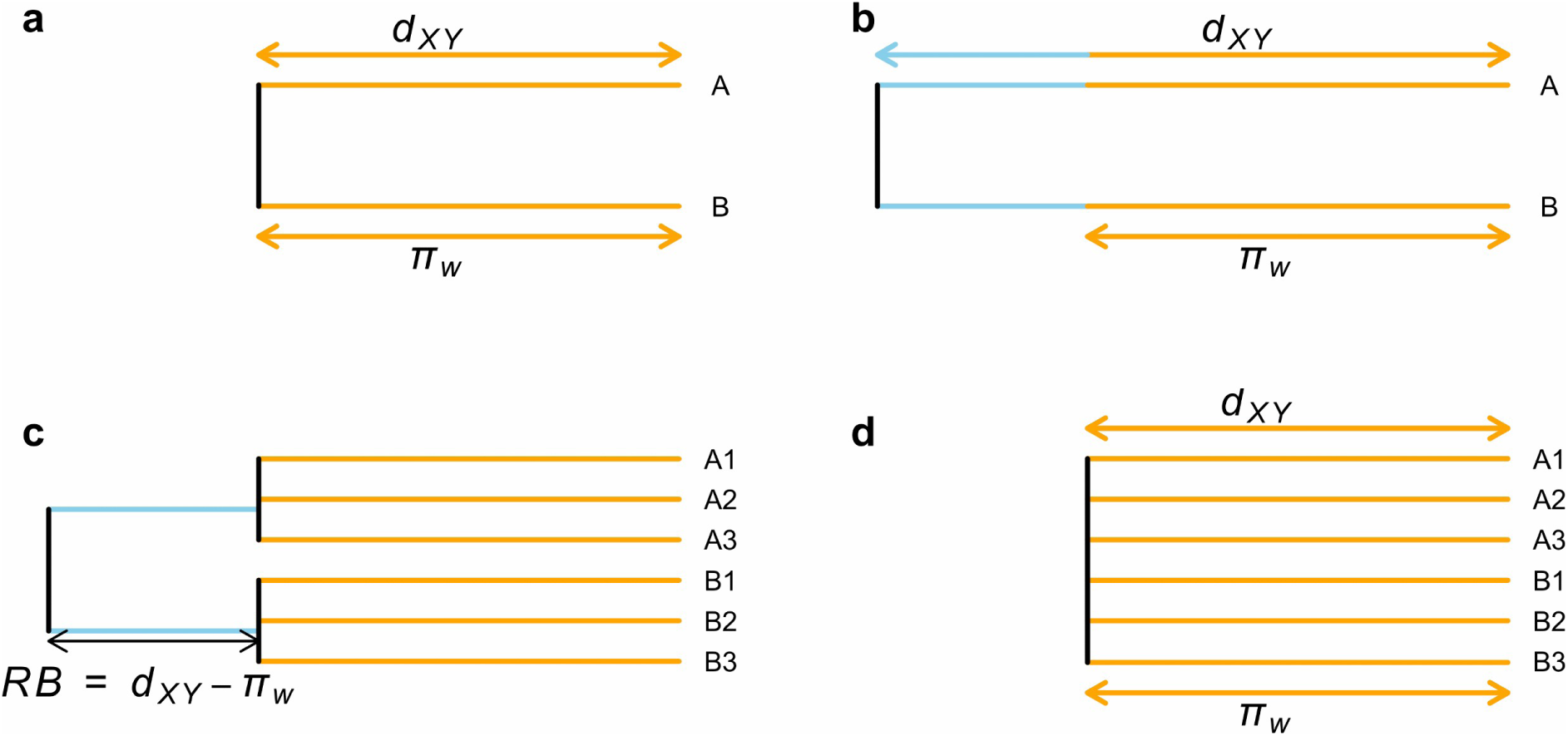
Simplified d_XY_ trees for genes that are isolated or flow between populations. **a** Immediately following isolation, diversity between the A and B populations (d_XY_) equals within-population diversity (π_w_, orange segments). **b** For regions that do not flow between A and B populations accumulation of mutations leads to an increase in d_XY_ above π_w_ (difference in blue). We assume π_w_ does not change over time. **c** Subdivision of populations leads to a d_XY_ tree with root branch (RB) length equal to d_XY_ minus π_w_. **d** For genes that flow freely between populations, d_XY_ equates to π_w_.

To perform such a scan, genomic windows need to be sufficiently large to give reasonable estimates of d_XY_. Given the mean SNP density is 0.031 we chose 50 kb windows to provide about 1,500 SNPs for d_XY_ estimates. A 25 kb overlap between adjacent windows yielded 18,556 genomic windows. Trees for each window could be classified according to how they partitioned the 18 populations based on their root division (Fig 3a). About 85% of trees gave a 1:17 division, with a single population as an outgroup, while other divisions such as 2:16 or 3:15 etc were observed at lower frequency. Each of these divisions may contain multiple partitions. For example, with the 1:17 division there are 18 possible partitions, each with a different outgroup. For the 2:18 division there are 18 x17 possible partitions. We were particularly interested in partitions with deep roots, with more than one population in each partition, as these might reflect barrier loci.

**Figure 3:**
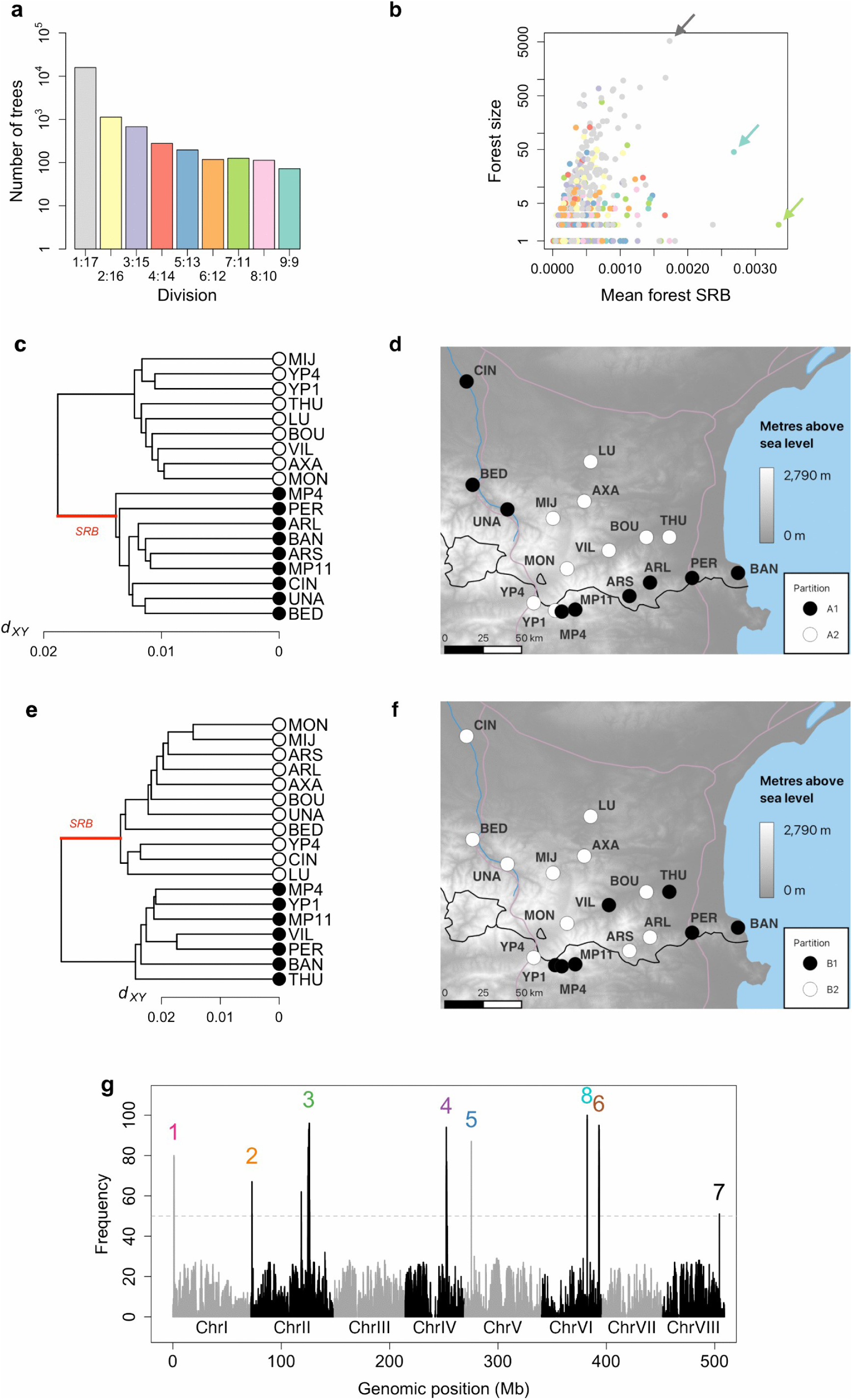
Identification and analysis of partition islands. **a** Histogram showing the number of 50 kb window trees corresponding to each division on a log scale. **b** Forests from a genomic d_XY_ tree scan, with the number of trees within each forest (forest size) plotted against the mean shortest root branch (SRB). Forest colour reflects the divisions shown in (a). The partition A outlier forest (turquoise), partition B outlier forest (green), and whole-genome partition forest (grey) are arrowed. **c** Mean of all trees appearing within the partition A outlier forest. Root division of the tree defines two population groups: A1 (black) and A2 (white). **d** Geographic distribution of groups A1 and A2. **e** Mean of all trees appearing within the partition B outlier forest. **f** Geographic distribution of partition B groups B1(black) and B2 (white). **g** Frequency with which different genomic regions occurred within outlying forests (mean SRB >= 0.002), across 100 tree-scan bootstrap replicates. Each labelled peak corresponds to a region present in over 50% of replicates (grey dotted line). Chromosome indicated by alternating grey-black. Trees summarising the results of the bootstrap analysis are given in Supplementary Fig. 2.

To identify such partitions, we first grouped trees into forests according to their similarity based on the cophenetic correlation coefficient^23^, termed *c* (equation 5 in Methods). Calculating *c* for all pairwise combinations of the 18,556 trees would be computationally prohibitive. We therefore used progressive elimination in which a seed tree was randomly selected and compared with all other trees. All trees with *c* above a threshold value were assigned to the same forest as the seed tree and removed from further comparisons. Another seed tree was then randomly selected from the remaining trees. This process was iterated until no trees remained. To display the depth of tree division, we calculated the length of the shortest root branch (SRB, highlighted in red in Fig. 1b-e) for each tree (Fig 3b). A high value of SRB indicates a deeply divided tree (the longest root branch was not used to avoid highlighting trees with a single outgroup).

In the first instance we used a *c* threshold of 0.5. We plotted mean SRB length against forest size and colour-coded forests according to the root divisions they gave (Fig. 3b), using the same colour scheme as in Fig 3a. The largest forest (grey arrow), corresponded the whole genome d_XY_ tree (a 1:17 division with YP4 as the outgroup, see Fig. 1c). Two forests stood out as having a high SRB (greater than 0.0025): one with a 9:9 division, comprising about 50 trees (turquoise spot, arrowed), and one with a 7:11 division comprising 2 trees (green spot, arrowed). The mean tree of the 9:9 forest is illustrated in Fig 3c. We refer to the populations separated by this root division as partition A, with one population group denoted A1 (black), and the other A2 (white). 76% of trees within this forest gave this partition or deviated from it by a single population being misgrouped, which we refer to as the A’ partition (Supplementary Fig. 2). The mean tree of the 7:11 forest is illustrated in Fig 3e. We refer to the populations separated by the root division as partition B, with groups B1 (black) and B2 (white). Both trees in this forest gave the same partition. Similar results were obtained for a *c* threshold of 0.4 (Supplementary Fig. 3).

### Identification of partition islands

To determine which genomic regions consistently ended up in forests with high SRB, we ran the tree-scanning algorithm 100 times, with a *c* threshold of 0.5, using independent random seed trees. For each run, we selected forests with SRB >0.002. In more than 50% of the replicates, 83 genomic windows (0.45% of all genomic windows) appeared in these high-SRB forests. 81 of these windows gave partition A, A’, or A’’ (A’’ deviates from A by two populations being misgrouped), and mapped to what we termed partition islands 1-7 (Fig. 3g, Supplementary Tables 2 and 3). Most of these islands contained 1-5 windows, except for island 3, which harboured 53 windows. Two windows gave partition B and mapped to a single chromosome region, denoted island 8. No other 50 kb windows gave the B partition. Similar results were obtained with a *c* threshold of 0.4 (Supplementary Fig. 4). Lowering the SRB threshold to 0.0015 recovered a high density of windows as the largest forest was captured (Supplementary Fig. 5).

To gain a higher resolution picture of the partition islands, we scanned the genome in windows one tenth the size (5 kb), identifying those that gave partitions A, A’, A’’ or B irrespective of SRB value. Windows clustered in regions from approximately 0.03 Mb to more than 1 Mb (Fig. 4; Supplementary Fig. 6). Windows giving A or B partitions were only found at or near the identified islands (purple dots Fig. 4). A cluster of A’ and A’’ windows was observed about 8 Mb to the left of island 3 (Fig. 4d). Given the low recombination rate in this region^19^, it is unclear whether this cluster is a distinct island or part of island 3. All islands contained many coding sequences of diverse predicted functions (Supplementary Fig. 7, Supplementary Tables 4 and 5).

**Figure 4:**
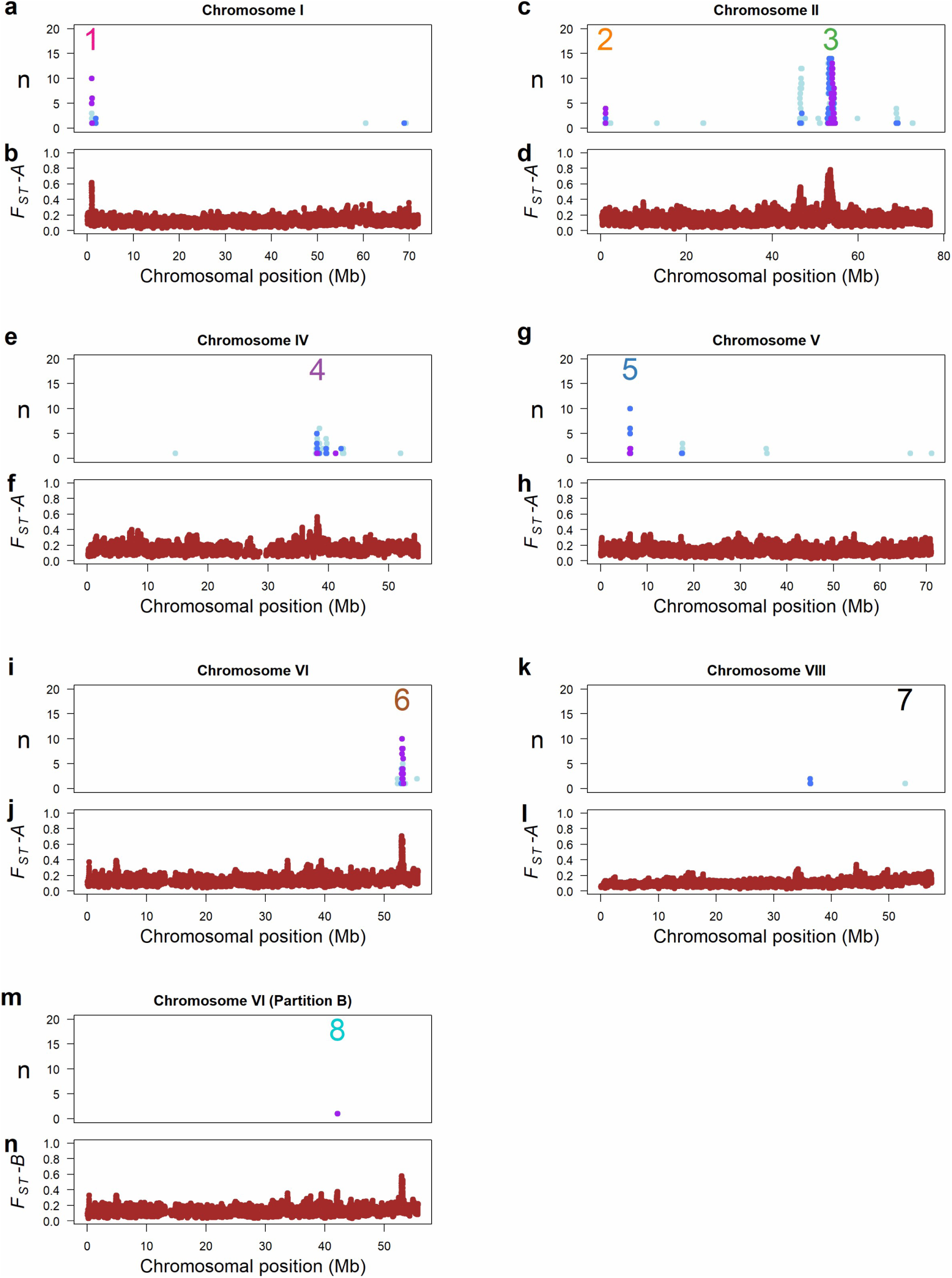
Chromosome scans for monophyletic island density and F_ST_ across eight islands. **a, c, e, g, i, k** Frequencies of 5 kb windows showing the A partition (purple), the A’ partition (one population misgrouped compared to the A partition, dark blue), or the A’’ partition (two populations misgrouped compared to the A partition, light blue) on chromosomes I, II, IV, V, VI, and VIII summed over 50 kb windows with 25 kb overlaps, mapped to the *A. majus* reference genome. Numbers 1-8 indicate the positions of the respective partition A islands. Close-ups of islands are shown in Supplementary Fig. 22. **b, d, f, h, j, l** Mean F_ST_-A for pairwise comparisons of all group A1 and group A2 populations, averaged in 10 kb windows with 9 kb overlaps across chromosomes I, II, IV, V, VI, and VIII. Chromosomes III and VII are shown in Supplementary Fig. 6. **m** Frequency of partition B windows on chromosome VI. **n** Mean F_ST_-B for pairwise comparisons of all group B1 and group B2 populations, averaged in 10 kb windows with 9 kb overlaps across chromosome VI.

### Some but not all islands exhibit F_ST_ peaks

To compare the islands identified by d_XY_ tree scans to those that might have been identified through F_ST_ analysis, we calculated mean value of F_ST_ (3) for all 81 pairwise (9×9) comparisons between A1 and A2 populations across 10 kb genomic windows, with 9 kb overlaps, yielding F_ST_-A. Four of the A partition islands exhibited F_ST_-A peaks, while three (islands 2, 5 and 7) did not (Fig. 4). For partition B, the mean value of F_ST_ for all 77 pairwise (7×11) comparisons between B1 and B2 populations was calculated, yielding F_ST_-B. F_ST_-B did not show a peak at the partition B island (Fig. 4n), though it did show a peak on the same chromosome at the position of island 6, caused by a difference in proportion of A1 and A2 populations in the B partition (4/7 (57%) of B1 populations were A1, whereas 5/11 (45%) of B2 populations were A1).

To understand why some of the partition islands did not show F_ST_ peaks we explored the relationship between F_ST_ and d_XY_ trees. F_ST_ equates to (d_XY_ - π_w_)/ d_XY,_ where π_w_ is the average within-population divergence for the populations being compared. d_XY_ is approximately equivalent to tree height, H, whereas π_w_ to the average length of the terminal branches, L (Fig. 2c, Supplementary Fig. 8a, b). Thus, F_ST_ should have similar values to (H-L)/H. Plotting F_ST_ against (H-L)/H for A partition gave a strong positive correlation as expected (Supplementary Fig. 8c). All windows for the partition-A islands that did not show F_ST_ peaks (islands 2, 5 and 7) had low values of (H-L)/H. A tree scan using F_ST_ as the metric instead of d_XY_ was able to identify island 5, but still not 2 and 7 (Supplementary Fig. 9). Thus, d_XY_ tree scans may be more sensitive than F_ST_ for detecting candidate barrier loci.

### Partition A corresponds to flower colour and islands exhibit clines in allele frequency

Incorporating phenotypic information showed that the A partition divided populations according to flower colour: the A1 group corresponded to *A.m.m.pseudomajus* and the A2 group to *A.m.m.striatum*. Island 3 harbours *FLA*^19^, island 4 harbours *SULF*^18^, island 5 harbours *RUB*^20^ and island 6 harbours *ROS* and *EL*^15,17^. The large size of island 3 (*FLA*) reflects low recombination rates, likely caused by location towards a centromere rather than inversion^19,21^. No flower colour genes have so far been shown to be associated with islands 1, 2 or 7.

To compare clines for all A-partition islands, we plotted SNP allele frequencies against geographic position, for four pools sampled around a hybrid zone^20^. YP4, YP1 are striatum (A2), and MP4 and MP11 are pseudomajus (A1, Fig. 1a, inset)). YP4 and YP1 are separated by an approximately 10 km mountain pass devoid of *Antirrhinum* plants^20^. For each island we chose the top 100 SNPs showing the greatest mean frequency difference between A1 and A2 populations. The SNP frequency in YP4 was subtracted from the frequency in each pool. As a control, we also included 50 kb regions containing *PALLIDA* (*PAL*) and *VENOSA (VE*) genes involved in flower colour but not showing signatures of divergence between populations^15^.

The *PAL* and *VE* gene regions showed no evidence of clines (Fig. 5a,b). By contrast, islands harbouring previously identified flower colour loci (*FLA*, *ROS EL*) showed steep clines across the hybrid zone, evident from bifurcation of SNP frequencies between YP1 and MP4 (SNP alleles were assigned randomly, so 50% should increase and 50% decrease in frequency across a cline). Frequencies of SNPs from *SULF*, *RUB* and islands 1 and 2 showed clines across the hybrid zone (YP1-MP4) and across the mountain pass (YP4-YP1). Island 7 showed weak clines across the mountain pass (YP4 - YP1) but not across the hybrid zone (YP1-MP4), indicating that selection on this locus is weaker than the other loci.

**Figure 5:**
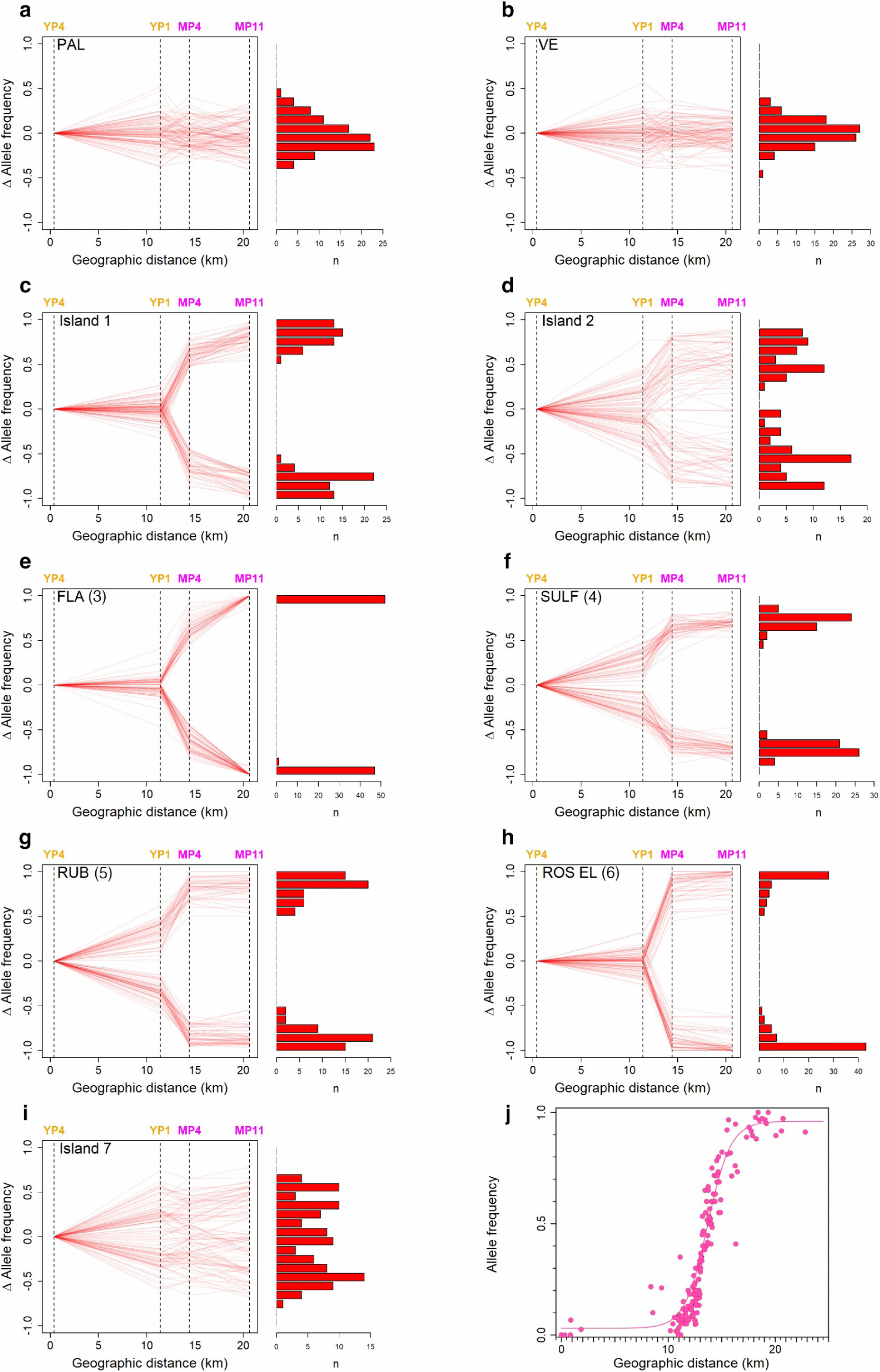
SNP frequencies, relative to the YP4 population, for 100 SNPs across a hybrid zone. Dotted lines indicate geographic position of each population pool (A1 names in magenta, A2 in yellow). For each genomic region, 100 SNPs were selected which showed the highest mean frequency difference between all 18 A1 and A2 populations. For each SNP, allele frequency in YP4 was subtracted from frequency in each pool. Histograms show SNP allele frequencies in MP11 for all 100 SNPs. **a** SNPs from 50 kb genomic region containing the *PALLIDA* coding sequence. **b** SNPs from 50 kb genomic region containing the *VENOSA* coding sequence. **c-i** SNPs from islands 1-7. For named islands, island number is displayed in brackets. **j** Cline in allele frequency for a SNP from partition island 1 (*CRE*), across the hybrid zone based on genotyping of individuals rather than pools.

To measure cline width more accurately for island 1, we determined allele frequencies at the hybrid zone using SNPs^19^ (no SNP data was available for islands 2 or 7). Island 1 SNPs showed a sharp increase in the frequency of pseudomajus alleles from the western to the eastern flank of the hybrid zone (Fig. 5j), centred at a similar location as other flower colour gene clines, supporting the hypothesis that this island harbours a barrier locus.

### Previously unassigned partition A islands harbour or are linked to flower colour loci

To test whether islands 1, 2 and 7 harbour flower colour loci, we analysed flower colour variation in F_2_ populations from crosses between *A. m. m.* var. *striatum* and *A. m. m.* var. *pseudomajus*. To remove colour variation caused by the *ROS*, *EL*, and *SULF* loci, we genotyped for these loci (Supplementary Figs. 11-14). For each genotype, we ranked flowers according to either spread of yellow or magenta intensity and assigned them to high or low bins, each containing 50% of all ranked flowers. We also genotyped for SNPs from islands 1, 2 and 7. SNP allele frequencies for all three islands were significantly different between the high and low yellow bins (Fig. 6a-c) but showed no significant differences between high and low magenta bins (Fig. 6c-e). Thus, islands 1, 2 and 7 either harbour or are linked to yellow flower colour loci.

**Figure 6:**
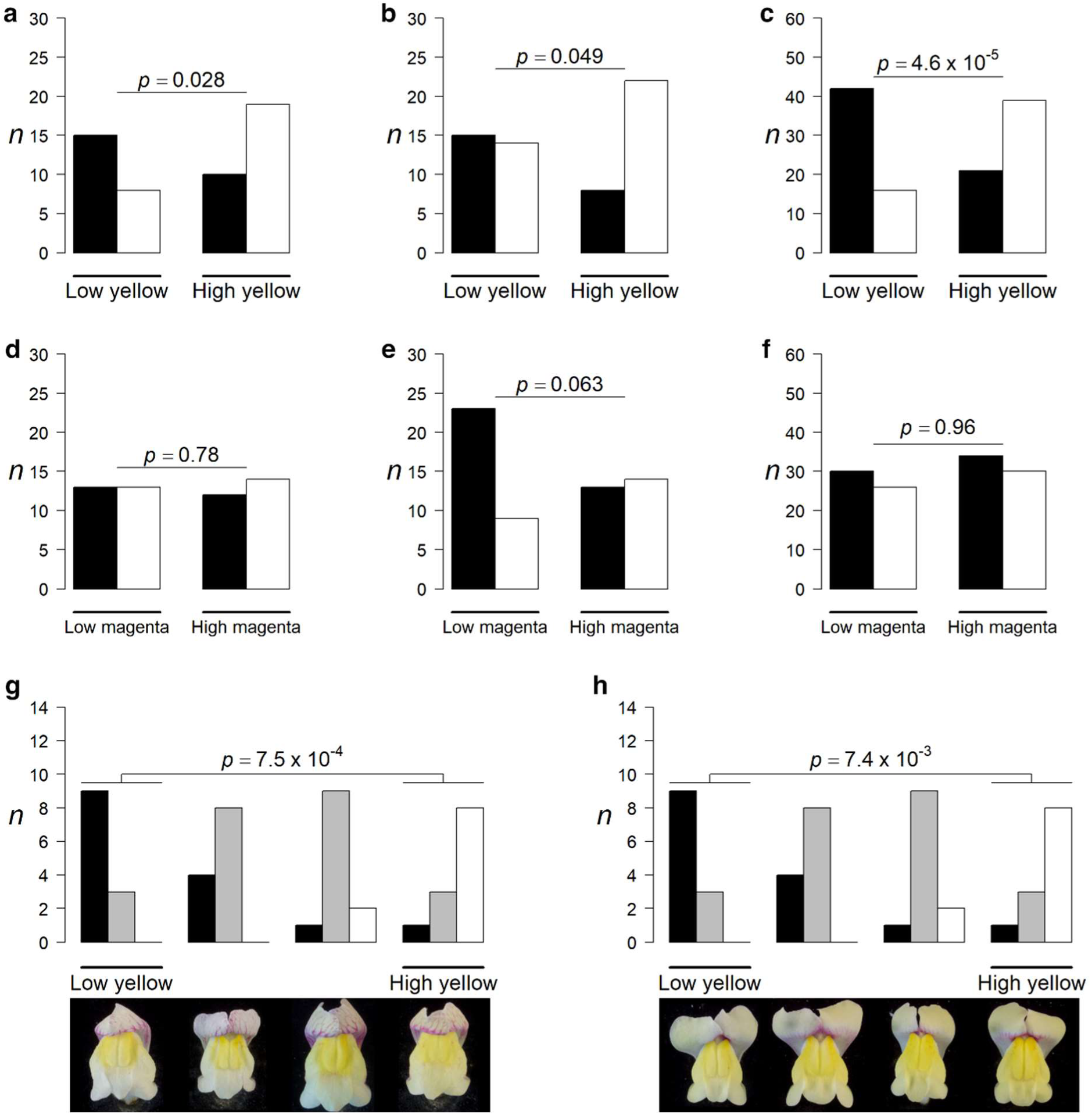
Flower colour ranking for islands 1, 2, and 7. **a** Frequency of island 1 homozygotes from an F_2_ of 164 plants from *A. m. m.* var. *pseudomajus* crossed to *A. m. m.* var. *striatum* (YP4 x Ventolà). Plants were genotyped for island 1, *ROS*, *EL*, and *SULF*. The same *ROS*, *EL*, *SULF* genetic backgrounds were grouped together. Photographs within each group were ranked according to yellow intensity, yielding a low and high yellow bin. White bars refer to the allele that is more abundant in the high yellow bin, black bars to the other allele. The *p*-value is based on a contingency *χ²* test between the low and high yellow groups (n=52). **b** As (a) but for F_2_ plants genotyped for island 2 SNPs. *p*-value based on n=59. **c** As (a) but for 259 F_2_ plants from a cross between a YP4 x Ventolà genotyped for island 7 SNPs. *p*-value based on n=118. **d-f** As (a-c) but with ranking based on magenta colour intensity rather than yellow. **g** Frequency of island 1 genotypes from an F_4_ population with a *FLA^s^/fla^p^ ros^s^/ros^s^ EL^s^/EL^s^* colour gene background. Photographs were ranked according to yellow intensity, yielding four quartiles of increasing yellow. The median photograph from each quartile is shown as representative. *p*-value based on n=24. **h** As (g) but for island 2 genotypes. *p*-value based on n=26. Ranked images are given in Supplementary Figs. 11-20.

To confirm that islands 1 and 2 are linked to yellow flower colour loci, we ranked and genotyped F_4_ populations that were homozygous for *FLA^s^ sulf^s^* ros^s^ *EL^s^* (superscript ‘s’ refers to striatum allele) and segregating for the SNPs at islands 1 or 2 (an equivalent family was not available for island 7). Ranked flowers were sorted into quartiles of increasing yellow intensity. For both islands, the striatum SNP allele frequency was significantly elevated in the highest compared to the lowest yellow quartile (Fig. 6g and h). Both islands showed a deficit of heterozygotes in both the highest and lowest quartiles, indicating that the alleles were codominant. These results were confirmed by replicate rankings, carried out by different observers (Supplementary Figs. 15-20). We named the yellow colour loci linked to islands 1, 2 and 7 *CREMOSA* (*CRE*), *AURINA* (*AUN*) and *XANTHIA* (*XAT*) respectively.

Genes underlying previously identified flower colour loci are differentially expressed between *A. m. m.* var. *pseudomajus* and *A. m. m.* var. *striatum*^17,18^. To identify candidate genes underlying island 1 (*CRE*), island 2 (*AUN*) and island 7 (*XAT*) we therefore performed differential expression analysis using DESeq2^24^ on RNA extracted from two pools (three plants per pool) of glasshouse-cultivated *A. m. m.* var. *striatum* (sampled at Z-ALE, near LU, which is A2, B2), and three pools of *A. m. m.* var. *pseudomajus* (from M-AUT, near CIN, which is A1, B2). Pools were mapped to a pseudomajus genome assembly, to enable detection of indels which may not be present in the reference genome. Out of 45,067 predicted coding sequences, 3,109 (6.9%) were differentially expressed between the pseudomajus and striatum pools (*p*-adjusted < 0.01) (Fig. 7). The high number of differentially expressed loci may reflect the limited number of individuals pooled or consistent differences across the geographic range of the varieties. BLASTN searches against the reference genome suggested that 76 differentially expressed genes (2.4% of 3,109) mapped to partition A islands. 61 of these mapped to islands harbouring previously identified flower colour loci: 30 to island 3 (*FLA*), 18 to island 4 (*SULF*) and 13 to island 6 (*ROS* and *EL*). Two mapped to island 5 (*RUB*).

**Figure 7:**
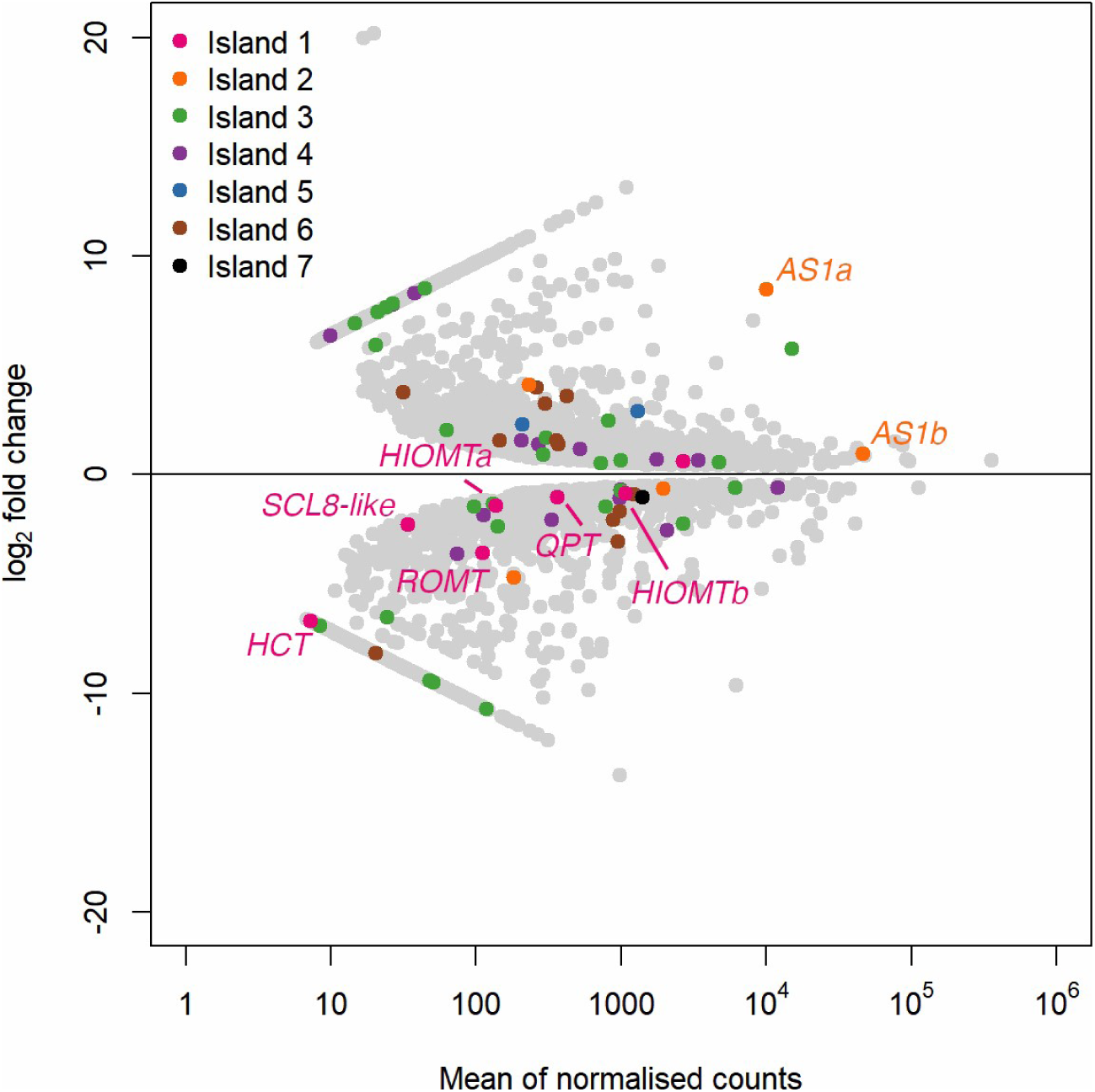
Transcript comparisons between *A. m. m.* var. *pseudomajus* and *A. m. m.* var. *striatum*. DESeq2 differential expression results between two *A. m. m.* var. *pseudomajus* and three *A. m. m.* var. *striatum* pools, each containing three individuals, mapped to an *A. m. m.* var. *pseudomajus* genome assembly. Positive fold changes are elevated expression in *A. m. m.* var. *striatum*, and negative values elevated expression in *A. m. m.* var. *pseudomajus*. All differentially expressed transcripts are shown in grey (*p* < 0.01) with a subset of points coloured according to monophyletic island membership. Island 1 candidate genes from BLAST searches are annotated, along with the island 2 aureusidin synthase transcripts. *AS1a*/*b* = Aureusidin synthase, *HIOMTa*/*b* = Hydroxyindole-O-methyltransferase, *ROMT* = Trans-resveratrol di-O-methyltransferase, *HCT* = Shikimate O-hydroxycinnamoyltransferase, *SCL8*-like = SCARECROW-like-8-like, *QPT* = Nicotinate-nucleotide pyrophosphorylase.

Of the remaining 13 differentially expressed genes, 7 mapped to island 1 (*CRE*). Two encoded hydroxyindole-O-methyltransferases which could influence colour through methylation of pigment precursors^25,26,27,28^. Another encoded a SCARECROW-like transcription factor^29^. Out of the remaining 6 differentially expressed genes, five mapped to island 2 (*AUN*) and one to island 7 *(XAT*). Of those mapping to *AUN*, two were the nearest homologues of the aureusidin synthase gene *AmAS1* in the pseudomajus and striatum assemblies (one of these copies was absent in the reference genome)*. AmAS1* is responsible for synthesising yellow aurone pigments from chalcones^30^ and is therefore a strong candidate for *AUN.* One gene, with similarity to sugar transporter, mapped to island 7 (*XAT*) and could influence flower colour as anthocyanins and aurones are modified through addition of sugar moieties^31,32^. No differentially expressed genes mapped to B-partition island 8, but as both pools were from B2 plants this result is not unexpected.

### Distribution of SNPs at partition A islands

To gain a clearer understanding of how windows and SNPs were distributed at the islands, we mapped their positions in regions spanning the islands. *FLA* (island 3) was the largest island (about 1 Mb), reflecting the low recombination rate of the chromosomal region in which it is located (< 0.1 cM/Mb compared to 3.125 cM/Mb at *ROS*^19^, island 6). A 0.5 Mb region downstream (left) of *FLA* mainly had A’ partition windows (dark blue diamonds, Fig. 8a), while the 0.5 Mb region upstream (right) of *FLA* mainly gave A partitions (purple diamonds). This difference reflected a recombinant *FLA* allele abundant in the YP1 population which borders the hybrid zone^19^ (inset in Fig. 1a). The recombinant allele carries striatum sequence upstream of *FLA* and pseudomajus sequence downstream. The upstream region therefore grouped YP1 with striatum populations (A partition), while the downstream region grouped YP1 with pseudomajus populations (A’ partition). The 100 SNPs from the island showing the highest mean frequency difference across the A partition were clustered around the *FLA* locus (magenta crosses). Similarly, SNPs showing a frequency difference greater than 0.8 across the A partition (yellow crosses, Fig. 8b), were clustered around *FLA*.

**Figure 8:**
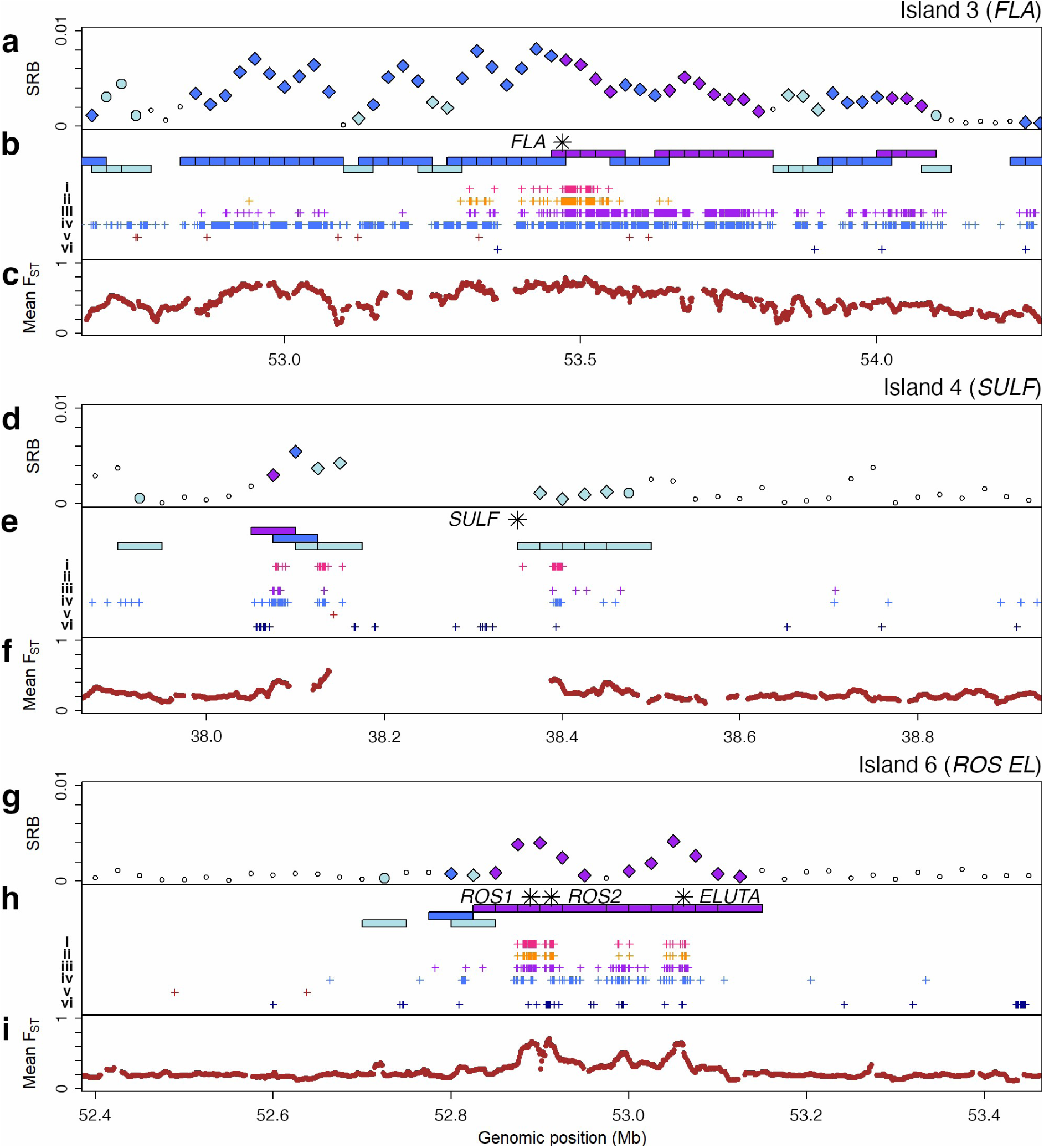
Windows and SNPs across islands 3, 4, and 6. **a, d, g** SRB for 50 kb window trees across three partition islands. Coloured diamonds indicate trees giving the A partition (purple), the A’ partition (dark blue), or the A’’ partition (light blue) in > 50% of bootstrap replicates (Fig. 3g). **b, e, h** 50 kb windows (rectangles, colour-coded as above) and SNPs below showing (i) The top 100 island SNPs showing the highest mean frequency difference between pseudomajus (A1) and striatum (A2) populations (magenta crosses) (ii) SNPs with mean allele frequency differences >= 0.8 between A1 and A2 populations (orange crosses). (iii) Partition-A SNP_XY_ trees (purple crosses). (iv) Partition-A’ SNP_XY_ trees (lighter blue crosses). (v) SNPs showing a mean read depth difference >= 0.95, depleted in pseudomajus. (vi) SNPs showing a mean read depth difference >= 95%, depleted in striatum. The locations of flower colour loci are indicated by asterisks. **c, f, i** Mean F_ST_-A, calculated from averages of π_w_, pairwise d_XY_ and pairwise π_t_, across all A1 and A2 populations, averaged in 10 kb windows with 9 kb overlaps.

We also used the equation for d_XY_ to determine trees for individual SNPs, termed SNP_XY_ trees. SNP_XY_ trees giving the A classification (purple crosses) were mainly upstream of *FLA*, while those giving the A’ classification (lighter blue crosses), were distributed throughout the island (the YP1 population is polymorphic for non-recombinant and recombinant *FLA* alleles). To display indels relative to the reference genome, we identified sites showing a difference in sequencing depth of >= 95% across a partition. Several sites were depleted in either pseudomajus (red crosses) or striatum (dark blue crosses). F_ST_-A was elevated throughout the 1 Mb *FLA* island (Fig. 8c).

For *SULF* (island 4), a single 50 kb A-partition window (purple diamond, Fig. 8d) was located about 300 kb to the left of the inverted duplication (indicated by asterisk) that generates the inhibitory small RNA. A’ and A’’ windows were distributed over a 0.5 Mb region, except for a 200 kb region to the left of *SULF* that is likely deleted in many striatum populations^18^. SNP and F_ST_ distributions reflected this pattern (Fig. 8e,f). For *ROS* and *EL* (island 6), A-partition windows and associated SNPs and F_ST_ peaks were focused in 100 kb regions around each locus (Fig. 8g-i).

Thus, the A-partition islands at previously identified colour loci were broadly centred on the loci and varied from 100 kb to 1 Mb, most likely reflecting variation in recombination rate. Islands 1, 2, 5, and 7 spanned 100 kb – 200 kb (Supplementary Fig. 21). Island 1 (*CRE*) showed a similar pattern around the candidate flower colour loci (Supplementary Fig. 21 a,b,c). Islands 2 (*AUN*), 5 (*RUB*), and 7 (*XAT*) lacked distinctive SRB or F_ST_ signals, but A-partition windows and SNPs were detected around each locus (Supplementary Fig. 21d-i).

### The partition B island may harbour a plant height gene

The B-partition island (island 8) was identified through two overlapping 50 kb windows (Fig. 9a-c). To determine whether a phenotype may be associated with this island, we analysed four populations sampled around a hybrid zone (see inset in Fig. 1). YP4 is B2, separated by the 10 km mountain pass from three B1 populations (YP1, MP4 and MP11) (Fig. 3f). Island 8 SNP frequencies showed shallow clines across the pass (Fig. 9e).

**Figure 9:**
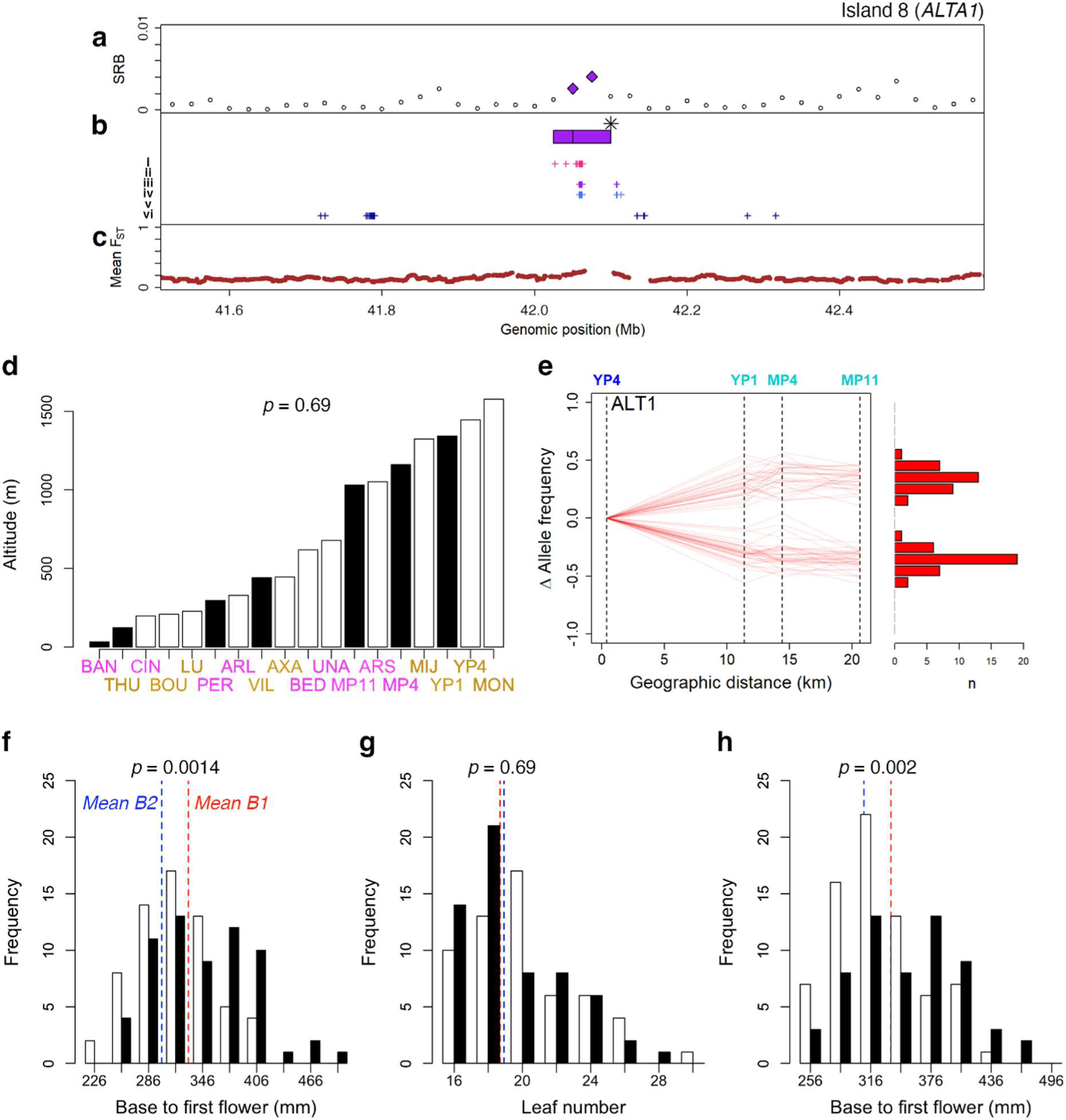
Windows, SNPs and altitudes for the B partition. **a** SRB for 50 kb window trees across island 8, with those giving partition B shown as purple diamonds. **b** Positions of 50 kb windows giving partition B with SNP distributions below. (i) The top 100 island SNPs showing the highest mean frequency difference between B1 and B2 populations (magenta crosses) (ii) SNPs with allele frequency difference >= 0.8 between B1 and B2. (iii) Partition B SNP_XY_ trees (purple crosses), (iv) Partition B’ SNP_XY_ trees (blue crosses). (v) SNPs with mean read depth difference > 95%, depleted in group B1. (vi) SNPsshowing a mean read depth difference > 95%, depleted in group B2. **c** F_ST_-B based on pairwise comparisons of all group B1 and group B2 populations, averaged in 10 kb windows with 9 kb overlaps. **d** Altitudes of populations. Black = B1, white = B2 and colour of labels indicates whether pseudomajus (magenta) or striatum (orange). p-value from a t-test for difference in altitude between B1 and B2 populations. **e** SNP clines as in Fig. 6 but for island 8 SNPs (B1 populations in cyan, B2 in blue). 100 SNPs were selected which showed the highest mean frequency difference between all 18 B1 and B2 populations. **f** Grouped histogram showing height from base to first flower of 128 plants homozygous for B1 (black) or B2 (white) alleles. X-axis values indicate the mean height of plants within each bin Vertical dashed lines show the mean height of B1 (red) and B2 (blue) homozygotes. p-values reflect a two-sample t-test between the heights of B1 and B2 plants. **g** As (f) but measuring leaf number rather than plant height. **h** As (f), but showing results for 133 plants homozygous for *ROS^p^* (black) and *ros^s^* (white) alleles.

YP4 is located at a higher altitude (1,446m) than the other three populations near the hybrid zone (1,343m, 1,161m and 1,031m, Fig. 9d). Plants sampled from altitudes at or greater than 1,350m tend to have reduced height and fewer nodes and branches when grown in a common environment^33^, raising the possibility that island 8 harbours a locus controlling one or more of these traits.

To test this possibility, we measured plant height and leaf number to the first flower for an F_2_ from a cross between YP4 (B2) and an *A. m. m.* var. *pseudomajus* accession sampled between MP4 and MP11 (Ventolà, B1) (Supplementary Table 1). Island 8 genotype showed significant linkage with plant height (t-test, *p* = 0.0014) (Fig. 9f). No significant difference between genotypes was found for leaf number (t-test, *p*-value = 0.69) (Fig. 9g), indicating that height differences were caused by differences in average internode length rather than number. As *ROS* was on the same chromosome as island 8 (Fig. 3g), we also genotyped the F_2_ population for this locus. *ROS* showed linkage with plant height though less significant than island 8 (t-test, *p* = 0.002) (Fig. 9h). Taking plants heterozygous for the island 8 marker, and ranking according to height, showed *ROS* had no significant effect (t-test, *p* = 0.078). By contrast, ranking *ROS* heterozygotes for height revealed that island 8 still had a significant effect on height (t-test, *p* = 0.033). Thus, island 8 harbours or is linked to a locus controlling plant height, hereafter named *ALTA1* (*ALT1*), with YP4 carrying a dominant allele conferring reduced height (heterozygotes were significantly shorter than B1 homozygotes (*p* = 0.046) but not significantly taller than B2 homozygotes (*p* = 0.077)). Island 8 contained a coding region with similarity to members of the cytochrome P450 family, which have previously been implicated in affecting plant height through regulation of brassinosteroid biosynthesis^34^ (Supplementary Table 5).

A previous study, in which seed sampled from natural populations was grown in a common environment in randomised blocks, reported that *A. m. m.* var. *striatum* (5 populations sampled) plants had significantly reduced height, node number and branch number than *A. m. m.* var. *pseudomajus* (8 populations)^33^. However, the two populations from the highest altitude were both striatum, raising the possibility that these differences were caused by differences in altitude. Rerunning their statistical model, including altitude and its interaction with variety, showed variety had no significant effect on plant height, node or branch numbers, while altitude and interaction terms did (Supplementary Table 7). Fitting a mixed model with population, family and block effects, showed that altitude was the only significant explanatory variable (Supplementary Table 8).

The previous study also found an altitudinal cline for plant height, node and branch number for *A. m. majus* var. *striatum* populations but not *A. m. majus* var. *pseudomajus*^33^. However, observation of a cline in striatum alone might again reflect sampling differences: the two highest altitude populations sampled (1,347m and 1,564m), were both striatum, with the highest elevation sampled for pseudomajus being 1,126m. Rerunning the same mixed model as above without the two highest altitude populations showed the effect of altitude became insignificant (Supplementary Table 9). Thus, the data can be most simply explained by reduced height, node and branch number for plants growing above 1,130m.

Our 18 sampled populations showed no significant correlation between populations classified as B1 or B2 and altitude (*p* = 0.69). However, the two populations at the highest altitude (MON at 1,578m and YP4 at 1,446m) were both B2 (Fig. 9d). Thus, the results are consistent with island 8 harbouring an *ALT1* allele conferring reduced height, favoured at altitudes greater than 1,350m, but with introgression of the allele into some lower altitude populations.

## Discussion

Blind genomic d_XY_ tree scans identify a single ancient multigene partition among the populations. The genomic islands giving this partition comprise less than 0.5% of the genome and harbour, or are linked to, eight loci controlling flower colour distributed over six chromosomes. Five of these loci (*SULF*, *FLA CRE, AUN* and *XAT*) control yellow, while three (*ROS*-*EL* and *RUB*) control magenta. Alleles at these loci interact to confer the complementary striatum and pseudomajus flower colour phenotypes. Except for *XAT,* which gives the weakest island signal, the loci exhibit steep clines in SNP frequencies across a hybrid zone between *A. m. m.* var. *pseudomajus* and *A. m. m.* var. *striatum*, indicating they harbour barrier loci^16,17,18,19,20,35^.

Interactions between the flower colour loci can be represented with a genotype-phenotype space (Fig. 10a). The pseudomajus genotype is at the top left, with alleles conferring spread magenta and restricted yellow, whereas the striatum genotype is bottom right, with alleles conferring spread of yellow, and restricted magenta. Hybrid genotypes, such as those conferring white or orange flower colours, occupy the remaining corner positions.

**Figure 10:**
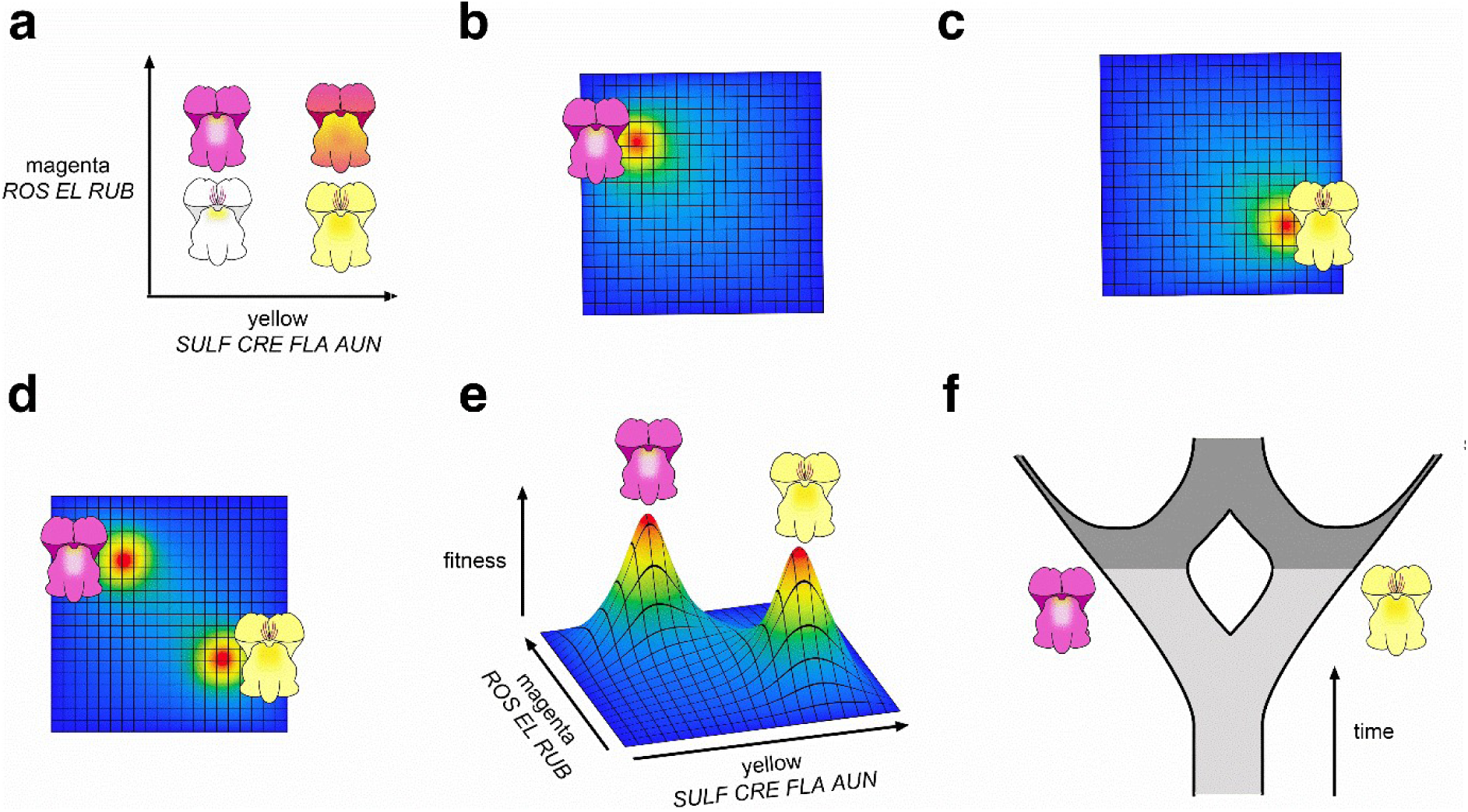
Genotype-phenotype and possible genotype-fitness spaces accounting for observed d_XY_ trees in *Antirrhinum*. **a** Genotype-phenotype space for flower colour. Magenta with a restricted yellow highlight (top left) corresponds to *A. m. m.* var. *pseudomajus*; whereas yellow with restricted magenta (bottom right) corresponds to *A. m. m.* var. *striatum*. **b, c** Genotype-fitness spaces for eco-adaptative barriers where one condition favours the *A. m. m.* var. *pseudomajus* genotype (b) and another favours the *A. m. m.* var. *striatum* genotype (c). Dark blue indicates lowest fitness, with fitness increasing up the colour gradient to a peak of red. **d, e** Genotype-fitness space viewed from the top (d) or side (e) where the phenotypes of *A. m. m.* var. *pseudomajus* and *A. m. m.* var. *striatum* occupy distinct adaptive peaks in a common environment, yielding hybrid incompatibility. **f** Hypothesis for origin of observed d_XY_ trees. In the absence of gene flow, genomes in different populations undergo divergence (increase in d_XY_). The merging path following secondary contact reflects extensive gene flow, with the outer paths representing the fraction of the genome that resists gene flow.

One hypothesis for how flower colour genes act as barrier loci is that they are eco-adaptive. In environmental conditions on one side of the partition, where *A. m. m.* var. *pseudomajus* is found, the top left position in genotype-phenotype space has the highest fitness (Fig. 10b); whereas on the other side of the partition, where *A. m. m.* var. *striatum* is found, the fitness peak lies at the bottom right (Fig. 10c). Divergent selection on the flower colour loci would then prevent flow of alleles between the varieties.

Several observations argue against this hypothesis as being the sole explanation for the partition. First, the hybrid zone is only 1 km wide, with no obvious difference in environment or pollinators^36^ between the two sides. This situation contrasts with other plant systems in which partitions correspond to different niches (e.g. different pollinators) and operate over a broader geographic scale^37,38,39,40,41,42^. Second, if there is an ecological difference across the partition, we would expect loci affecting traits unlinked to flower colour to contribute to the partition islands. For example, in the niche-differentiated systems cited above, as well as flower colour, traits such as flower shape and nectar composition are affected by loci distributed throughout the genome. Yet all the striatum-pseudomajus partition islands harbour or are linked to genes controlling flower colour. Loci controlling other traits could be located within or near the islands, though there is no evidence for inversions creating supergenes^43^ (the largest island, harbouring *FLA*, shows the same low recombination rates when both varieties are backcrossed to a common parent^19^). Third, if ecology is the sole driver, we would expect other multigene partitions to be observed among the 18 populations, given the wide range of ecological conditions they occupy^44^, from coastal to montane habitats. Yet we only observe a single ancient multigene partition.

A more likely hypothesis is that rather than having a single peak, the fitness landscape has two peaks within the same environment (Fig. 10d and e). Hybrid genotypes fall into a fitness valley leading to hybrid incompatibility. Such a fitness landscape is plausible because a yellow highlight on magenta, or a magenta highlight on yellow, could both be effective for signalling pollinator entry. Each signal requires coadaptation of several components (the different genes controlling flower colour), allowing more than one fitness peak to be produced through reciprocal sign epistasis^45^. Hybrid genotypes may be less attractive because overlapping yellow and magenta reduces colour contrast (e.g. orange flower at top right of Fig. 10a), and/or because a white background is less salient to pollinators in the ecological context of striatum and pseudomajus (bottom left, Fig. 10a). The two-peak hypothesis does not rule out eco-adaptive effects: the relative heights of the two peaks may vary according to environment.

According to the twin peak hypothesis, divergence between alleles in the two varieties may be accounted for by an ancestral population split (Fig. 10f). The depth of d_XY_ tree division at the flower colour loci is 2-3 x 10^-3^. Assuming a mutation rate of 10^-8^ per base pair per generation, the ancestral split may have occurred of the order of 10^5^ generations ago, consistent with previous estimates^15^. In the absence of gene flow, genomes in different populations would have undergone divergence (increase in d_XY_) after the split. Following secondary contact (dark grey, Fig. 10f), perhaps at the end of the last ice age 10^4^ years ago, gene flow would erode genome divergence except for regions near hybrid-incompatible loci. Such loci may arise infrequently on a microevolutionary timescale as they depend on populations arriving at equivalent but hybrid-incompatible adaptive solutions. Colour patterns with a signalling role may be particularly prone to this route, as illustrated by the many equivalent aposematic butterfly wing patterns^46^, accounting for why the only multigene partition to be found among the *Antirrhinum* populations corresponds to this trait.

### A candidate eco-adaptive barrier locus

In addition to the multigene 9:9 partition, we found a single island that gives a 7:11 partition. SNPs at this locus exhibit shallow clines across the mountain pass lying between a high and lower altitude striatum populations. The island is linked to a gene controlling plant height, *ALT1*, with the allele in the high-altitude population giving reduced height, consistent with seed harvested from such populations giving shorter plants^33^. Thus, *ALT1* may be an eco-adaptive locus, with alleles conferring reduced height selected for at high altitude.

A difficulty with this hypothesis is the lack of correlation between the 7:11 partition and altitude. Although the partition divides two highest populations, both striatum, from the two lowest, both pseudomajus, intermediate populations are not partitioned according to altitude. Possibly an *ALT1* allele conferring reduced height in ancestral high altitude striatum populations introgressed into lower altitude populations through compensatory allele frequency changes at other height loci. Compensation is plausible because unlike the epistatic interactions of flower colour loci, genes controlling plant height are largely additive and exhibit considerable variation within natural populations^47,48^. Alternatively, introgression could reflect selective pressures other than altitude, such as the extent of vegetation coverage^49^ and the presence or absence of shade^50^, which select for reduced height in some low-altitude populations.

### Limitations and strengths of blind tree-scanning

A limitation of our method is that it depends on barrier loci diverging sufficiently that they give a detectable d_XY_ increment. Recently arisen barrier loci, or loci in very highly recombining regions, would therefore evade detection.

A strength is that blind tree scanning may have greater resolving power than F_ST_ scans for identifying candidate barrier loci. Several of the islands we identify (islands 2, 5, 7 and 8) do not exhibit F_ST_ peaks when the relevant populations are compared. Some of the additional islands we identify could be spurious. However, all four exhibit clines based on DNA sequence pools^20^, and are linked to barrier-relevant traits (flower colour for islands 2, 5 and 7; plant height for island 8). Island 5 is detected using a tree approach based on F_ST_ instead of d_XY_, indicating that the greater resolving power arises, at least in part, through screening for tree structure, which may capture weaker groupings than screening for F_ST_ peaks.

Perhaps the most important advantage of blind tree scanning is that can identify ancient barriers without bias from prior classifications. Relationships between genomic partitions and those based on observed phenotypes can then be evaluated, providing insights into how barriers across the genome, under different evolutionary forces, manifest themselves and contribute to speciation.

## Methods

### DNA extraction from pooled leaf tissue for whole genome sequencing

Leaves were sampled from 20-60 individuals from each population and pooled within populations (Supplementary Table 1). DNA was extracted from each leaf pool, sequenced, and mapped to the *Antirrhinum majus* reference genome^21^. SNPs were filtered to ensure all were biallelic.

Genomic DNA was isolated using a cetyltrimethylammonium bromide (CTAB) method on 2-5g of leaves harvested as final weight for pooled samples as described^51^. Field samples were either placed in bags of silica or water-moist paper towel and kept cool at 4°C until they were courier posted by overnight delivery to the John Innes Centre and, upon arrival, frozen at - 80°C. Greenhouse samples were stored in foil at −80°C. A GPS reading was taken at each wild population sampling location (recorded in Supplementary Table 1). Short read sequencing (2 x 150bp) of DNA extracted from leaf tissue was carried out as described^52^. Continuous long read (CLR) sequencing on the PacBio platform was also carried out for *A. m. m.* var. *pseudomajus* sample Z-NDM as described^52^.

### RNA extraction from petals

Total RNA was isolated from petal tissues using the Qiagen RNeasy Plant Mini Kit, including DNase I treatment. For a comparison of *A. m. m.* var. *pseudomajus* (accession Ac1266) and *A. m. m.* var. *striatum* (accession Ac1125) we harvested petal lobes from dissected flowers just before opening. Three independent samples were used for reproducibility, and each sample was a pool of 3 individuals, each contributing 1-2 flowers. Samples were sequenced by Novogene using an Illumina HiSeq 2500.

### Growth conditions and crosses

Greenhouse cultivated accessions were grown in JI compost soil mixes as described^53^, with supplemental lights in winter giving 12 to 16 hour days. Plants were grown outside in the summer using the same soil mixes, in pots or plugs in trays placed upon raised benches. Plants stocks were maintained through inter-sibling crosses. To carry out crosses, pre-anthesis floral buds were opened with forceps and young anthers were removed. Once the flower was open (within two to four days) pollen from another plant was applied to the stigma, using the forceps to carry pollen or the whole stamen. Seed capsules were harvested after four to six weeks. Panels of hybrid progeny were genotyped across partition island loci to ensure correct transmission of parental genotypes.

### Flower photography

Flowers were placed on a black velvet background with a scale bar and a Small Grey & Colour Separation Chart (Danes Picta BST13) for monitoring light level, colour and white balance. We used an Olympus XZ-1 (10 Megapixels) Camera, with side-on lighting from table lamps fitted with halogen 42W 630 lumen (2800k) warm white light bulbs. Camera settings were set to the closest White Balance of 3000K, with no flash, an Exposure Time of 1/20-1/40 sec, F stop 8.0, ISO 200, RAW images, Macro On, aspect 4:3, high definition.

### Measuring plant height and leaf number

Plant height was determined by measuring the distance from the base of the plant (where the stem meets the soil) to the position where the flower (pedicel & bract) meet the stem. A string was used to get tight access to the stem and follow any bends along its length. The string was then measured against a ruler.

Leaf number was determined by counting up all the leaves (or petioles for any fallen leaves) appearing at nodes along the stem, starting from the first leaves and ending at the last leaf below the first flower/bract. The early leaves (vegetative phase) are usually in pairs at each node, before single leaves at the transition to the inflorescence.

### KASP genotyping

KASP Genotyping was performed as described^18^. Fluorescent signals were detected using a BioRad CFX96 light cycler, and data processed using BioRad CFX Manager v3.1. AFLP methods used standard PCR and agarose gel analysis as described^19^. KASP / AFLP oligos for islands 1-6 are as described^19^. Island 7 and 8 oligos are recorded in Supplementary Table 10.

### Assembly of *A. m. m.* var. *pseudomajus* and *A. m. m.* var. striatum genomes

*A. m. m.* var. *pseudomajus* accession Z-NDM and *A. m. m.* var. *striatum* accession Z-VCO were grown from seed, and leaf tissue was harvested and sequenced. Raw Illumina reads were trimmed using Trim Galore! [https://github.com/FelixKrueger/TrimGalore] with parameters -q 25 and --stringency 3 to remove low-quality sequence and adapters. To infer genome properties prior to assembly, GenomeScope2^54^ was used to evaluate heterozygosity rate and haploid genome length based on the k-mer spectrum. Contig-level assembly of PacBio CLR reads was carried out using the Canu package (version 1.9)^55^. Redundant heterozygous contigs in the assembly were then removed using purge_dups^56^. After identifying and masking repetitive sequence in the genome using RepeatMasker^57^, gene prediction was conducted using BRAKER2 pipeline^58^ by integrating plant proteins retrieved from SwissProt^59^ and RNAseq data aligned to the genome assembly. The overall quality of the assembly was assessed based on the LAI (LTR Assembly Index)^60^, while the completeness of gene annotation was evaluated by BUSCO^61^.

### Mapping reads to reference genome and generating SYNC files

FASTQ reads for each of the sequenced pools were mapped to the *A. majus* reference genome V3.0 (Genome Warehouse accession number GWHBJVT00000000) and to the *A. m. m.* var. *pseudomajus* genome using BWA-MEM, with the `-M` flag set for downstream compatibility with Picard (http://broadinstitute.github.io/picard/). Output SAM files were sorted using SAMtools, and Picard MarkDuplicates was used to remove duplicate reads. Local realignment of reads around indels was carried out using GATK version 3.5.0^62^. GATK RealignerTargetCreator was used to generate a list of intervals for realignment using GATK IndelRealigner. Processed BAM files were combined to generate MPILEUP files using SAMtools mpileup^63^. The minimum mapping quality threshold (`-q`) was set to 40, with a minimum base quality threshold (`-Q`) of 30. Orphaned reads were included in variant calling by using the `-A` flag. The `-B` flag was used to disable probabilistic realignment in base alignment quality calculations, as this can increase misalignments and false SNP calls. One MPILEUP file was generate for each of the eight *A. majus* chromosomes. MPILEUP files were converted to the Popoolation2 SYNC format using mpileup2sync.jar, which offers a more concise representation of allelic depth across populations^64^.

### Population genomic analyses

Within-population diversity, π_w_, was estimated as

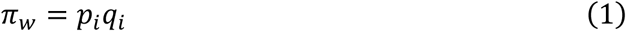

where p_i_ and q_i_ refer to the frequencies of alleles p and q within a single population, population i. d_XY_, between-population diversity, was estimated using

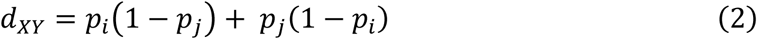

where i and j refer to a pair of populations, i and j. We also calculate 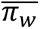 (mean *π_w_* between i and j). F_ST_, relative divergence, is estimated as

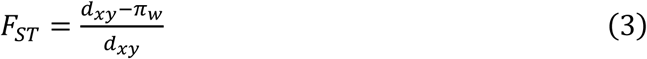

Nei’s standard genetic distance was calculated as

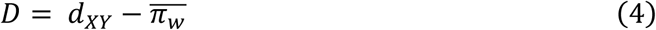

### treeXY population analyses

Population genomic statistics were calculated using the treeXY software version 1.1. Statistics were summarised in windows – unless otherwise stated, analyses detailed here used a 50 kb window size, with 25 kb overlaps between adjacent windows. treeXY was run on all SYNC files, yielding windows for all eight chromosomes. Window size was set to 50 kb using `-w`, and window overlap to 25 kb using `-o`. To output treeXY-filtered SYNC files, `--write_sync` was set, and `--compute_trees` was used to enable computation of SNP trees. `--ignore_multiallelic` was set such that multiallelic sites would be skipped. SNPs were required to be present in all 18 samples (`-A 18`). Otherwise, default parameters were used (minimum depth = 15, maximum depth = 200, minimum allele depth = 2).

### Constructing mean d_XY_ and D trees

treeXY outputs CSV files containing population genetic statistics summarised in windows of user specified size, with the option of additional single site statistics. These tables were read using the *generate_dXY_trees.R* companion script. Distance matrices were populated with between-population statistics calculated for each genomic window. UPGMA hierarchical clustering trees were generated for each distance matrix using the base R *hclust* function, with clustering method set to average. To generate mean trees across forests, the mean was taken across all windows using the *Reduce* function.

### Whole genome sweep analysis

A whole genome selective sweep was simulated within the one population, MP11, by fixing all polymorphic sites for one allele, randomly sampled based on the existing allele frequencies. To do this, treeXY was run with default settings, with the `--write_sync` flag enabled to output a depth-filtered SYNC file. A custom Python script, *artificial_sweep.py*, was run with the filtered SYNC file as input. At all genomic sites where more than one allele showed depth >= 2 (the default allelic depth threshold,) one allele was randomly sampled. Frequencies of ambiguous (N) alleles were ignored. The probability of sampling an allele was weighted according to its frequency prior to sweeping, with common alleles more likely to be sampled than rare ones. After sweeping, the depth of the chosen allele was changed to equal the sum of the total site depth, and all other allele frequencies were set to 0. The modified SYNC file, with the swept MP11 population, was written to a new file for downstream analyses.

### Whole genome trees with d_XY_, D, and maximum likelihood

For d_XY_ and D trees, a modified version of the treeXY script was run, which calculated the mean for both genetic distance measures across all biallelic sites and between all pairs of populations. Results from all eight SYNC files were averaged, and used to populate distance matrices. These matrices were then summarised as UPGMA trees using the base R *hclust* function. For maximum likelihood (ML) trees, input SYNC files were prepared which included the *A. sempervirens* Pont Napoléon population (Supplementary Table 1) along with the 18 *A. m. m.* var. *pseudomajus* and *A. m. m.* var. *striatum* pools. Filtered SYNC files were obtained by running treeXY on these SYNC files with the `--write_sync` option enabled. Filtered SYNC files were concatenated into a whole genome biallelic SYNC file. At each site, and in each population, a consensus base call was made by selecting the allele with the highest frequency. This yielded a set of aligned consensus sequences, which was converted into FASTA format. To generate an ML tree from this alignment, IQ-TREE^65^ version 2.3.2 was used, with the arguments `-m MFP -allnni -B 10000`, and the sempervirens outgroup specified using `-o`. ML trees were rooted and drawn in RStudio, using the *phangorn* library. d_XY_, D, and ML trees were generated before and after applying the whole genome selective sweep within the MP11 population.

### Calculating shortest root branch and cophenetic correlation coefficient

The shortest root branch (SRB) of a given tree was calculated from its cophenetic matrix, obtained using the base R cophenetic function. SRB is equal to the maximum value within the cophenetic matrix, subtract the second highest value. The similarity of hierarchical clustering trees was estimated using the cophenetic correlation coefficient, *c*^23^. To compare two trees, the Pearson correlation coefficient was calculated between their cophenetic matrices, where:

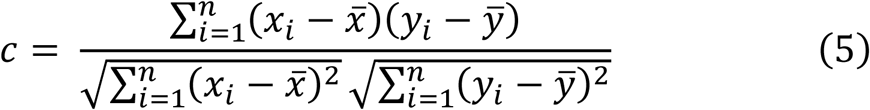

*x* and *y* correspond to the distances in each matrix. When carrying out the grouping tree scan to populate forests, a *c* threshold of 0.5 was chosen to capture trees showing moderate or high topological similarity. To test the replicability of grouping tree scan results, a bootstrapping approach was implemented. Each bootstrap replicate consisted of one grouping tree scan. For each scan, all forests were summarised based on the mean SRB of their trees. The forest showing the highest mean SRB was deemed the outlier forest, and recorded. Once all bootstrap replicates had been completed, all outlier forest regions were recorded, along with the frequency with which they had been detected.

### Classifying trees based on root division

Root division was determined using the base R cutree function, with *k* = 2, which bisects trees at their topmost branch to yield the two outermost clades. To classify trees based on root division, populations within each clade were recorded, and compared to a user specified signature.

### Identification of SNPs and Indels from SYNC files

SNPs were identified using the custom Python script *snp_tables.py*. A site was deemed to contain a SNP if its read depth met the minimum threshold of 10 (*-d 10*) in all populations (*-n 18*), and two alleles were present in at least two populations (*-a 2*). Indels were identified using *indel_tables.py*. Indels were included if they met the minimum read depth threshold of 5 (*-d 5*) in at least 2 populations (*-a 2*).

### Identification of top 100 SNPs, based on mean frequency difference

SNP tables, output by the *snp_tables.py* script, were read using the *split_SNP_tables.R* script. For each SNP table, mean frequency difference was calculated as the absolute difference between the sum of allele 1 frequency between A1 and A2, or B1 and B2, pools.

### Generation of island summary plots

The complete pipeline for generating island summary plots is documented within the *island_summary_pipeline.R* script. Requisite data are generated by running treeXY, as well as the companion python scripts *snp_tables.py* and *indel_tables.py*. In short, treeXY was run twice, to generate data for 50 kb windows, 10 kb windows, and d_XY_ SNP trees. Indels and SNPs were called from the same set of input SYNC files, using island coordinates to limit the searches to regions of interest (specified using *-m* and *-M*). Multi-panel plots were generated using *island_summary_pipeline.R*, and its associated R scripts.

### Geographic cline analysis

Genotyping of plants across the Planoles hybrid zone and clinal analysis of the *CRE* island, as shown in Fig. 5j, was carried out as described^20^. Pool-seq cline analysis was carried out using the custom R script *densiCline.R* and a panel of four populations spanning the Planoles hybrid zone (YP4, YP1, MP4, and MP11). For each island, 100 SNPs were chosen based on difference in allele 1 frequency (SNPs which were detected in the 18 population data, but not the hybrid zone data, were ignored). SNP frequency in each pool was plotted minus the frequency in YP4, in order to visualise the relative frequency change.

### Colour ranking of flower photographs

Flower photographs were processed in Photoshop. Flower photographs included a Danes Picta BST13 colour chart, which was selected with the eyedropper tool to standardise white balance. Adjusted images were saved in the JPEG format. Images were cropped to retain only the flowers, before being loaded into a Photoshop canvas. Each image was, in turn, visually appraised for the intensity of its yellow or magenta colour, and positioned within the rank accordingly. Positions were adjusted as more photographs were incorporated, until a final ranking was reached. Upon completion, the ranking was split into either two halves, or four quartiles. Within each half/quartile, genotype data for *CRE*, *AUN*, or *XAT* were checked, and the proportion of each marker allele was inferred.

### RNAseq differential expression analysis using DESeq2

Ribo-depleted total RNA libraries from *A. m. m.* var. *pseudomajus* M-AUT and *A. m. m.* var. *striatum* Z-ALE were mapped to the *A. m. m.* var. *pseudomajus* assembly using HISAT2^66^. Output SAM files were sorted using Samtools, and transcripts were assembled using StringTie^67^, operating in expression estimation mode (`-e`) and reporting gene abundances (`-A`). To prepare data tables for analysis, read counts were extracted across all samples using the StringTie prepDE.py Python script. Differential expression analysis was carried out using DESeq2 version 1.32.0^24^ using default parameters.

## Acknowledgements

We thank the John Innes Centre (JIC) Horticultural Services for providing growth facilities and maintenance of plant material, and JIC Research Computing for use of High Performance Computing facilities. We also thank Tingting Li for carrying out a replicate flower colour ranking analysis and assisting with photography.

This work was supported by the Biotechnology and Biological Sciences Research Council (grants BB/S009256/1, BB/G009325/1, BBS/E/JI/230002C, and BBS/E/J/000PR9773 to EC, and Norwich Research Park Biosciences Doctoral Training Partnership grant BB/M011216/1 to DR), the National Natural Science Foundation of China (grant 32030007, to YX), and an ANR funded French Laboratory of Excellence project (‘LabEx TULIP’, to MB). This research was also funded in whole or in part by the Austrian Science Fund (FWF) [P 32166] (to DF). HS was funded by the EU’s Horizon 2020 research and innovation programme under the Marie Skłodowska-Curie Grant Agreement No.101034413.

## Data availability

Raw DNA and RNA data, along with the *A. m. m.* var. *pseudomajus* and *A. sempervirens* genome assemblies have been uploaded to NCBI SRA / WGS under accession number PRJNA1232105. The *A. majus* reference genome V3.0 is available at the NGDC Genome Warehouse under accession number GWHBJVT00000000.

## Code availability

The treeXY software and documentation are available at https://github.com/DR-Antirrhinum/treeXY. Other scripts used to carry out the analyses presented here are available at https://github.com/DR-Antirrhinum/phylogenetic-forests.

## Author contributions

DR designed and carried out bioinformatic analyses of DNA and RNA, developed research software, managed data, and redrafted the manuscript with EC. DB carried out genotyping, DNA / RNA extraction for sequencing, and flower photography, and assisted with flower colour ranking and manuscript discussion. LC designed and carried out crosses to provide experimental plant populations. AW designed RNA and DNA mapping pipelines, and contributed to manuscript discussions. MB and CA together undertook fieldwork to identify and collect tissue from wild plant populations. SZ carried out the assembly of the *A. m. m.* var. *pseudomajus* genome, helped with data transfer and organisation. HS carried out the reanalysis of the common garden statistical data from Marin *et al.* (2020). DF carried out cline analysis of the CRE locus, and contributed to project supervision and manuscript discussions. YX administered and supervised genome sequencing, assembly and data transfer. EC conceived and administered the project, wrote and redrafted the manuscript with DR, designed experiments and contributed to methodology.

## Additional information

Supplementary Information is available for this paper.

Correspondence and requests for materials should be addressed to Enrico Coen, Yongbiao Xue, or David Field

