## Supplementary Data for "Blind genomic tree scans identify loci underlying adaptive peaks in Antirrhinum"

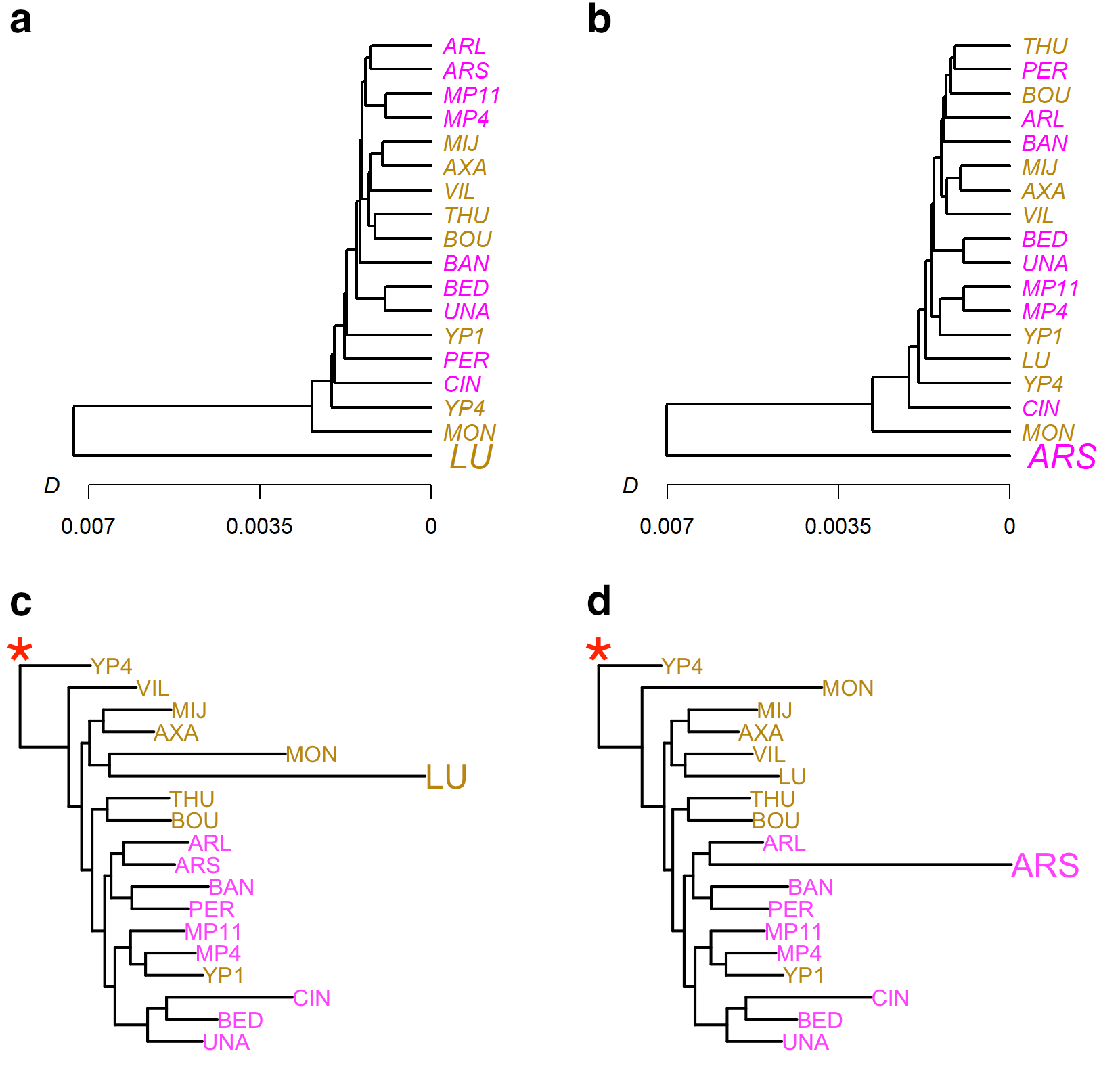

### **Supplementary Figure 1: Effect of population selective sweeps on maximum likelihood trees**

**a** Whole genome *D* tree after simulating a selective sweep across all polymorphic sites in the LU population. **b** Whole genome *D* tree after simulating a selective sweep across all polymorphic sites in the ARS population. **c** Maximum likelihood whole-genome tree, rooted on an *Antirrhinum sempervirens* outgroup, based on consensus sequences of genomic SNPs after simulating a selective sweep in the LU population. Red asterisk indicates the root, based on outgroup *A. sempervirens* (not shown). **d** Maximum likelihood tree generated after simulating a selective sweep in the ARS population, rooted on *A. sempervirens*. Magenta labels indicate *A. m. m.* var. *pseudomajus* populations, and yellow labels show *A. m. m.* var. *striatum* populations.

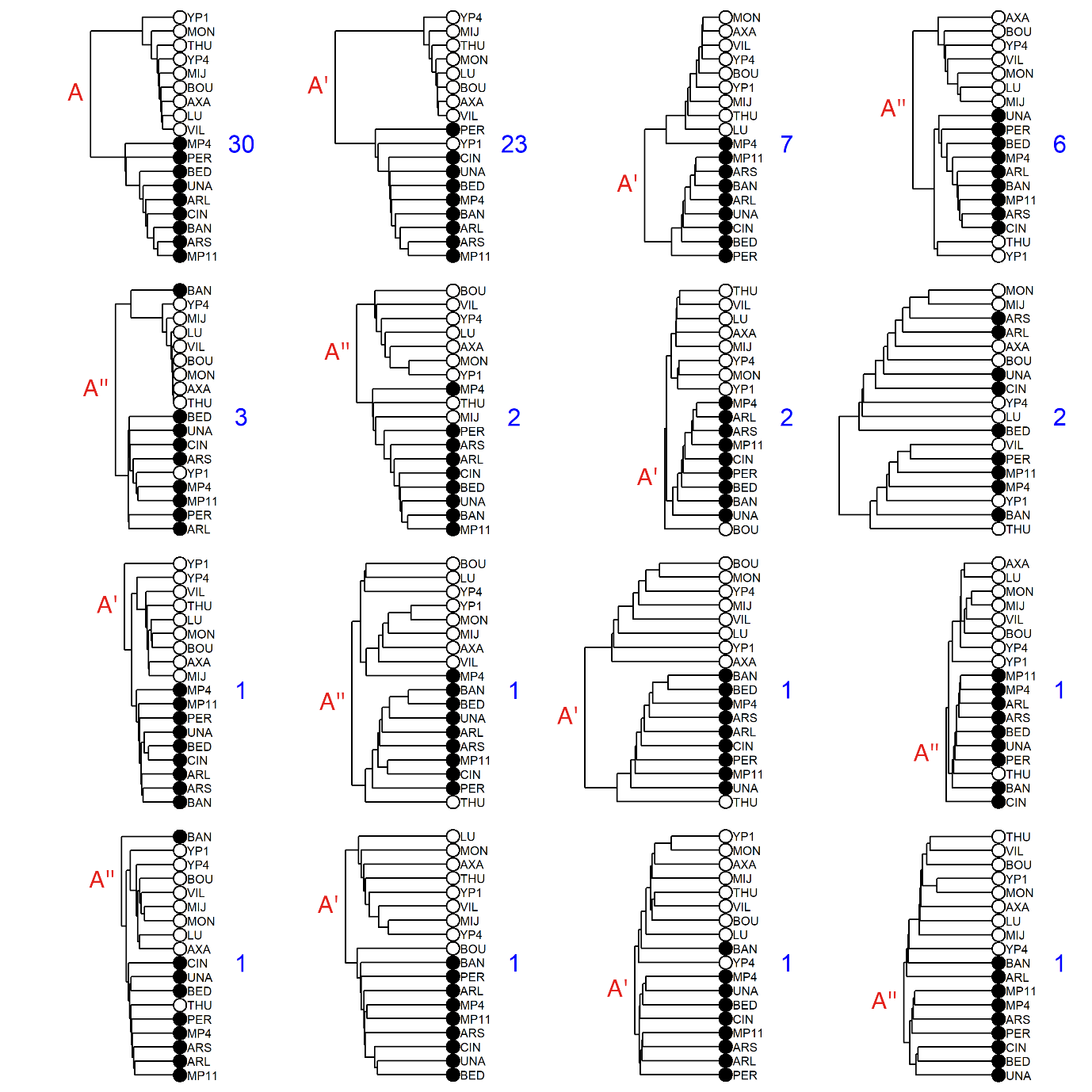

### **Supplementary Figure 2: Diversity of tree partitions present within an outlier forest**

Representative trees showing all partitions appearing within the turquoise outlier forest shown in Fig. 3b. For each partition, the tree with the highest SRB is shown. Number of trees within each partition is indicated. Black circle indicates A1 group, white indicates A2. Partitions are labelled A, A’ (one population misgrouped compared to A), or A’’ (two populations misgrouped compared to A) in red, at the root of each tree (one partition, corresponding to two trees, does not correspond to any of these classifications).

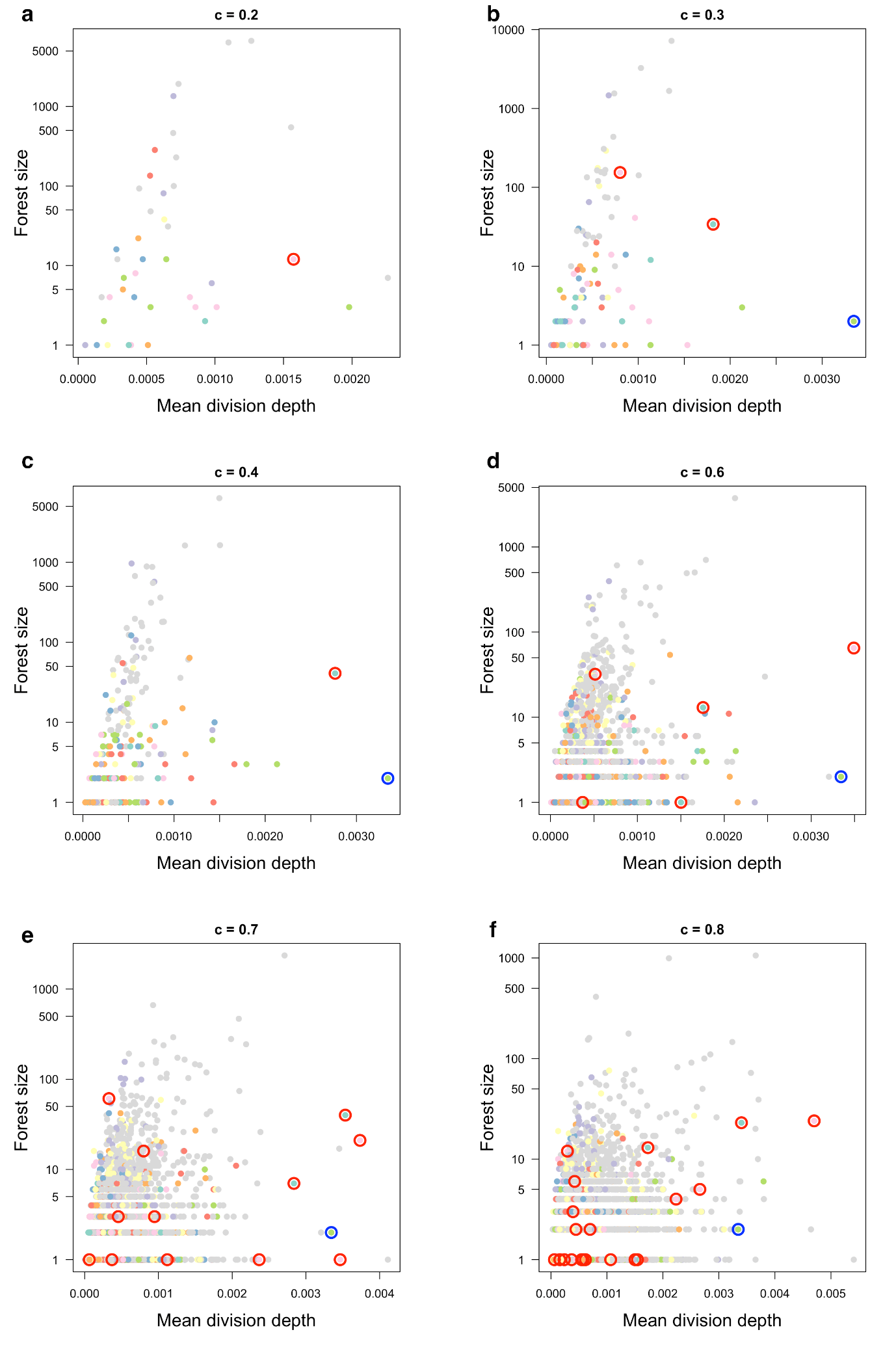

### **Supplementary Figure 3: Results from individual grouping tree scans using different *c* thresholds**

**a-f** Forests from genomic d_XY_ tree scans using varying cophenetic correlation coefficient (*c*) cutoffs. Forest colour reflects the partitions shown in main text Fig 3a. Forests showing partition A / A’ topologies (red) and the partition B topology (blue) are circled. The same seed tree was used for each c-value.

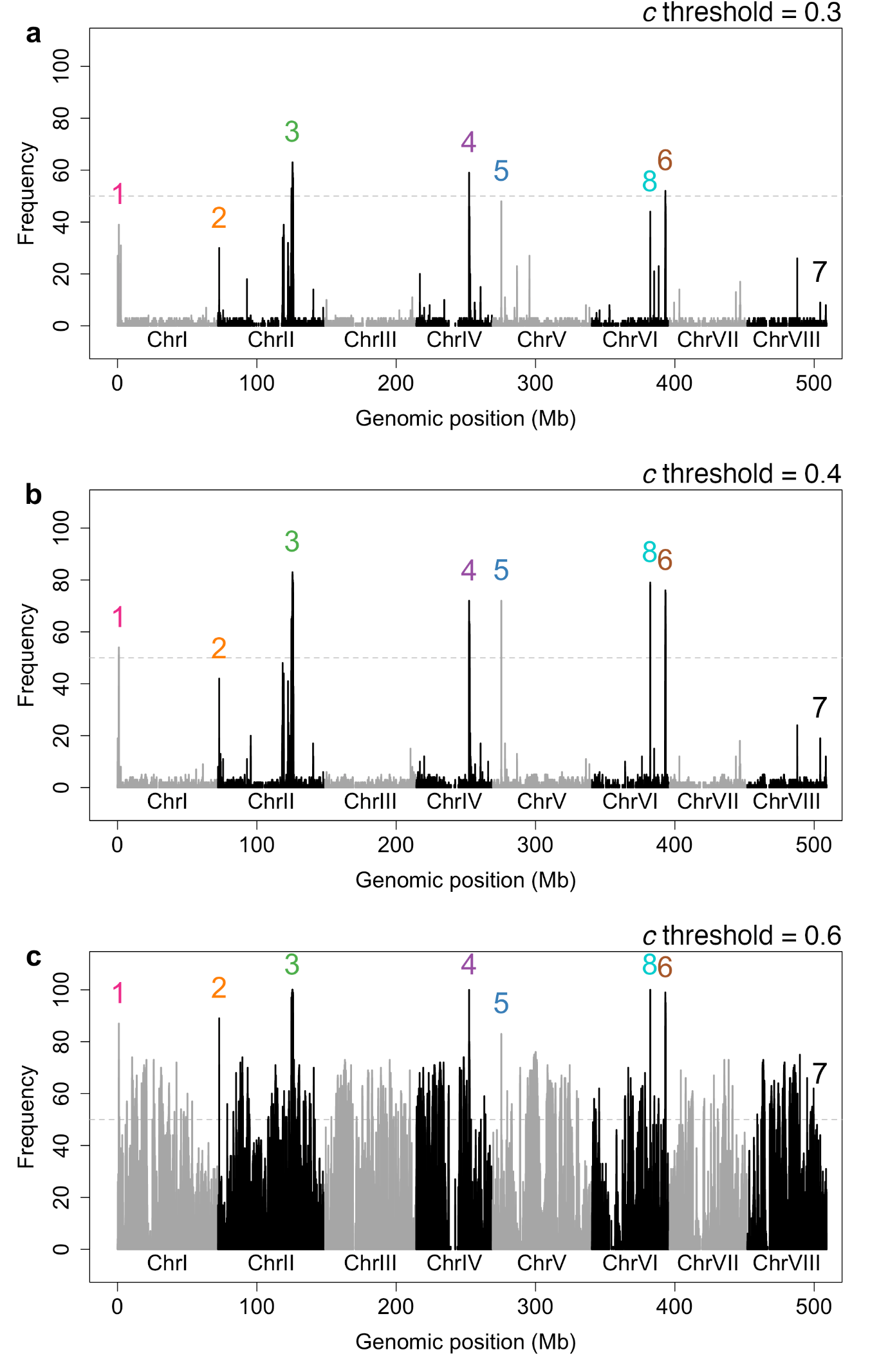

### **Supplementary Figure 4: Grouping tree scan bootstrap analyses with different *c* thresholds**

Frequency with which different genomic regions occurred within outlying forests (mean SRB >= 0.002), across 100 tree-scan bootstrap replicates, using varied cophenetic correlation coefficient (*c*) thresholds. Numbers 1-6 show the locations of the partition islands identified in Main Text Figure 3. Chromosome indicated by alternating grey-black.

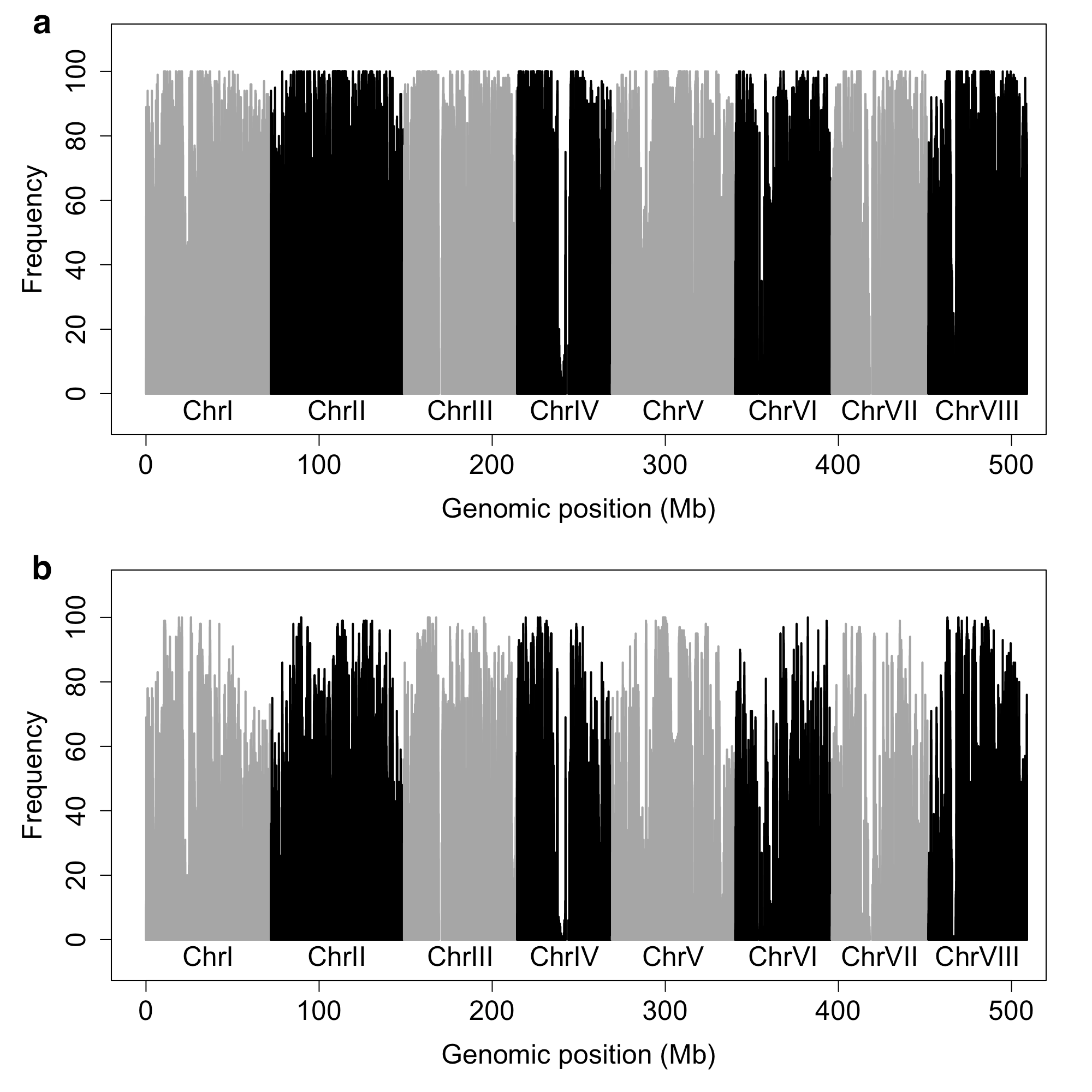

### **Supplementary Figure 5: Grouping tree scan bootstrap analyses using varied *SRB* thresholds, and fixed seed trees**

**a** Frequency with which different genomic regions occurred within outlying forests (mean SRB >= 0.001), across 100 tree-scan bootstrap replicates (*c* threshold = 0.5). Chromosome indicated by alternating grey-black. **b** As (a), but with mean SRB >= 0.0015**.**

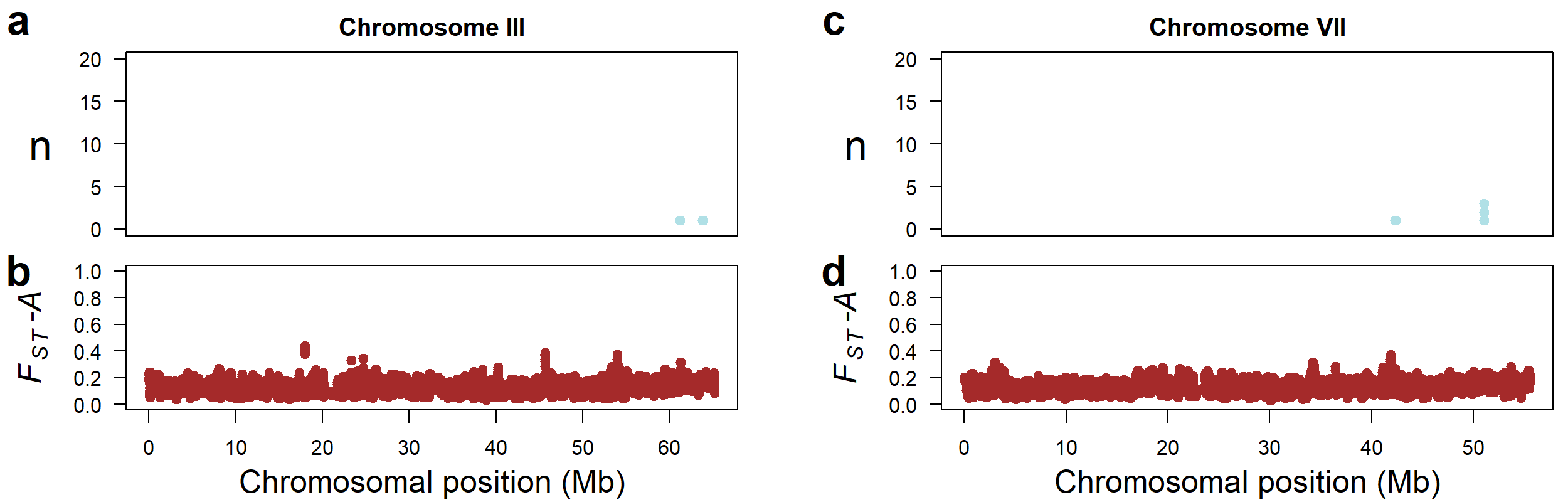

### **Supplementary Figure 6: Chromosome scans for monophyletic density and F_ST_ on chromosomes not harbouring partition islands**

**a, c** Frequencies of 5 kb windows showing the A partition (purple), the A’ partition (one population misgrouped compared to the A partition, dark blue), or the A’’ partition (two populations misgrouped compared to the A partition, light blue) across chromosomes III and VII, summed over 50 kb windows with 25 kb overlaps, mapped to the *A. majus* reference genome. **b, d** Mean F_ST_ for pairwise comparisons of all Group A1 and Group A2 populations, averaged in 10 kb windows with 9 kb overlaps.

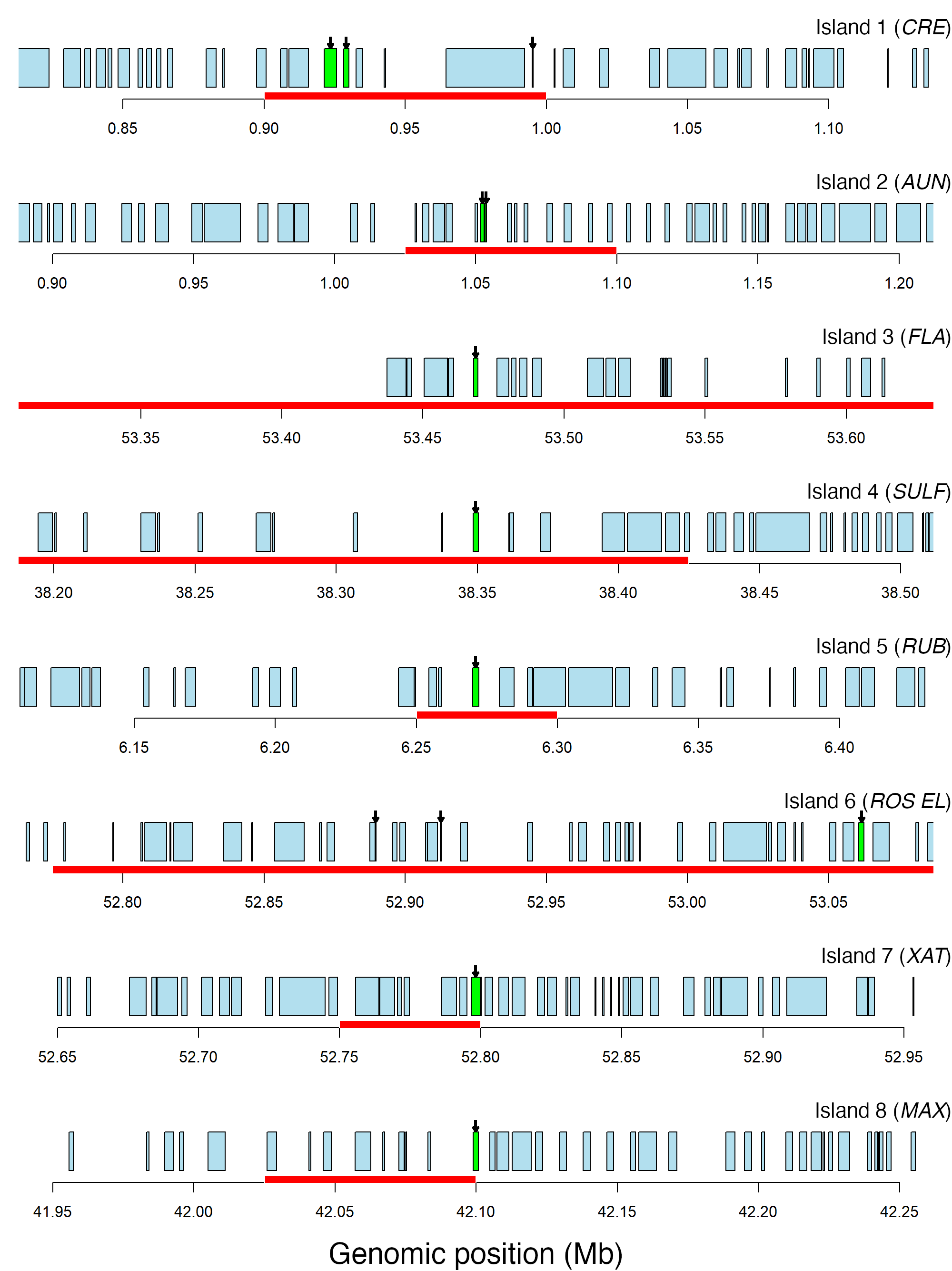

### **Supplementary Figure 7: Coding sequences within partition islands**

**a-h** Boxes representing the positions of predicted coding sequences within 300 kb sections centred on candidate loci within partition islands 1-8. Green, arrowed boxes represent candidate colour loci from prior studies (*FLA*, *SULF*, *ROS EL*), coding sequence prediction (Island 8) or differential expression analysis (main text Figure 8). Island boundaries are represented by a red rectangle (*FLA*, *SULF*, and *ROS EL* exceed the 300 kb size limit).

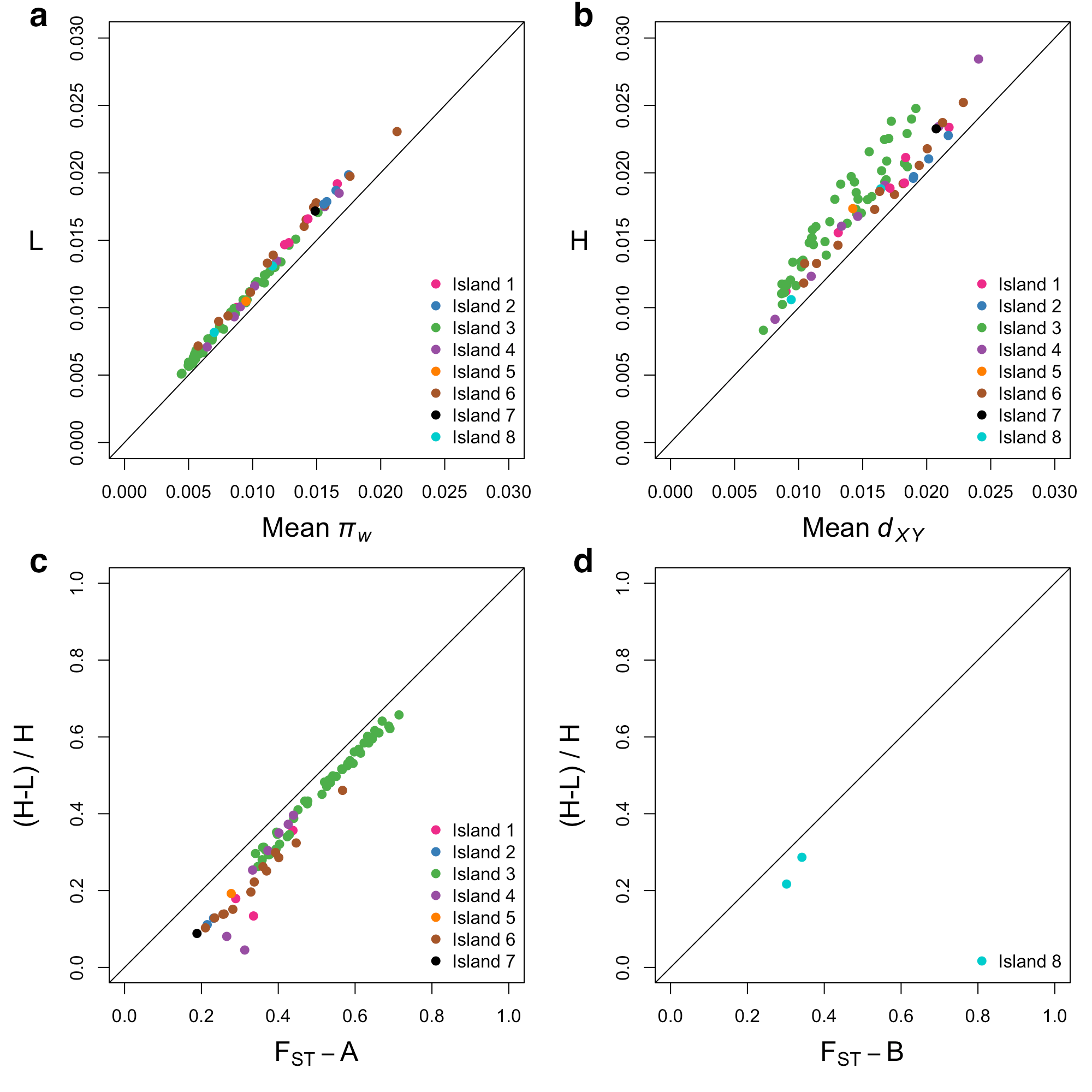

### **Supplementary Figure 8: Comparison of d_XY_ tree measurements to mean d_XY_, π_w_, and F_ST_ for partition island windows**

**a** Mean terminal branch length (L) plotted against mean π_w_ for 50 kb windows overlapping partition islands. **b** Tree height (H) plotted against mean d_xy_ for 50 kb windows overlapping partition islands. **c** Tree height (H) minus mean terminal branch length (L) divided by H compared to F_ST_-A for 50 kb windows overlapping partition islands. Points are coloured according to island membership. **d** H minus L divided by H compared to F_ST_-B for island 8 windows.

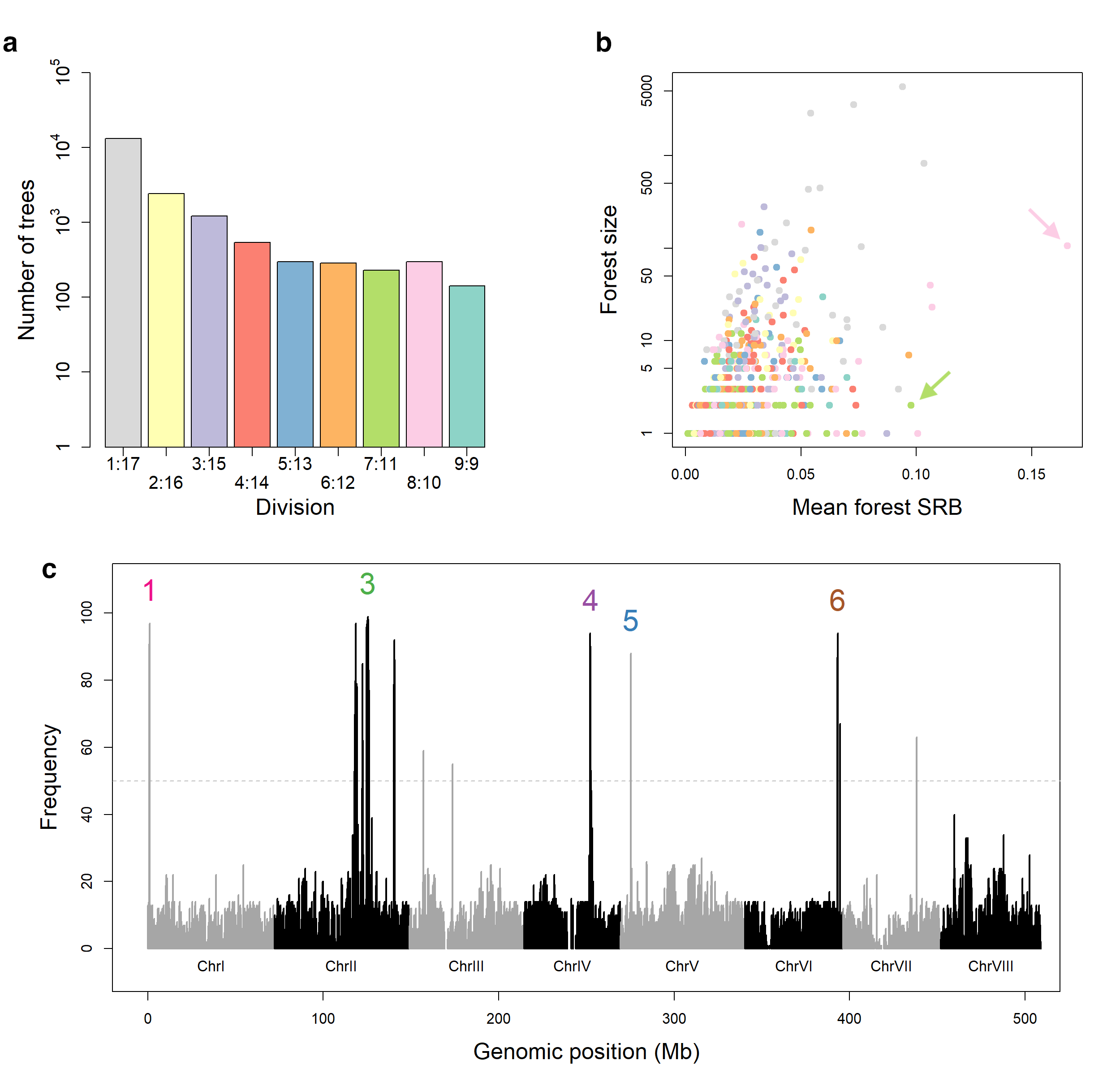

### **Supplementary Figure 9: Grouping tree scan results using F_ST_**

**a** Histogram showing the number of 50 kb window F_ST_ trees corresponding to each partition. **b** Forests from a genomic F_ST_ tree scan, with the number of trees within each forest (forest size) plotted against the mean shortest root branch (SRB). Forest colour reflects the partitions shown in Supplementary Fig 9a. The partition A outlier forest (pink) and partition B forest (green) are arrowed. **c** Frequency with which different genomic regions occurred within outlying forests (mean SRB >= 0.08), across 100 tree-scan bootstrap replicates. Numbers 1, 3, 4, 5, and 6 show the locations of the partition A islands with bootstrap values >= 50. Chromosome indicated by alternating grey-black.

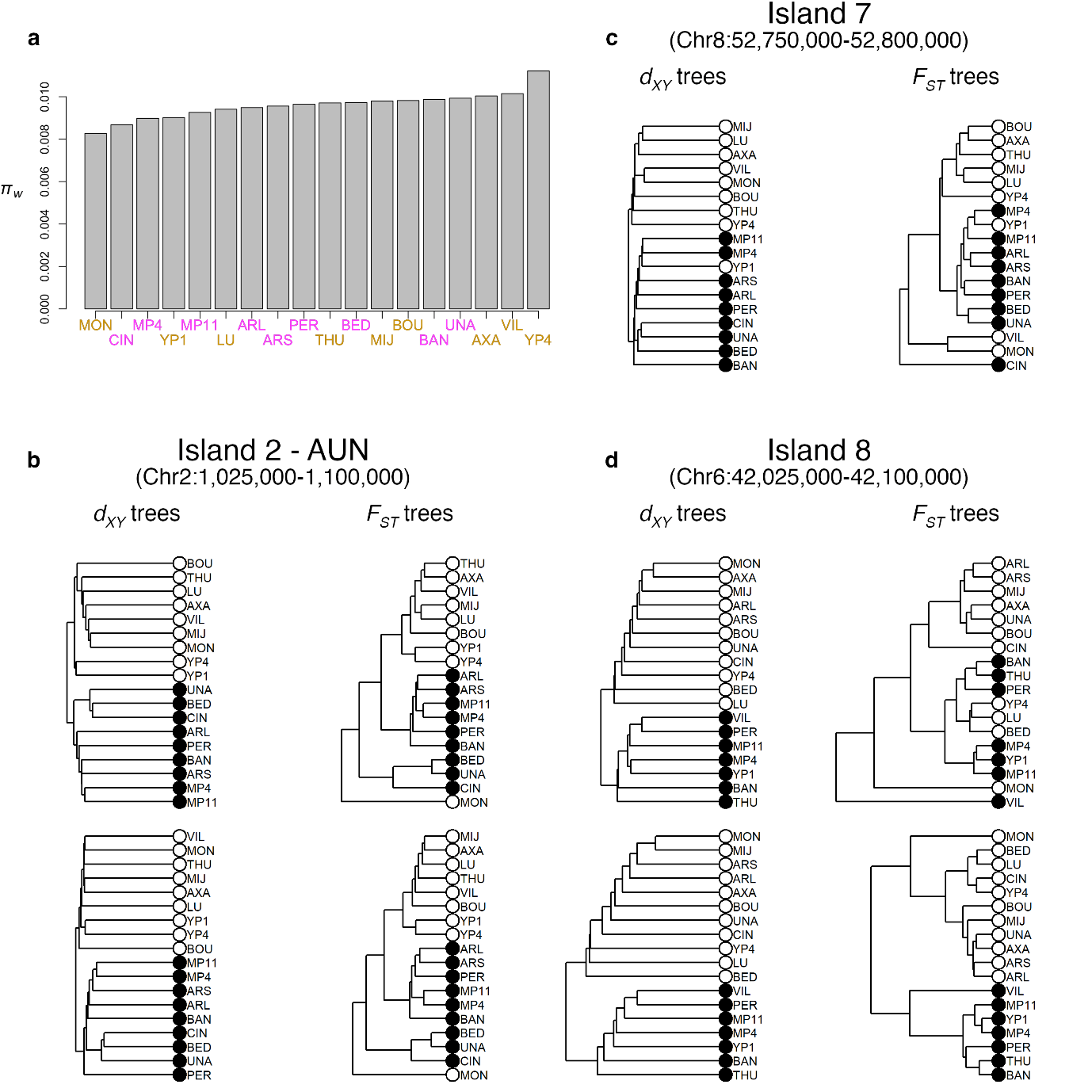

### **Supplementary Figure 10: π_w_, and F_ST_ compared to d_XY_ trees**

**a** Whole genome mean π_w_ for each of the 18 pools of *A. m. m.* var. *pseudomajus* (magenta labels) and *A. m. m.* var. *striatum* (yellow labels). **b, c** d_XY_ and F_ST_ trees from the 50 kb windows comprising partition islands 2 and 7. Black circles denote A1 populations, and white circles A2 populations. **d** d_XY_ and F_ST_ trees from partition island 8, the B partition island. Black circles denote B1 populations, and white circles B2 populations.

*ROS ^p^ / ROS ^p^ el ^p^ / el ^p^ SULF ^p^ / -*

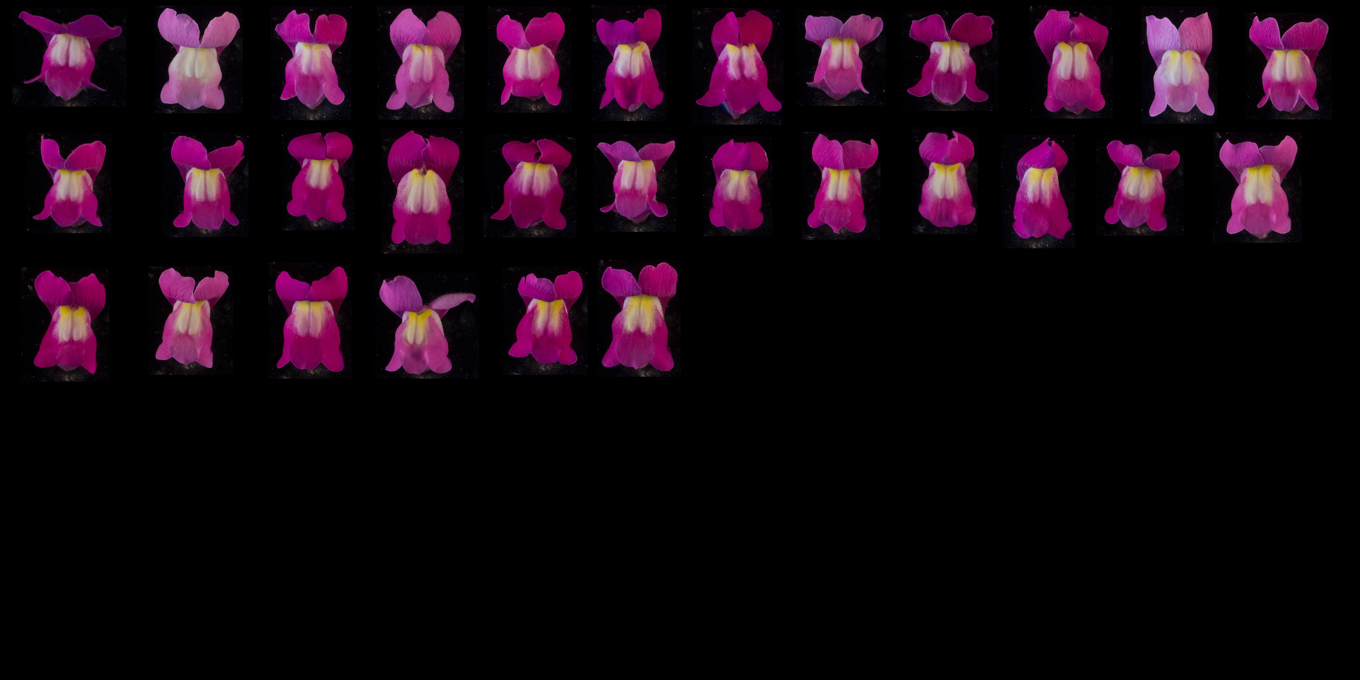

*ROS ^p^ / ROS ^p^ el ^p^ / el ^p^ sulf ^s^ / sulf ^s^*

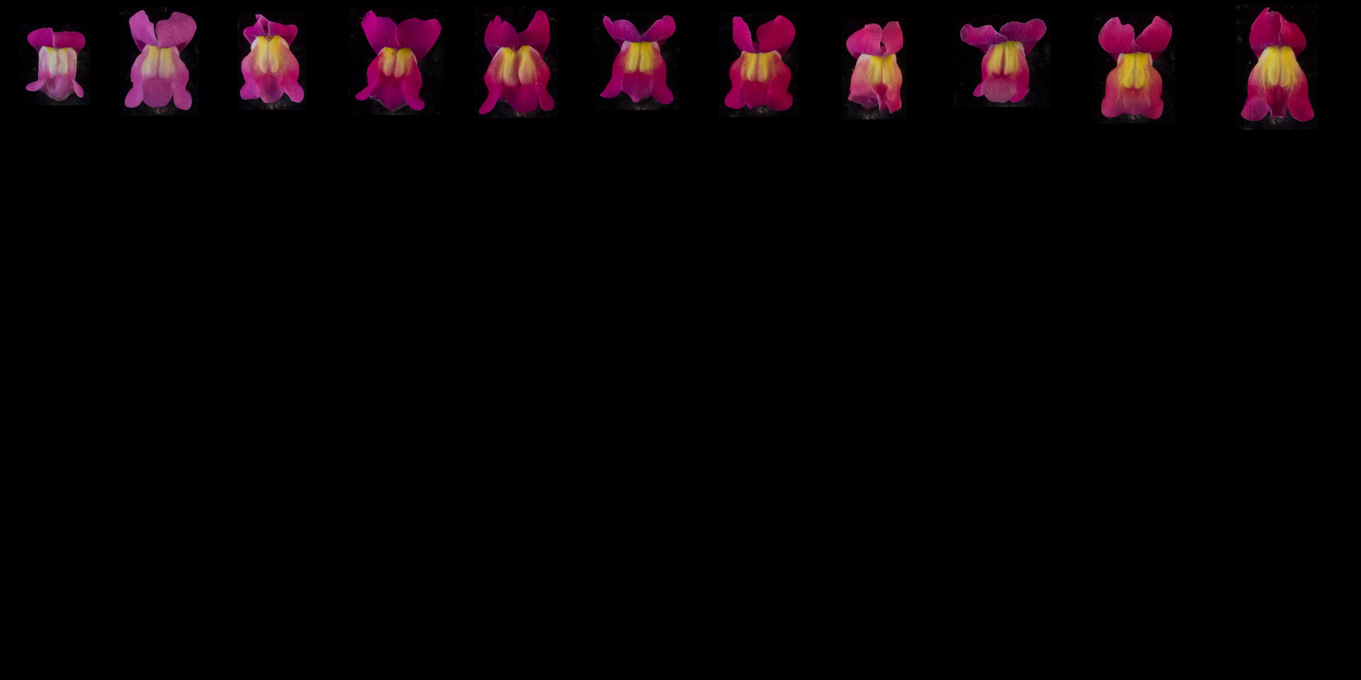

*ROS^p^ / ros^s^ EL^s^ / el^p^ SULF ^p^ / -*

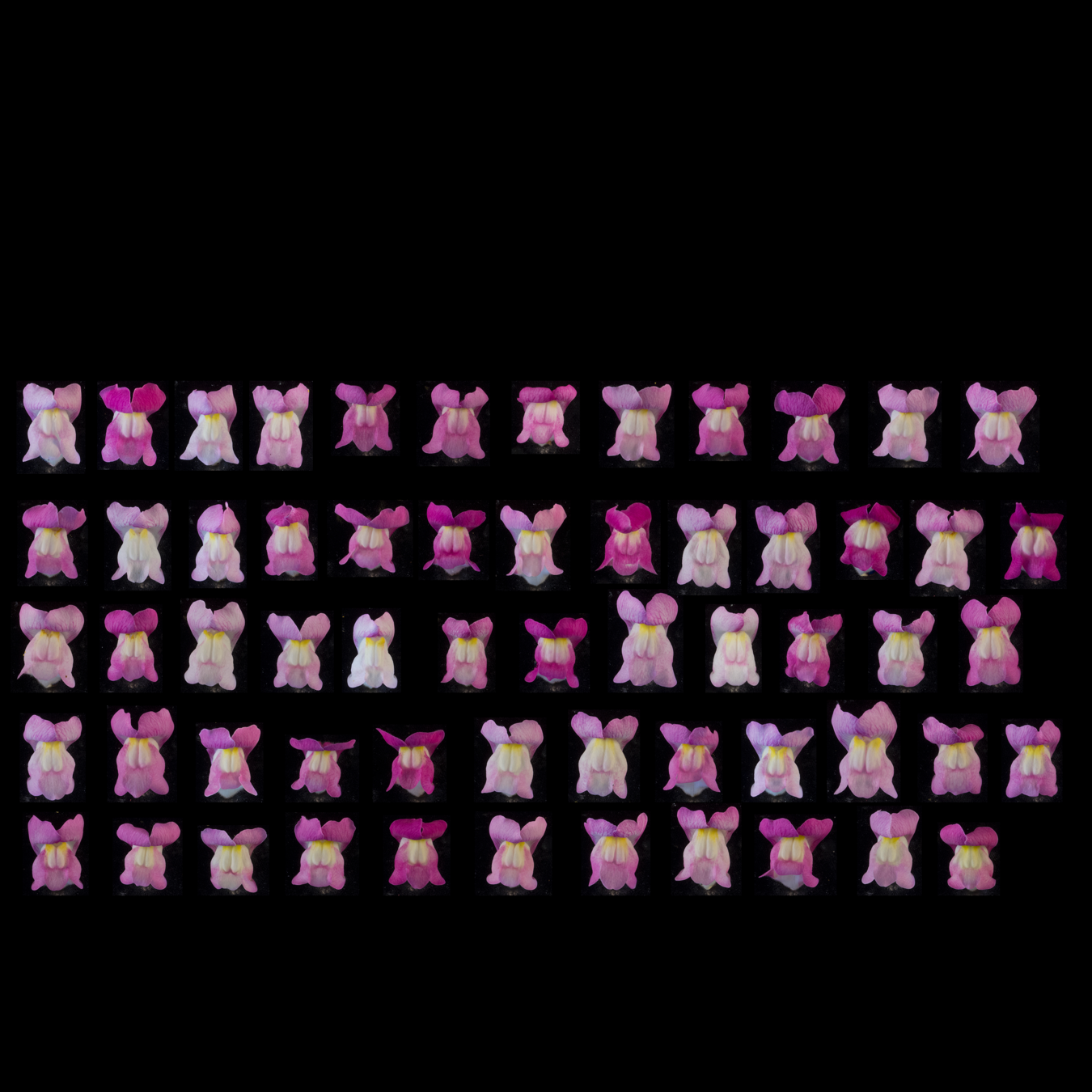

*ROS ^p^ / ros ^s^ EL^s^ / el ^p^ sulf ^s^ / sulf ^s^*

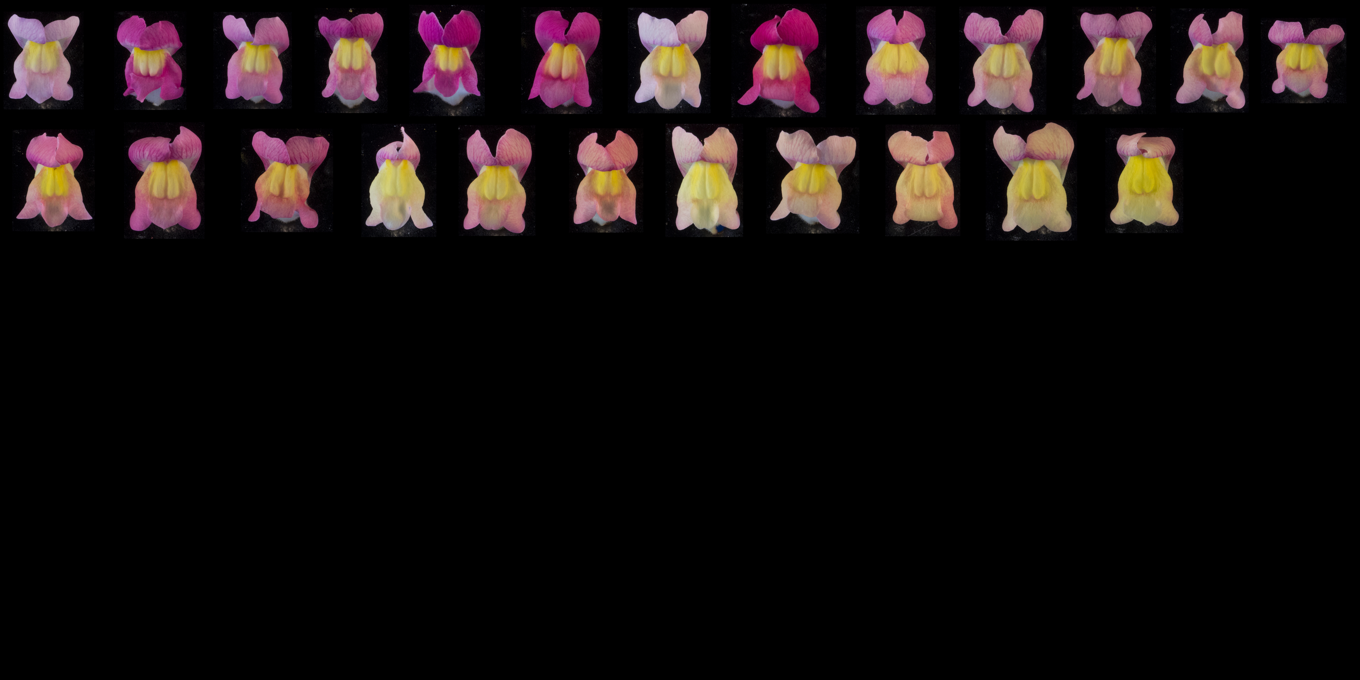

*ros^s^ / ros^s^ EL^s^ / EL^s^ SULF ^p^ / -*

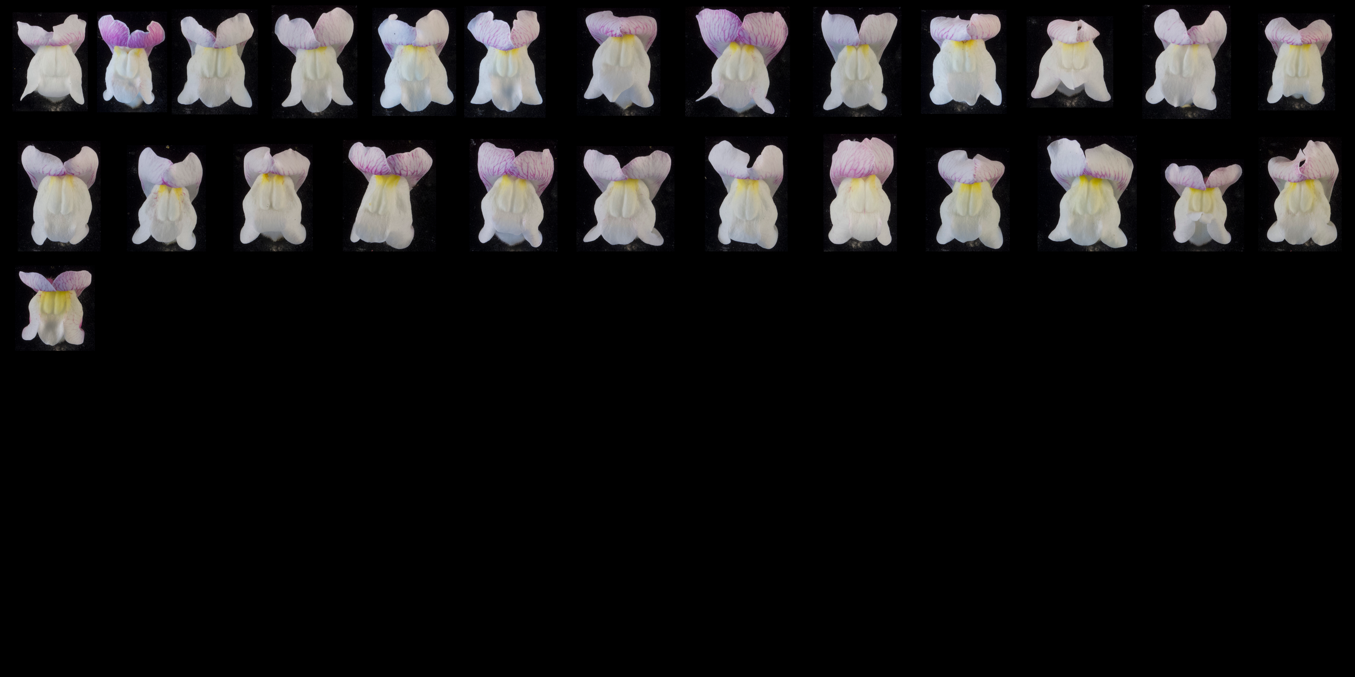

*ros^s^ / ros^s^ EL^s^ / EL^s^ sulf^s^ / sulf^s^*

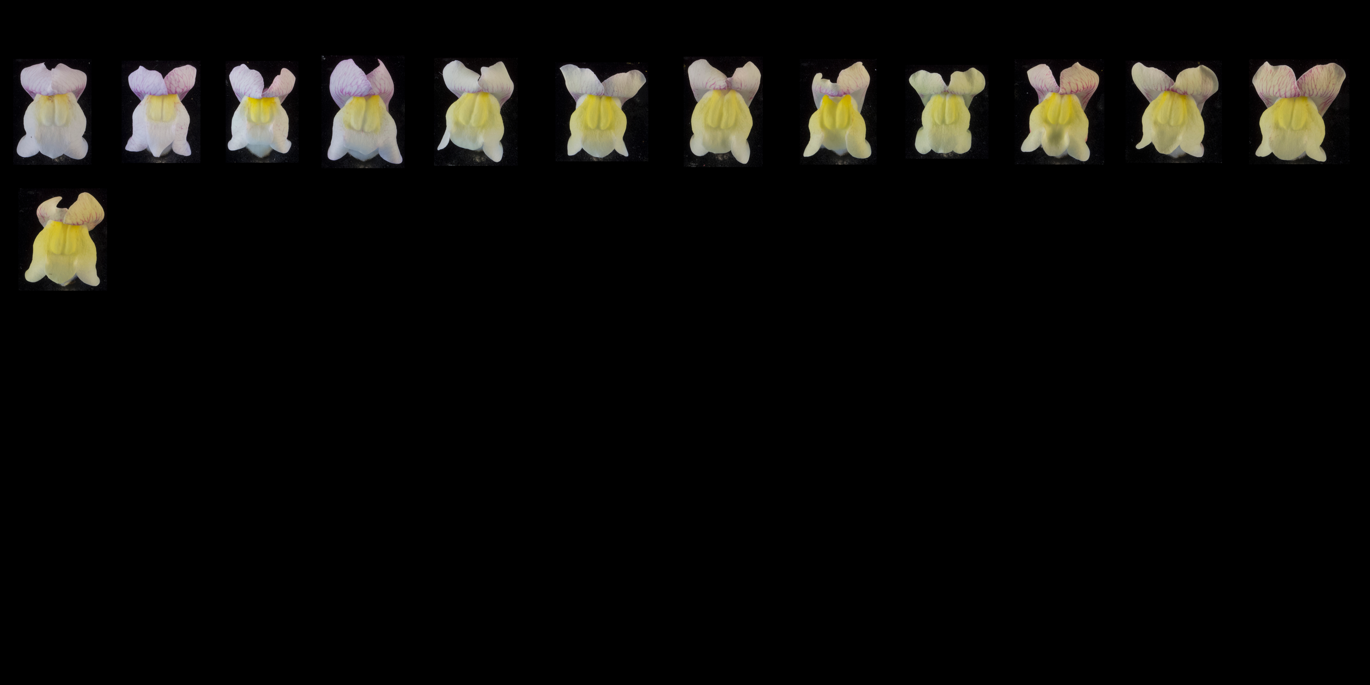

### **Supplementary Figure 11: Yellow colour rankings of F_2_ family J109, grouped by genotype**

Flowers ranked by extent of yellow: family J109 (F_2_ of *A. m. m.* var. *pseudomajus* x *A. m. m.* var. *striatum*). Individual accessions were grouped according to *ROS, EL*, and *SULF* genotype, before being ranked for the extent of yellow pigmentation. The flower deemed to show the most restricted yellow was placed in the top left position, with the amount of yellow increasing along each row from top left to bottom right. Rankings were made without prior knowledge of island 1 or island 2 SNP genotypes. Superscript ‘s’ within the genotype denotes the *A. m. m.* var. *striatum* allele, and superscript ‘p’ denotes the *A. m. m.* var. *pseudomajus* allele.

*ROS ^p^ / ROS ^p^ el ^p^ / el ^p^ SULF ^p^ / -*

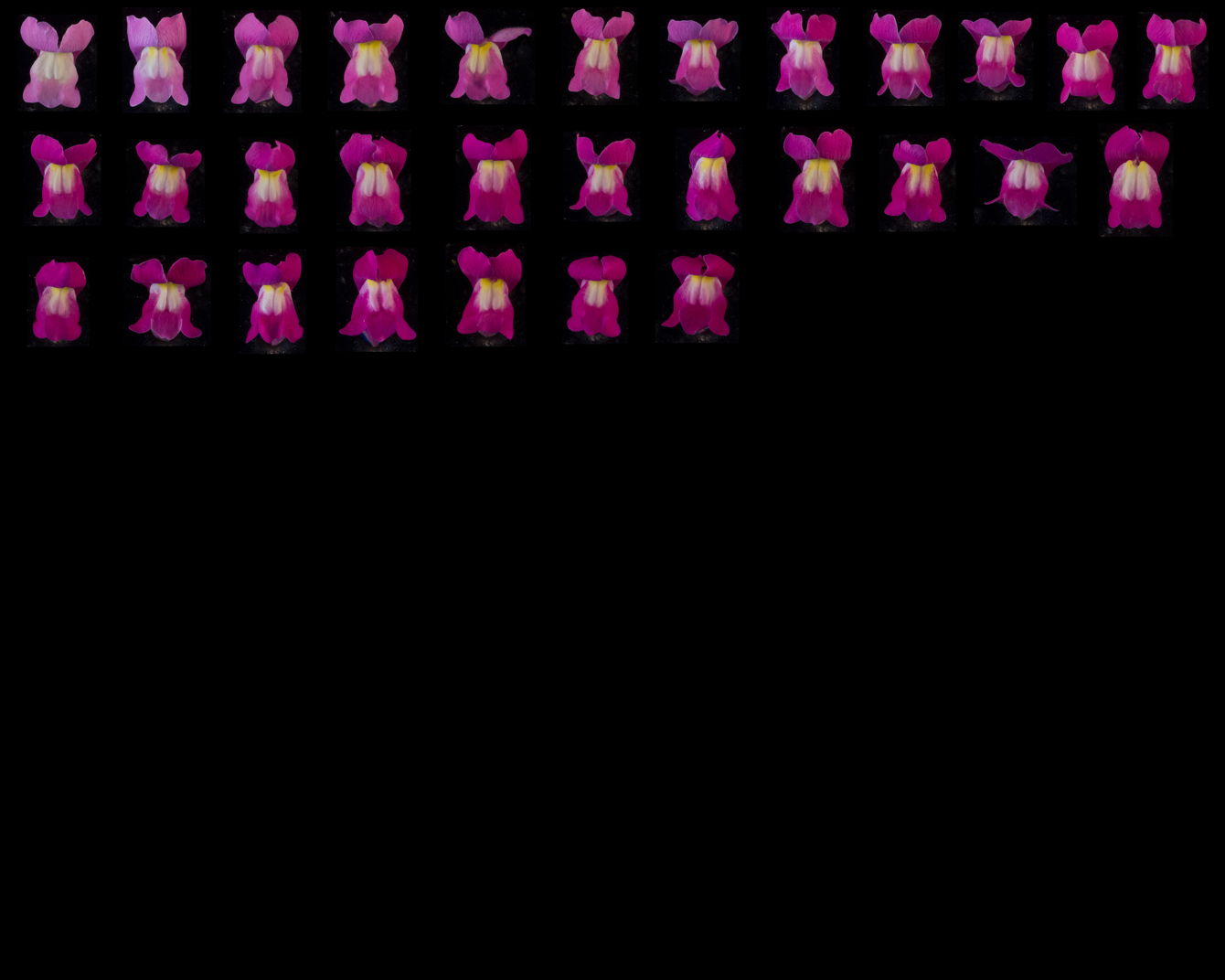

*ROS ^p^ / ROS ^p^ el ^p^ / el ^p^ sulf ^s^ / sulf ^s^*

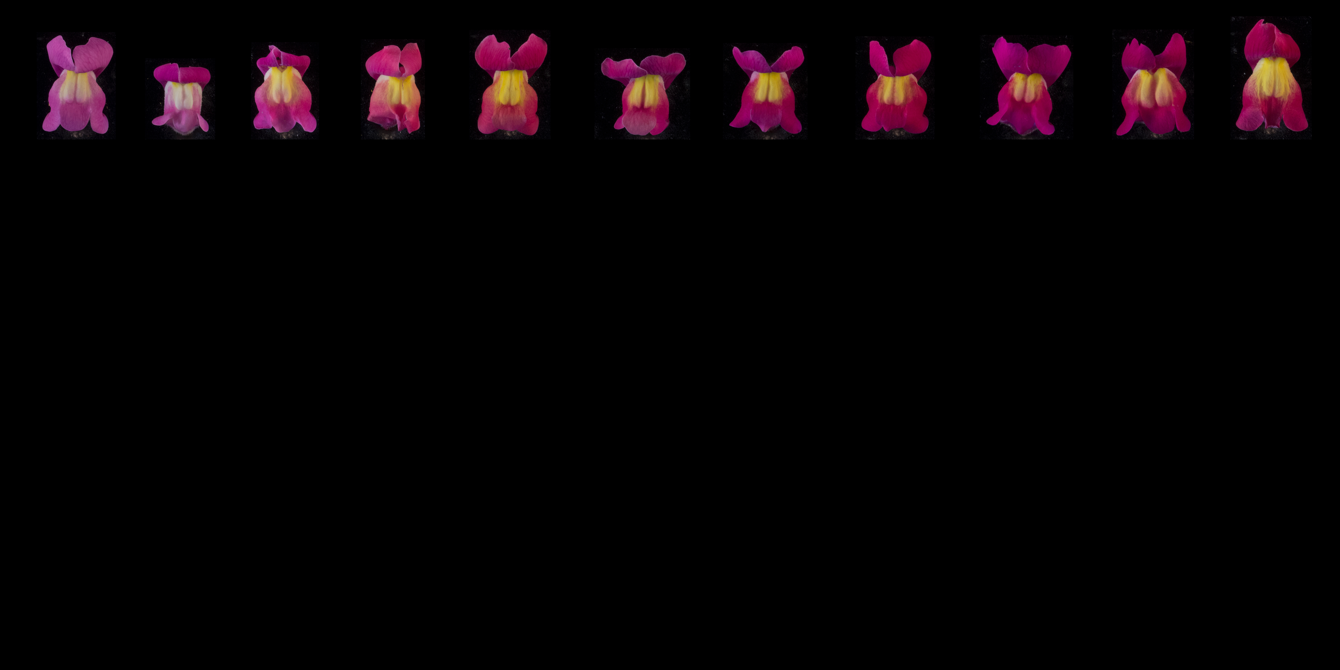

*ROS^p^ / ros^s^ EL^s^ / el^p^ SULF ^p^ / -*

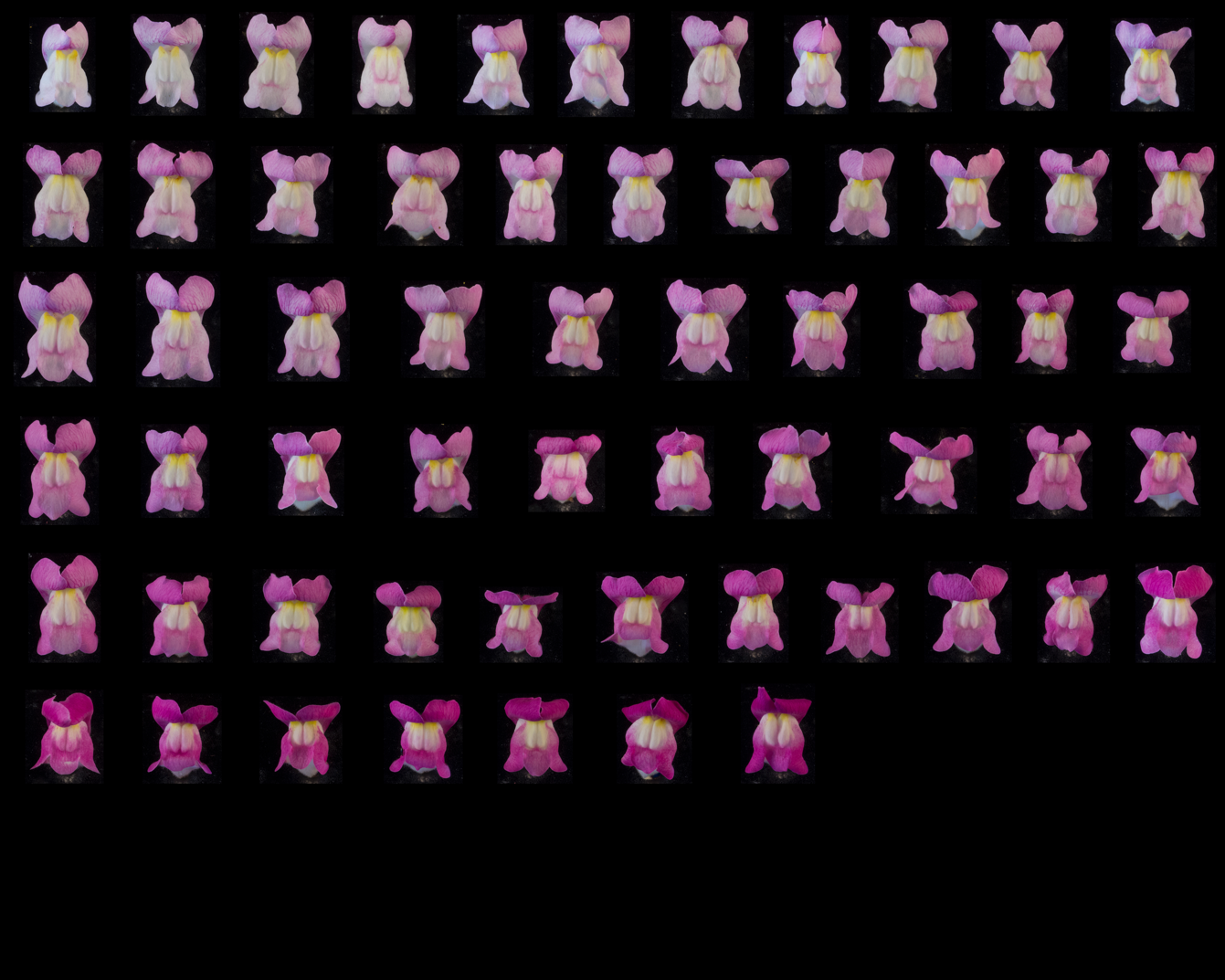

*ROS ^p^ / ros ^s^ EL^s^ / el ^p^ sulf ^s^ / sulf ^s^*

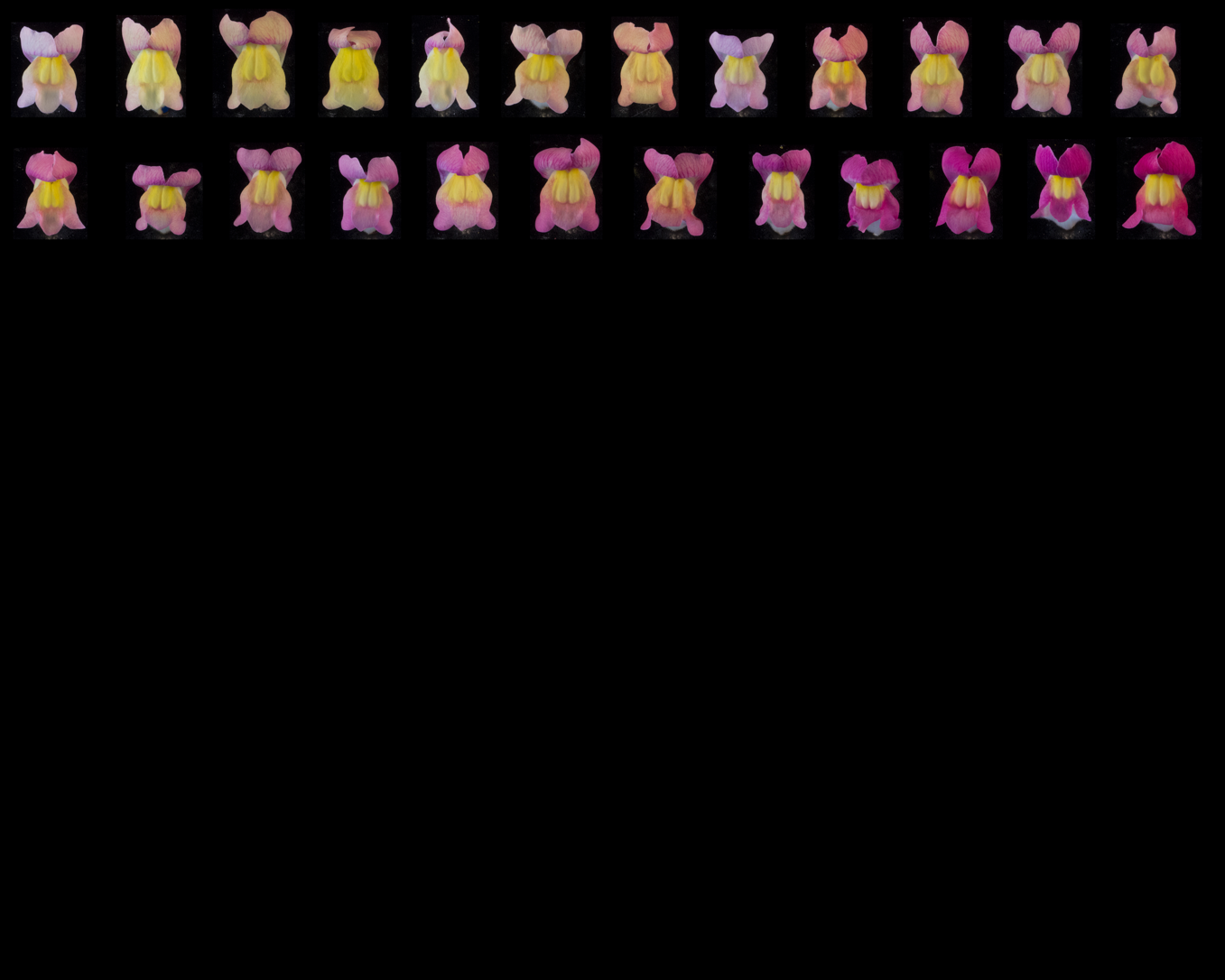

*ros^s^ / ros^s^ EL^s^ / EL^s^ SULF ^p^ / -*

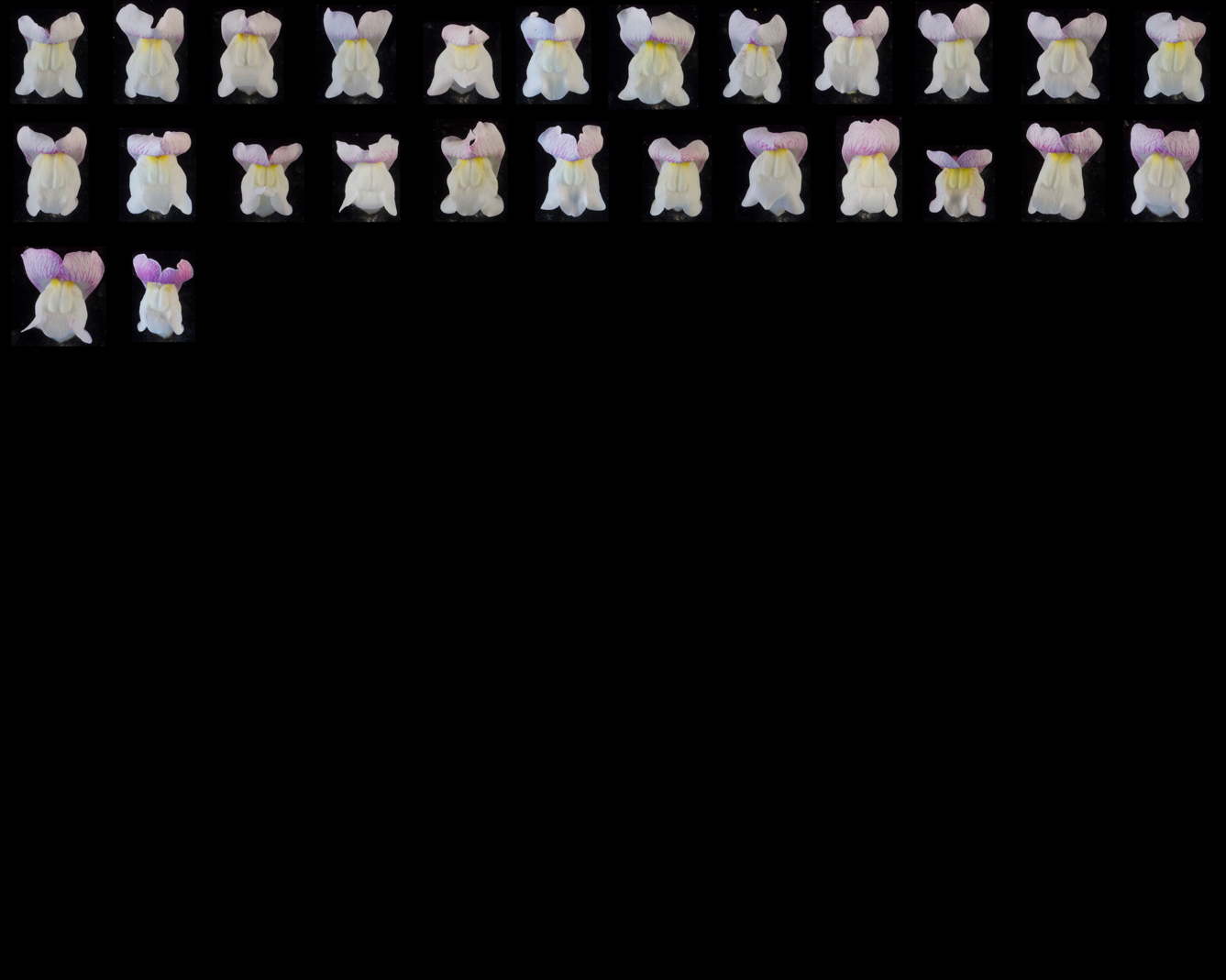

*ros^s^ / ros^s^ EL^s^ / EL^s^ sulf^s^ / sulf^s^*

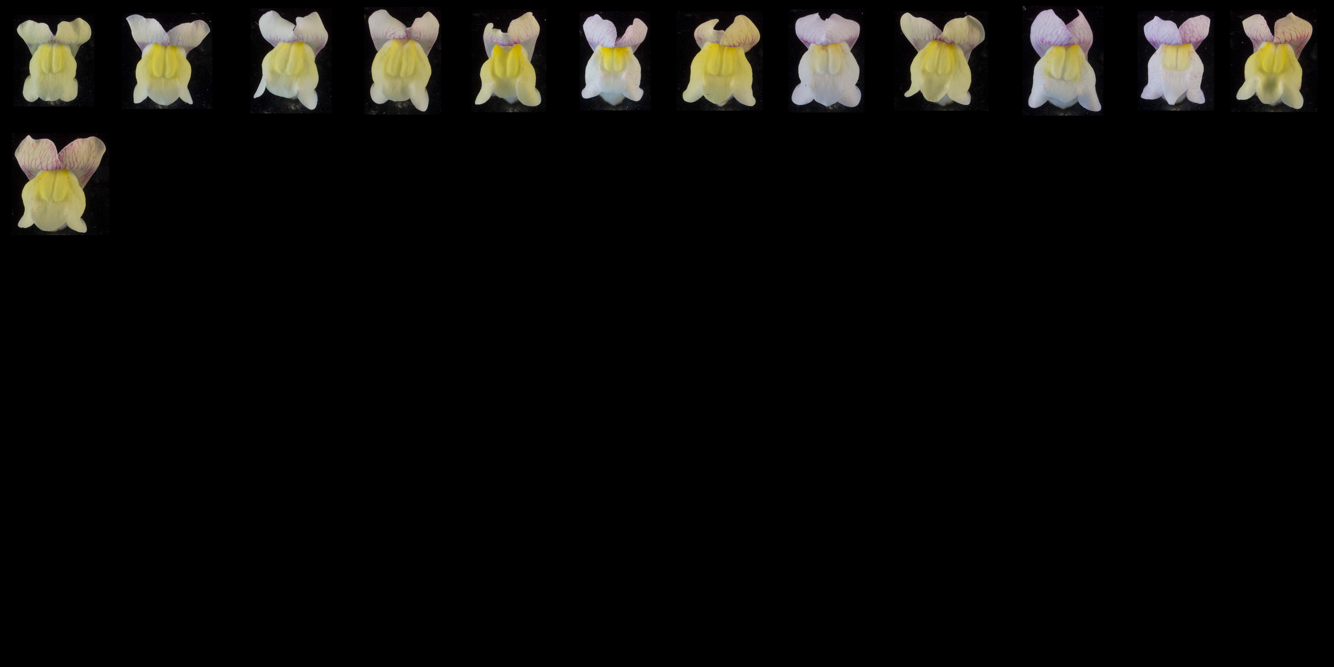

### **Supplementary Figure 12: Magenta colour rankings of F_2_ family J109, grouped by genotype**

Flowers ranked by extent of magenta: family J109 (F_2_ of *A. m. m.* var. *pseudomajus* x *A. m. m.* var. *striatum*). Individual accessions were grouped according to *ROS, EL*, and *SULF* genotype, before being ranked for the extent of magenta pigmentation. The flower deemed to show the weakest magenta was placed in the top left position, with the strength of magenta increasing along each row from top left to bottom right. Rankings were made without prior knowledge of island 1 or island 2 SNP genotypes. Superscript ‘s’ within the genotype denotes the *A. m. m.* var. *striatum* allele, and superscript ‘p’ denotes the *A. m. m.* var. *pseudomajus* allele.

*ROS ^p^ / ROS ^p^ el ^p^ / el ^p^ SULF ^p^ / -*

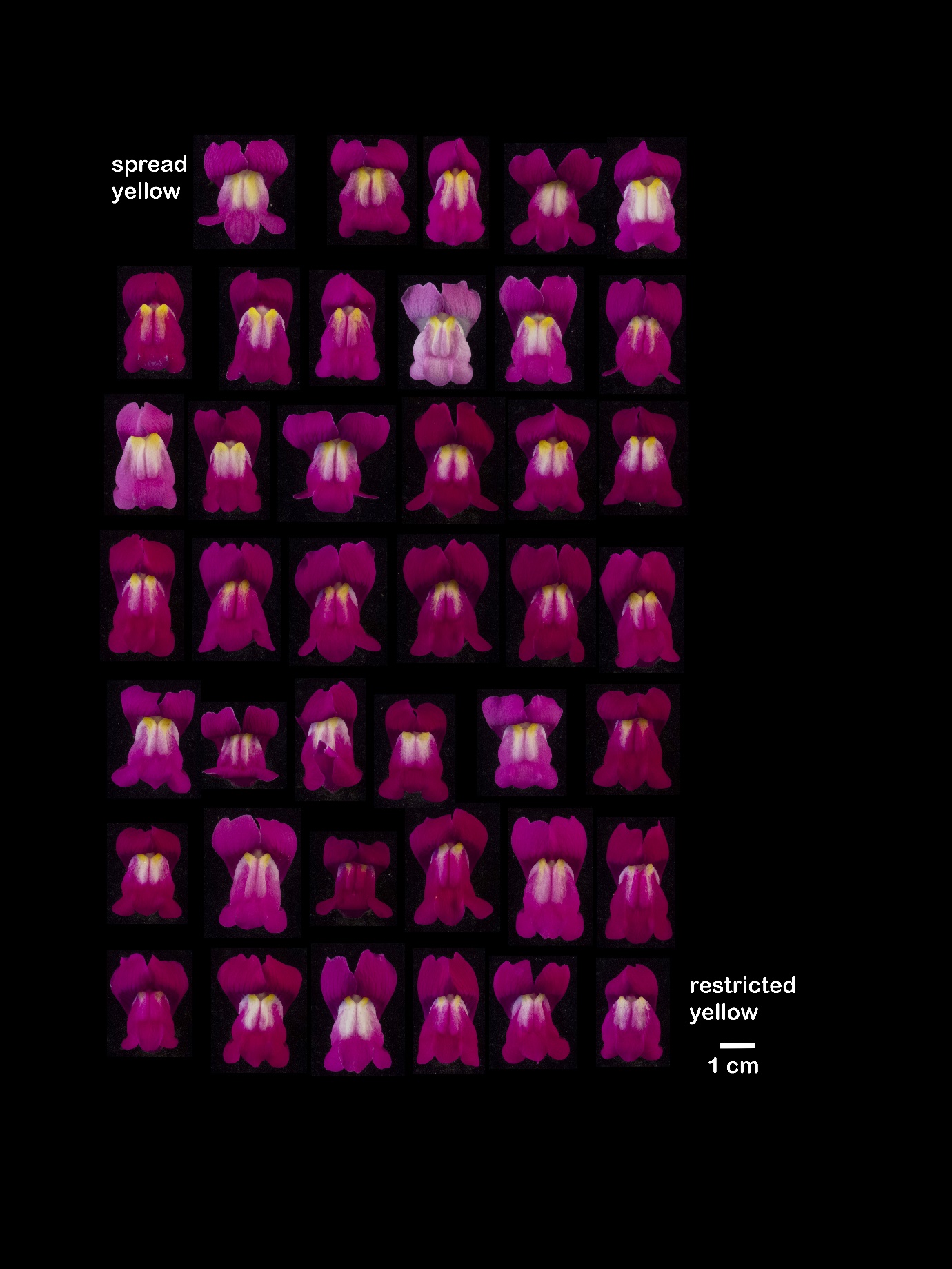

*ROS ^p^ / ROS ^p^ el ^p^ / el ^p^ sulf ^s^ / sulf ^s^*

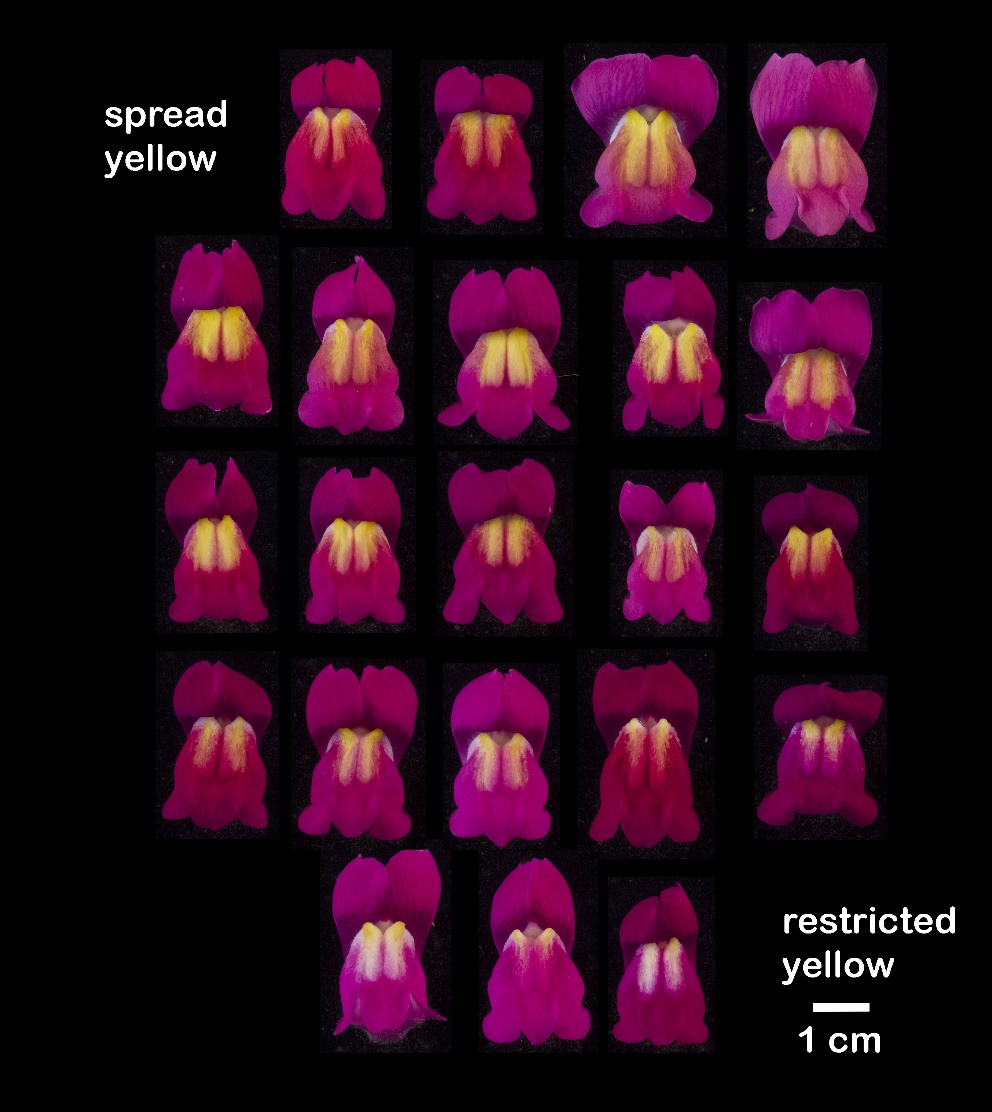

*ROS^p^ / ros^s^ EL^s^ / el^p^ SULF ^p^ / -*

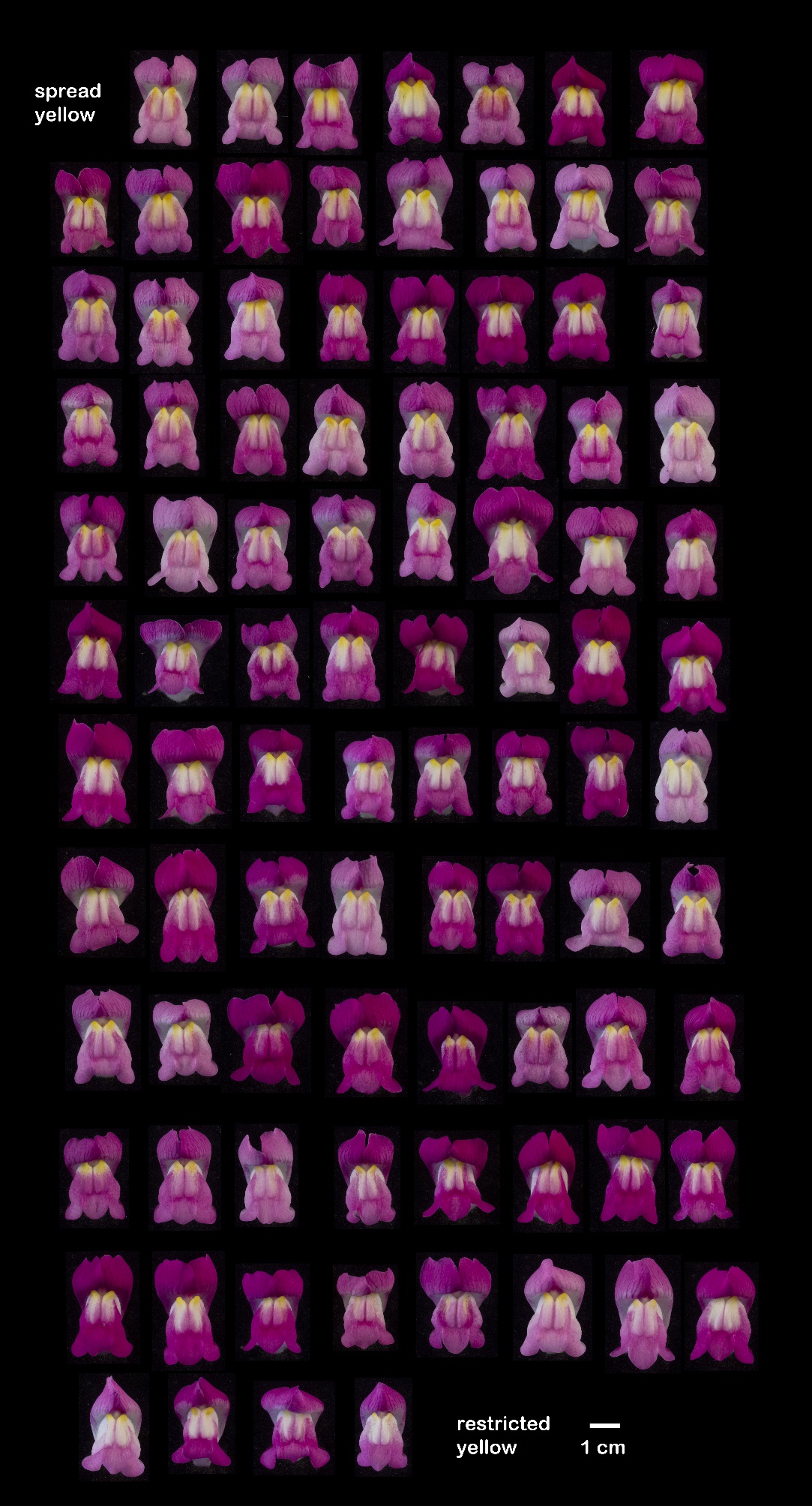

*ROS ^p^ / ros ^s^ EL^s^ / el ^p^ sulf ^s^ / sulf ^s^*

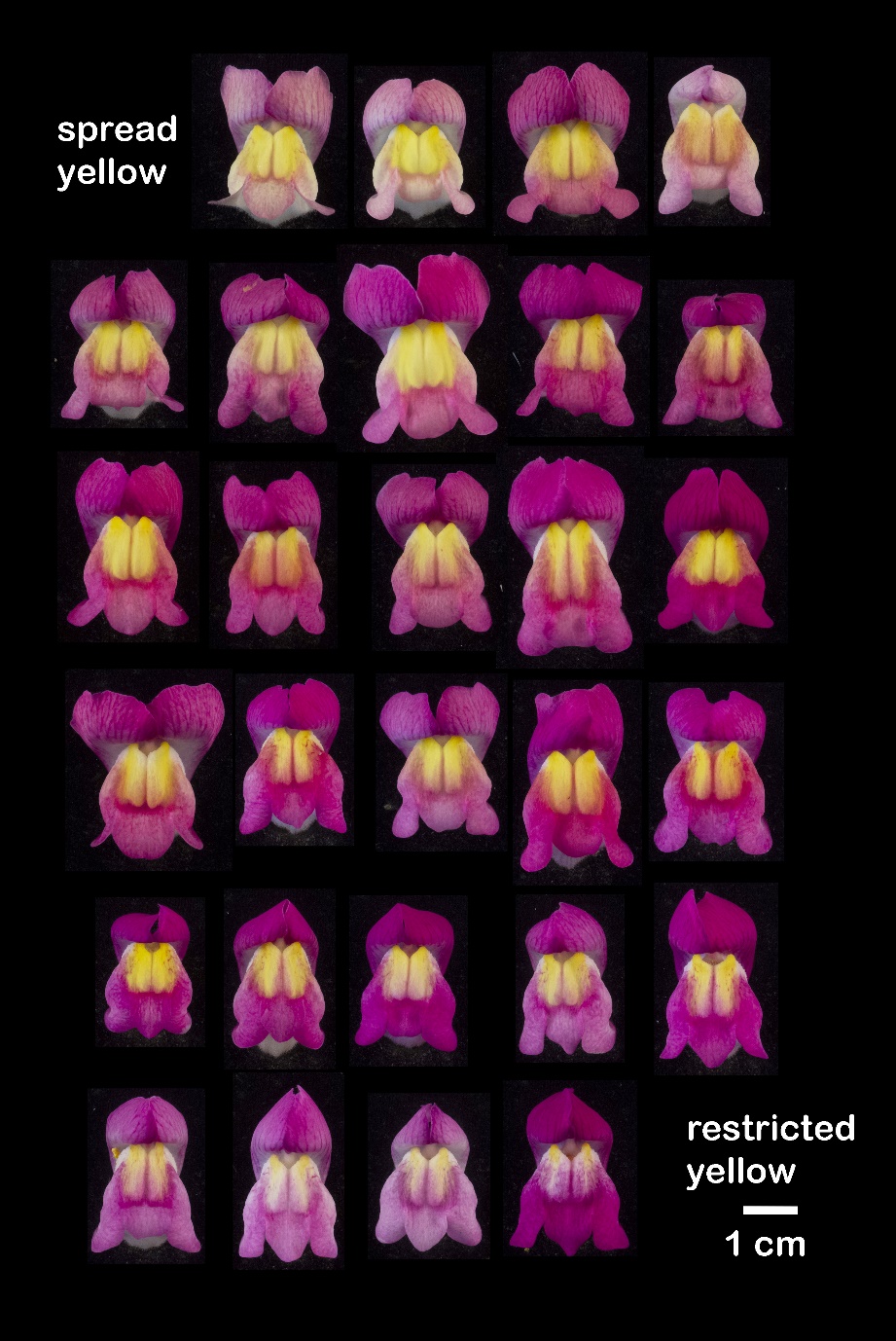

*ros^s^ / ros^s^ EL^s^ / EL^s^ SULF ^p^ / -*

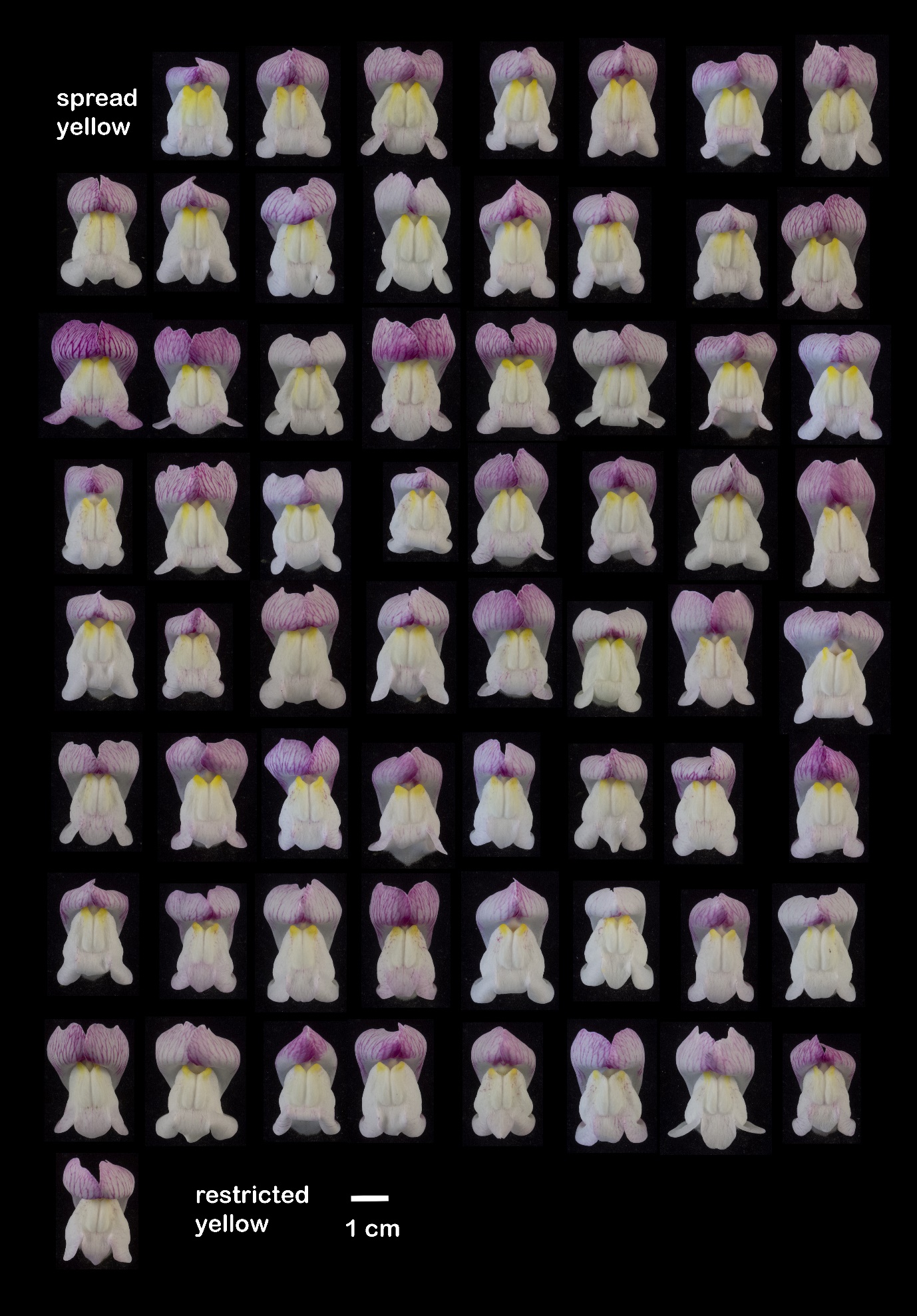

*ros^s^ / ros^s^ EL^s^ / EL^s^ sulf^s^ / sulf^s^*

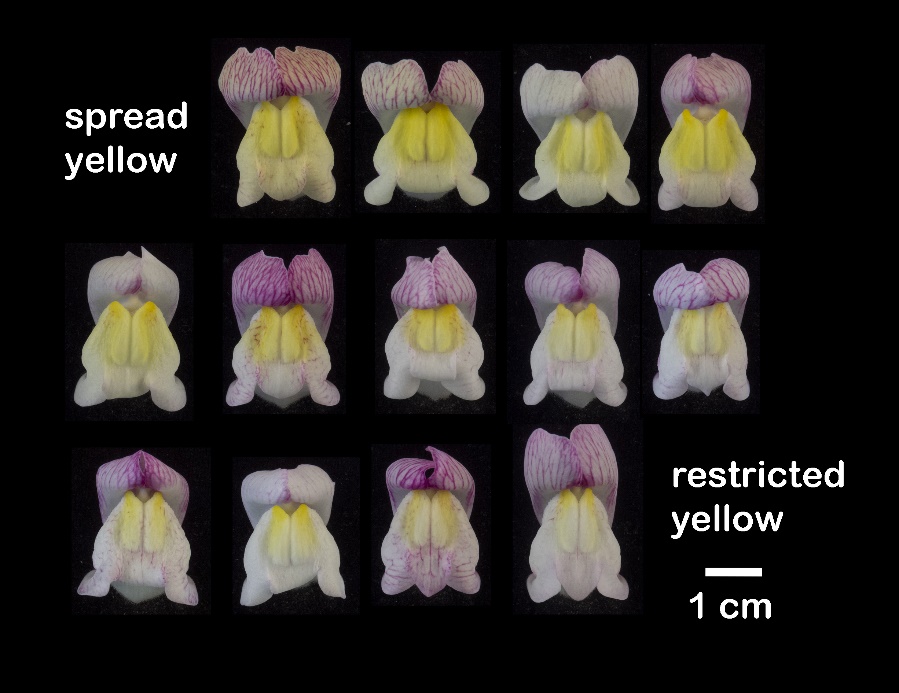

### **Supplementary Figure 13: Yellow colour rankings of F_2_ family C197, grouped by genotype**

Flowers ranked by extent of yellow: family C197 (F_2_ of *A. m. m.* var. *pseudomajus* x *A. m. m.* var. *striatum*). Individual accessions were grouped according to *ROS, EL*, and *SULF* genotype, before being ranked for the extent of yellow pigmentation. The flower deemed to show the most restricted yellow was placed in the bottom right position, with the amount of yellow increasing along each row from bottom right to top left. Rankings were made without prior knowledge of island 8 SNP genotypes. Superscript ‘s’ within the genotype denotes the *A. m. m.* var. *striatum* allele, and superscript ‘p’ denotes the *A. m. m.* var. *pseudomajus* allele.

*ROS ^p^ / ROS ^p^ el ^p^ / el ^p^ SULF ^p^ / -*

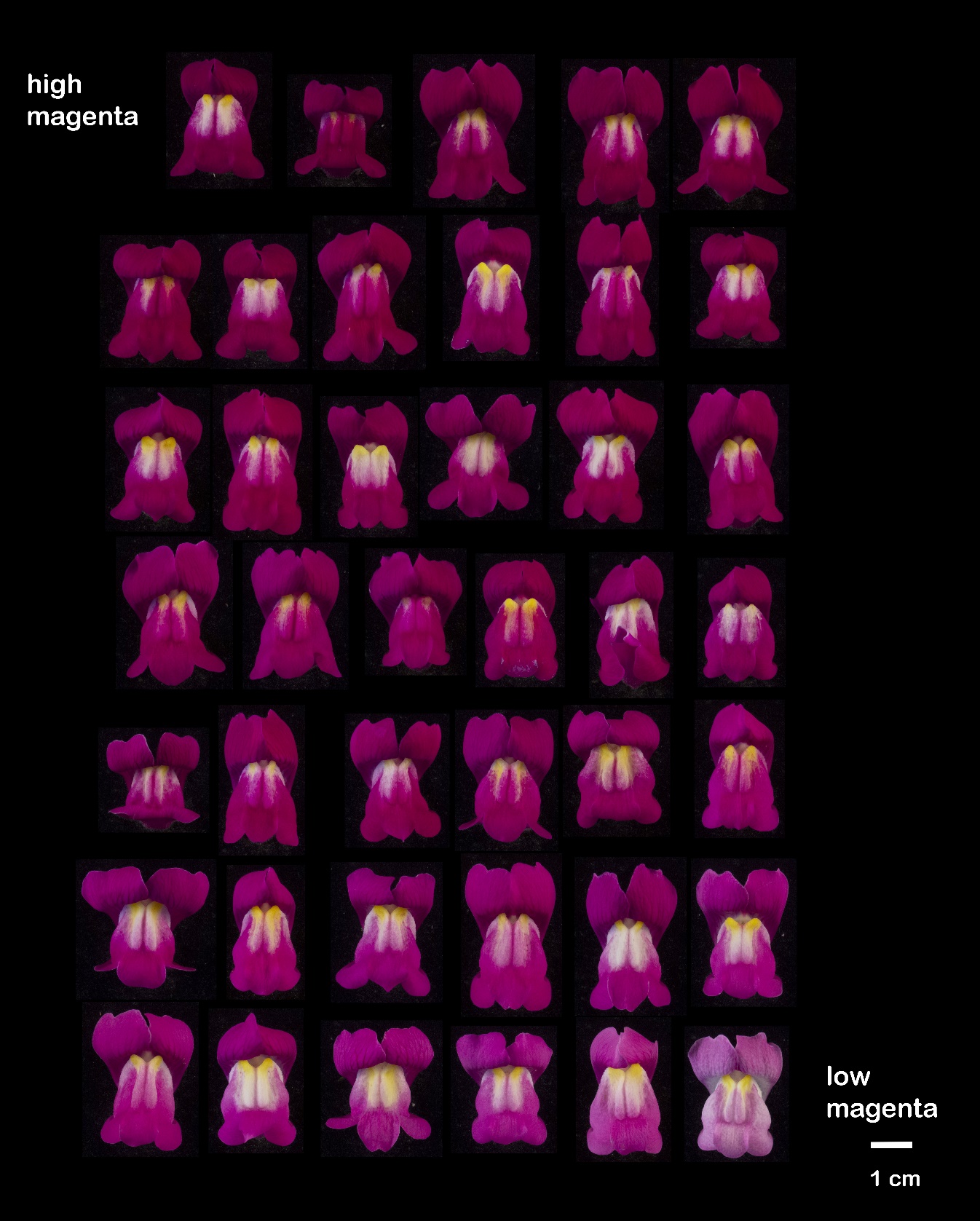

*ROS ^p^ / ROS ^p^ el ^p^ / el ^p^ sulf ^s^ / sulf ^s^*

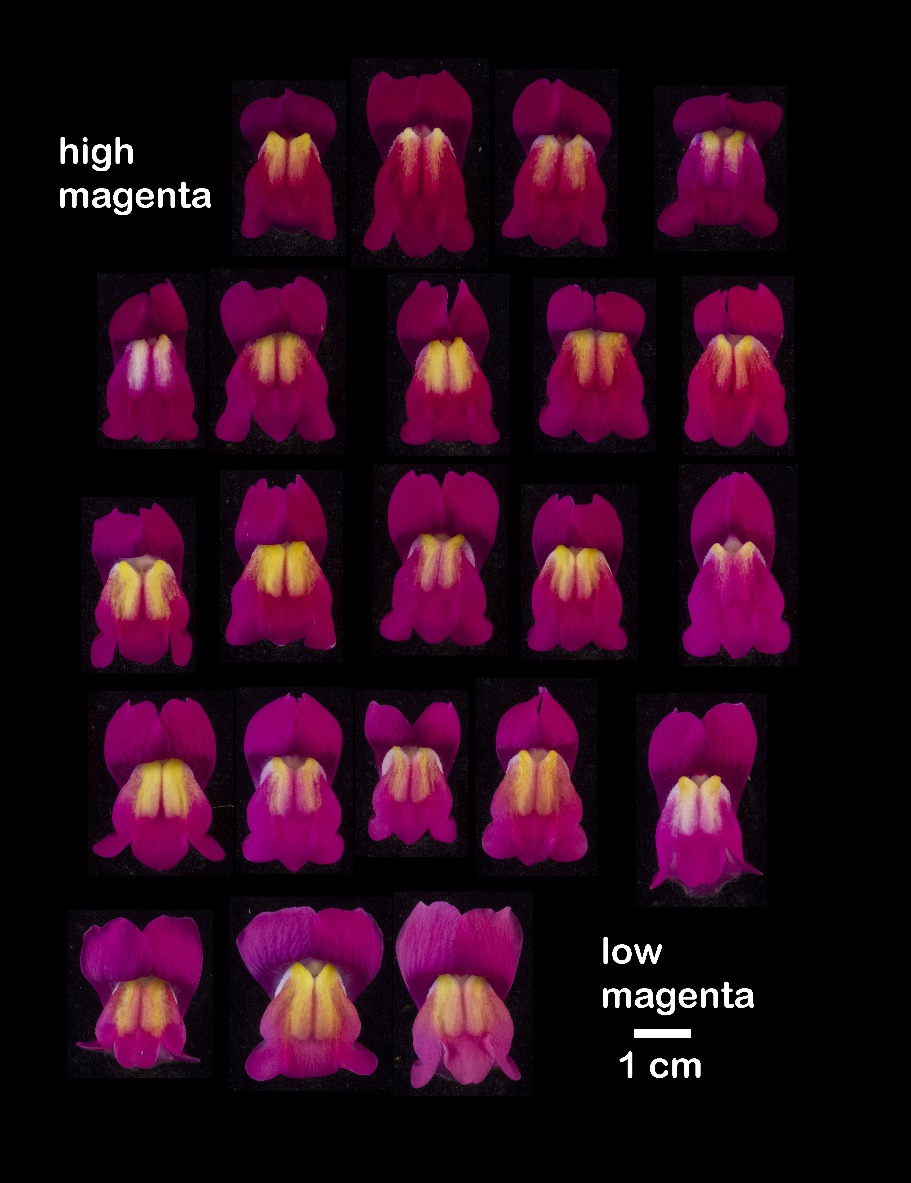

*ROS^p^ / ros^s^ EL^s^ / el^p^ SULF ^p^ / -*

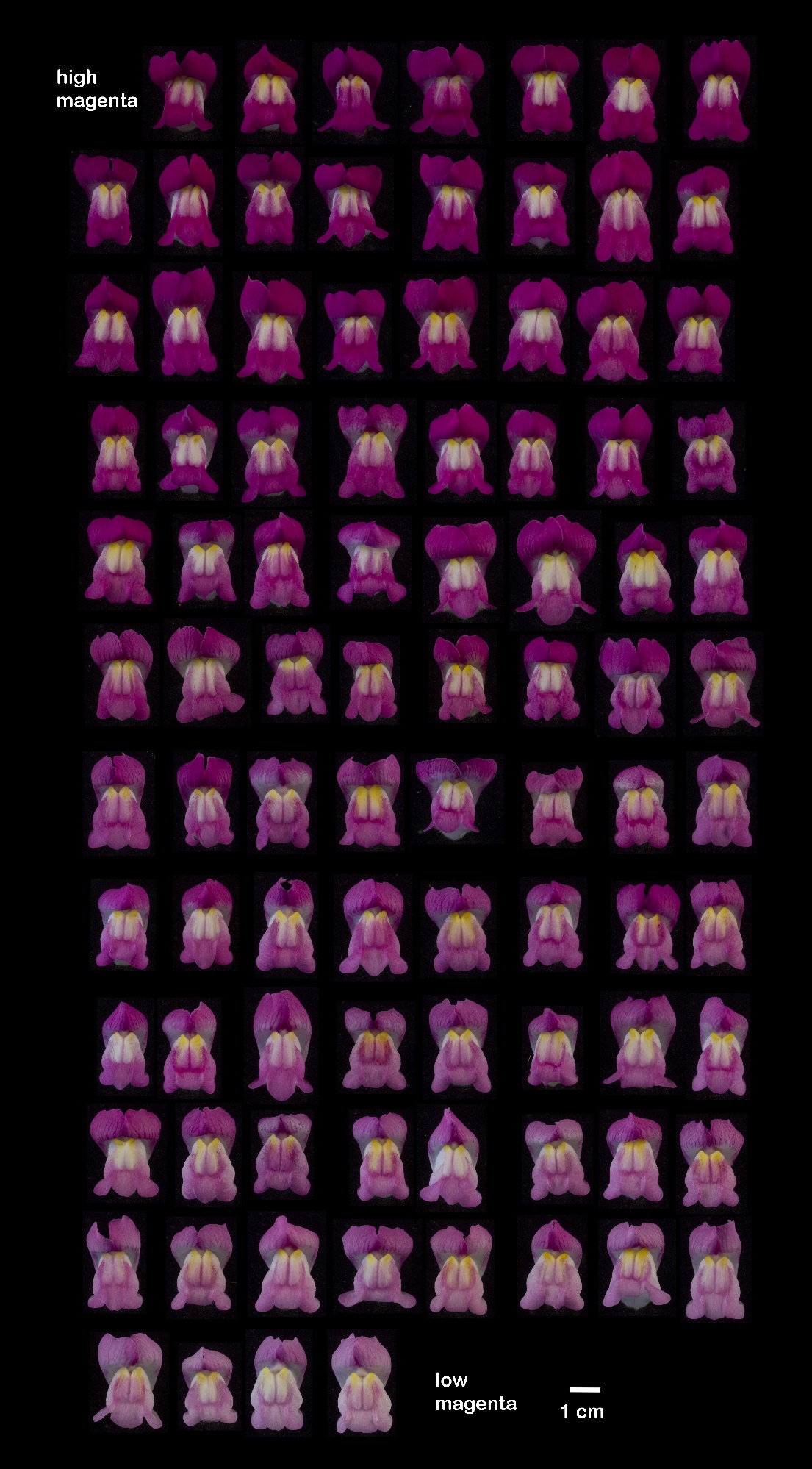

*ROS ^p^ / ros ^s^ EL^s^ / el ^p^ sulf ^s^ / sulf ^s^*

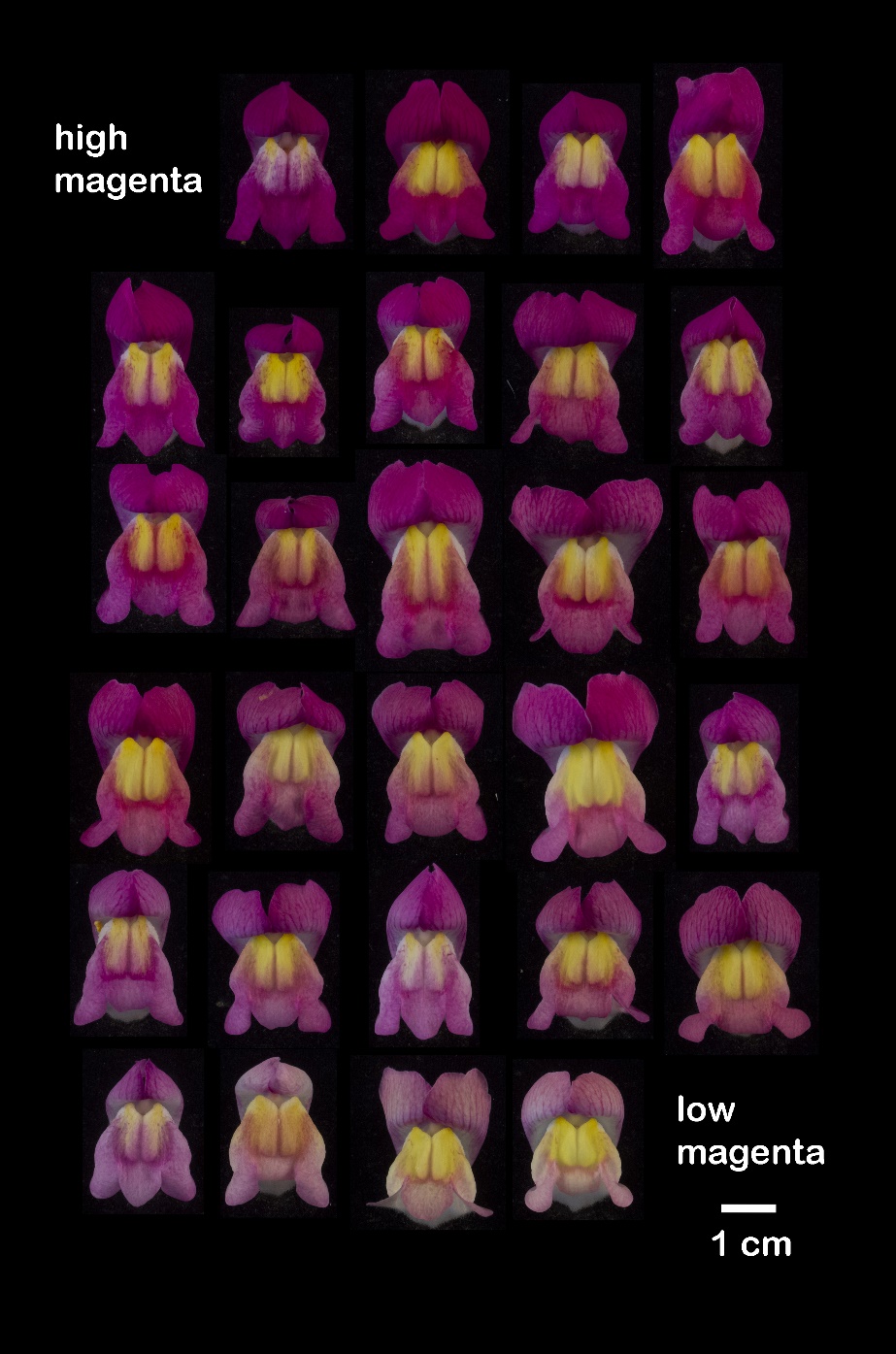

*ros^s^ / ros^s^ EL^s^ / EL^s^ SULF ^p^ / -*

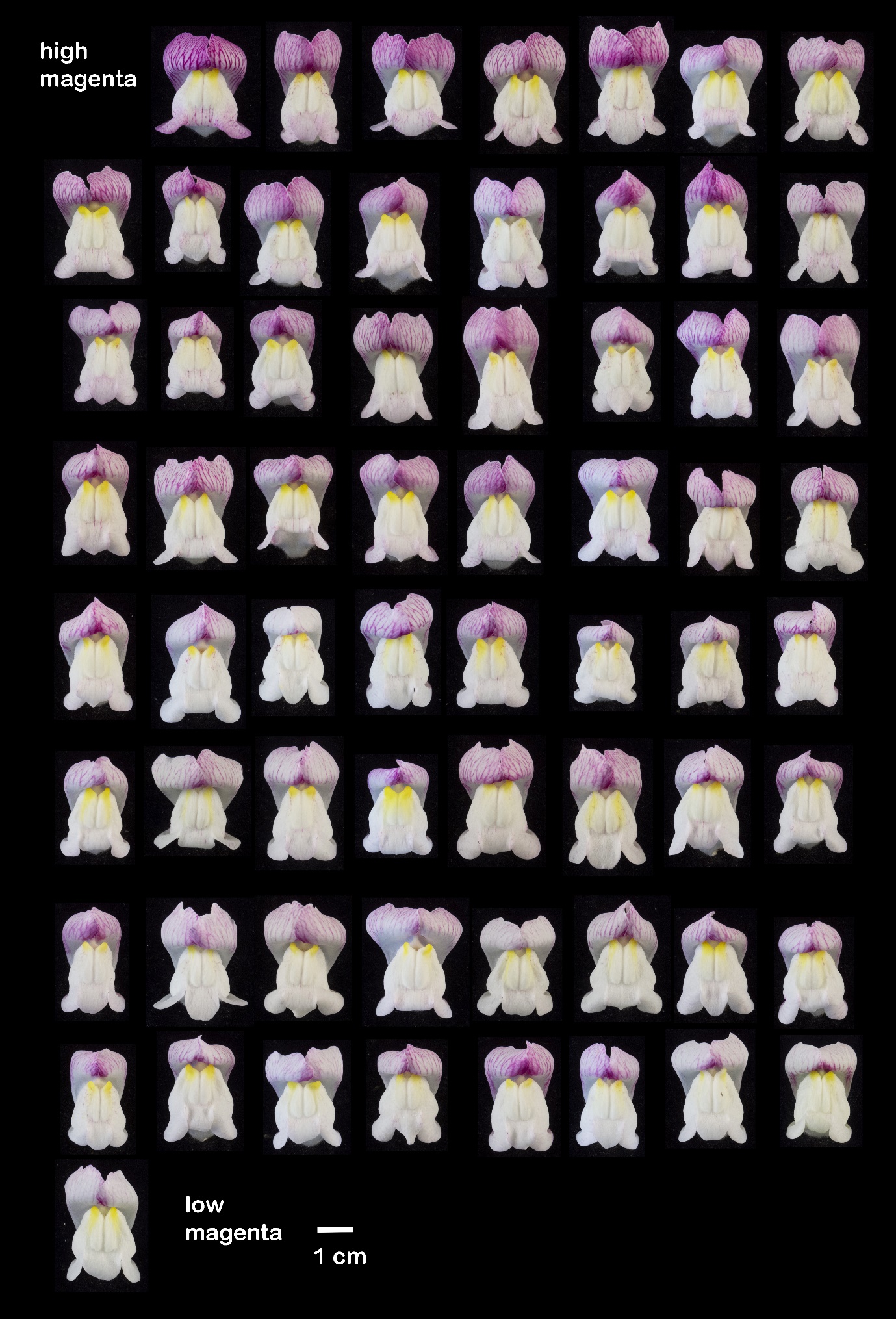

*ros^s^ / ros^s^ EL^s^ / EL^s^ sulf^s^ / sulf^s^*

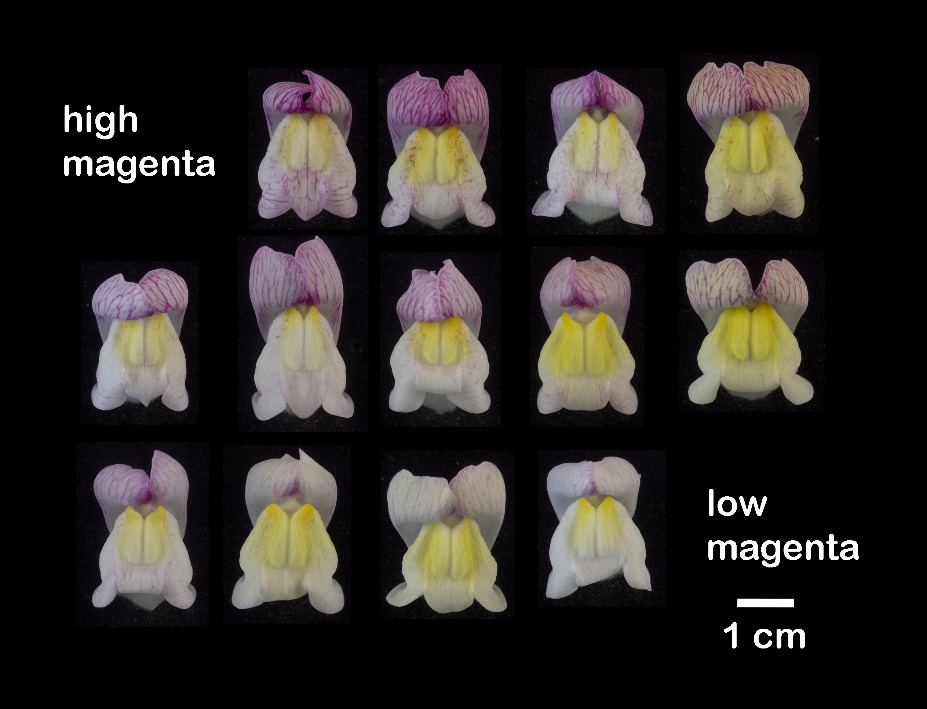

### **Supplementary Figure 14: Magenta colour rankings of F_2_ family C197, grouped by genotype**

Flowers ranked by extent of magenta: family C197 (F_2_ of *A. m. m.* var. *pseudomajus* x *A. m. m.* var. *striatum*). Individual accessions were grouped according to *ROS, EL*, and *SULF* genotype, before being ranked for the extent of magenta pigmentation. The flower deemed to show the weakest magenta was placed in the bottom right position, with the strength of magenta increasing along each row from bottom right to top left. Rankings were made without prior knowledge of island 8 SNP genotypes. Superscript ‘s’ within the genotype denotes the *A. m. m.* var. *striatum* allele, and superscript ‘p’ denotes the *A. m. m.* var. *pseudomajus* allele.

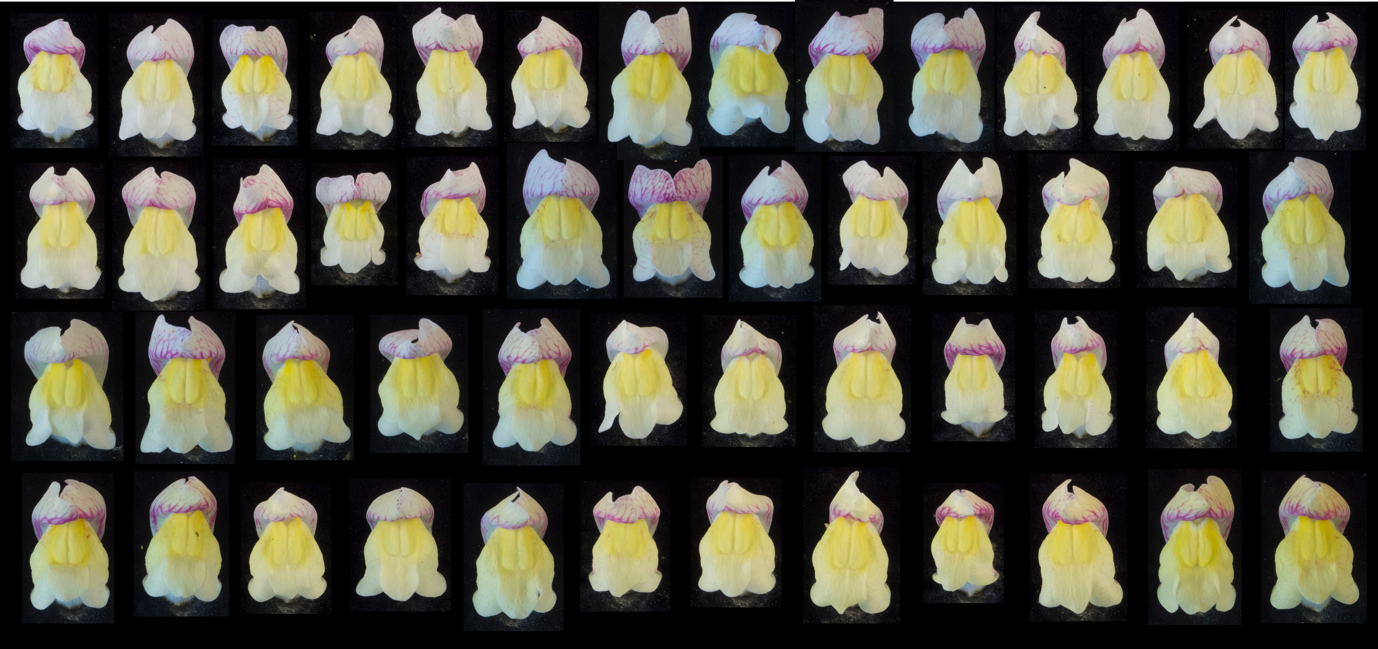

### **Supplementary Figure 15: Yellow colour ranking analysis of an F_4_ family segregating for island 1 SNPs (Replicate 1)**

Flowers from F_4_ individuals segregating for island 1 SNPs were ranked for the extent of yellow pigmentation. The flower deemed to show the most restricted yellow was placed in the top left position, with the amount of yellow increasing along each row from top left to bottom right. In each case, rankings were made without knowing the island 1 SNP genotypes. Rankings were carried out by three independent researchers, who all found similar results: Daniel Richardson (Replicate 1, Supplementary Figure 15), Desmond Bradley (Replicate 2, Supplementary Figure 16) and Tingting Li (Replicate 3, Supplementary Figure 17).

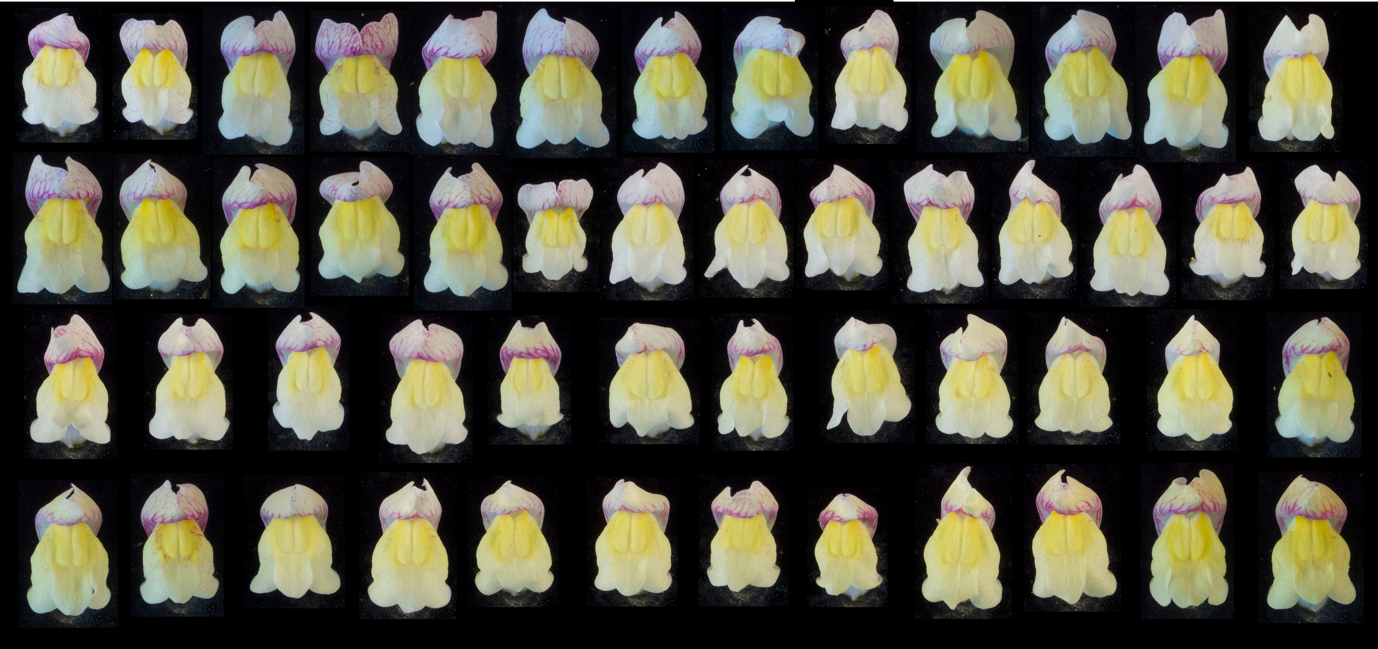

### **Supplementary Figure 16: Yellow colour ranking analysis of an F_4_ family segregating for island 1 SNPs (Replicate 2)**

Replicate ranking carried out by Desmond Bradley.

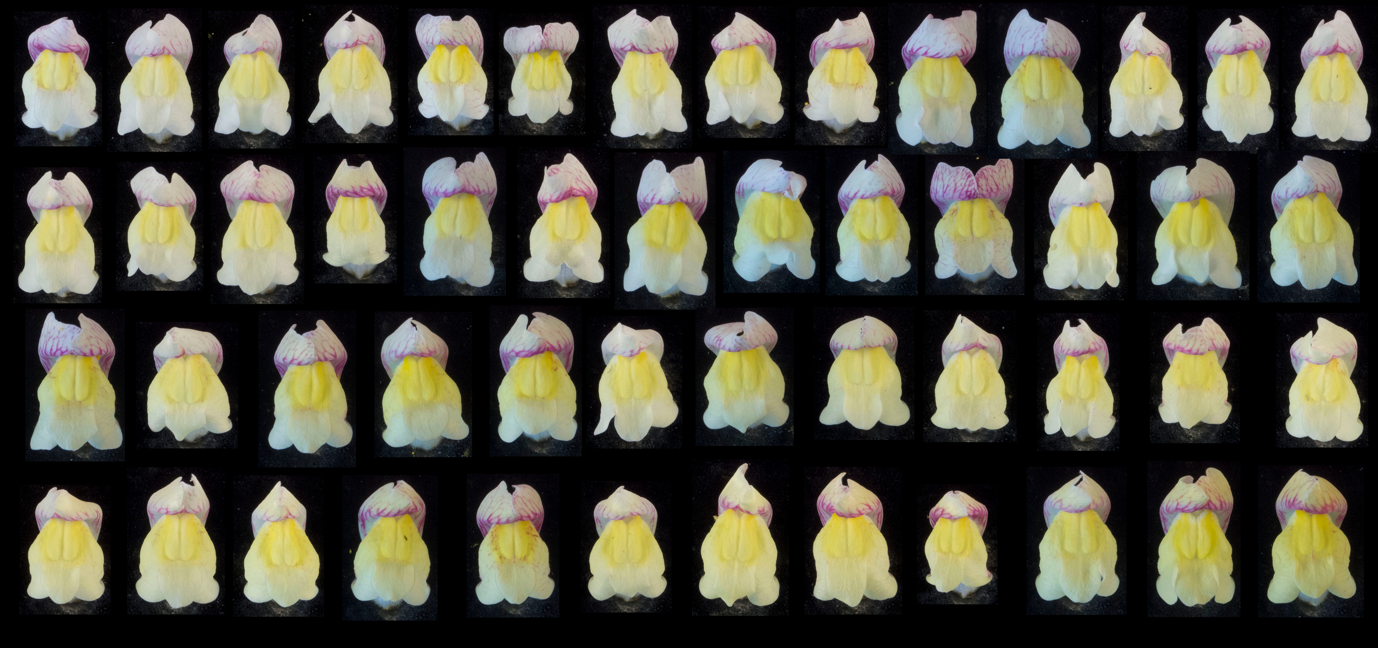

### **Supplementary Figure 17: Yellow colour ranking analysis of an F_4_ family segregating for island 1 SNPs (Replicate 3)**

Replicate ranking carried out by Tingting Li.

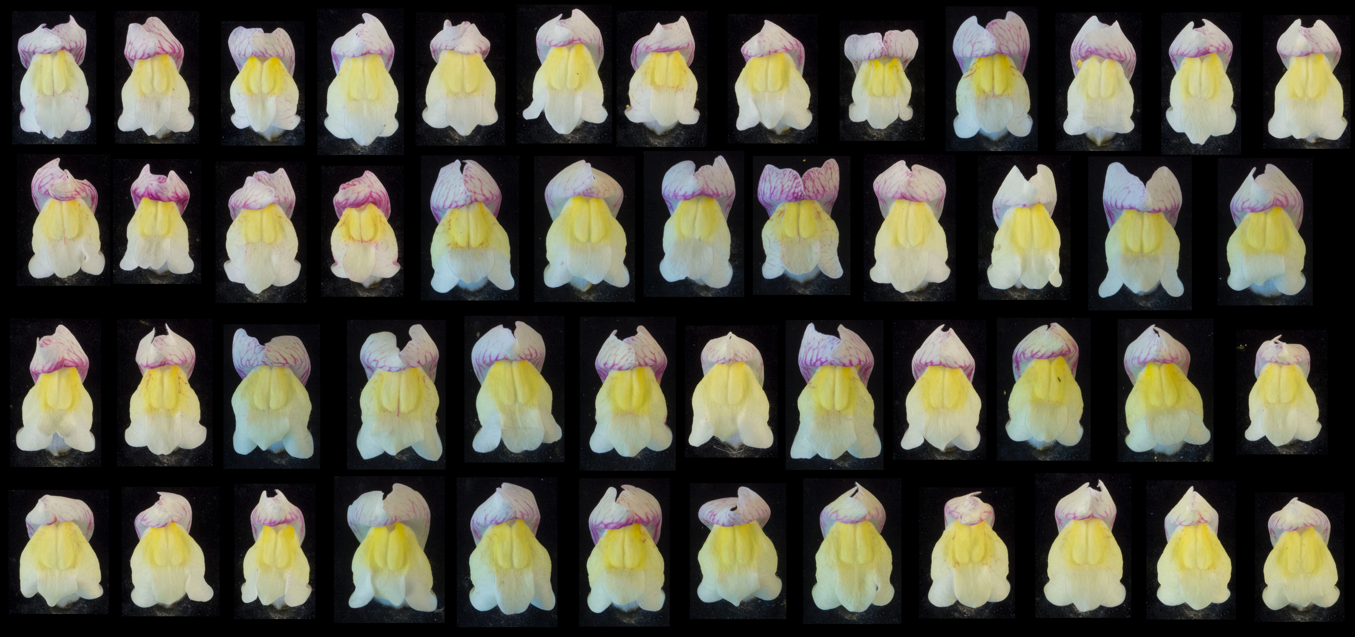

### **Supplementary Figure 18: Yellow colour ranking analysis of an F_4_ family segregating for island 2 SNPs (Replicate 1)**

Flowers from F_4_ individuals segregating for island 2 SNPs were ranked for the extent of yellow pigmentation. The flower deemed to show the most restricted yellow was placed in the top left position, with the amount of yellow increasing along each row from top left to bottom right. In each case, rankings were made without knowing the island 2 SNP genotypes. Rankings were carried out by three independent researchers, who all found similar results: Daniel Richardson (Replicate 1, Supplementary Figure 18), Desmond Bradley (Replicate 2, Supplementary Figure 19) and Tingting Li (Replicate 3, Supplementary Figure 20).

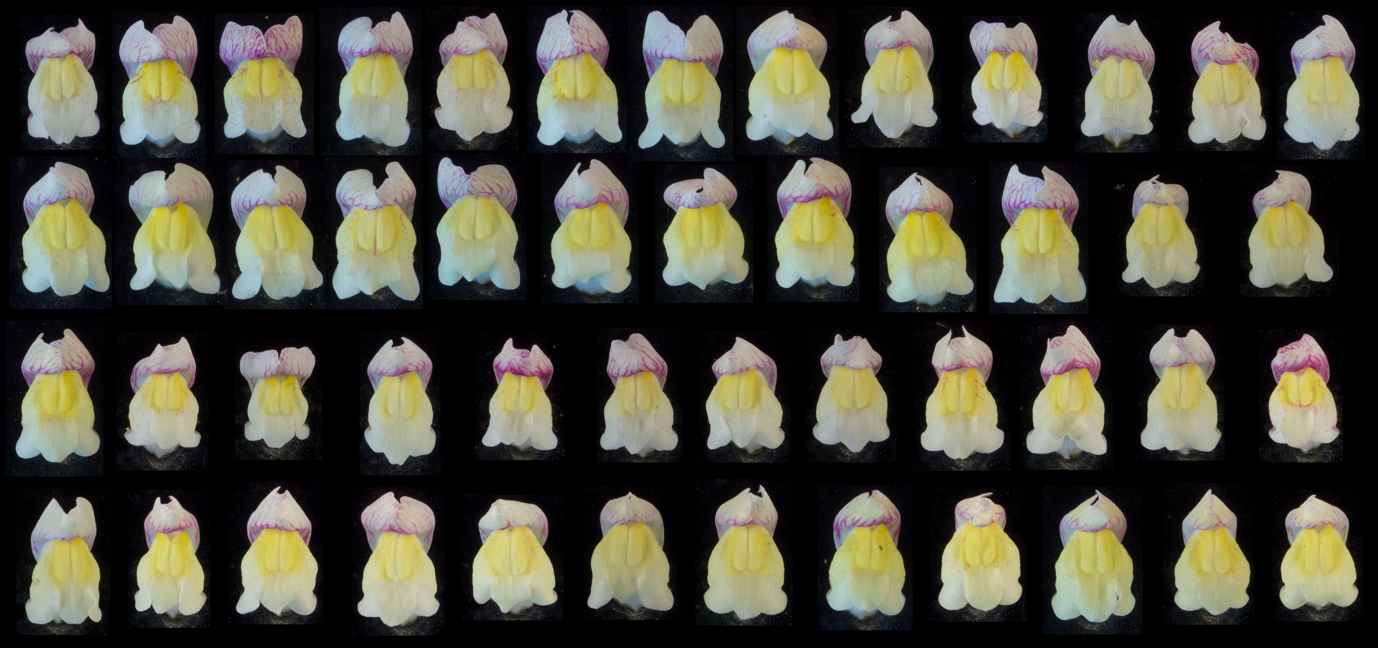

### **Supplementary Figure 19: Yellow colour ranking analysis of an F_4_ family segregating for island 2 SNPs (Replicate 2)**

Replicate ranking carried out by Desmond Bradley.

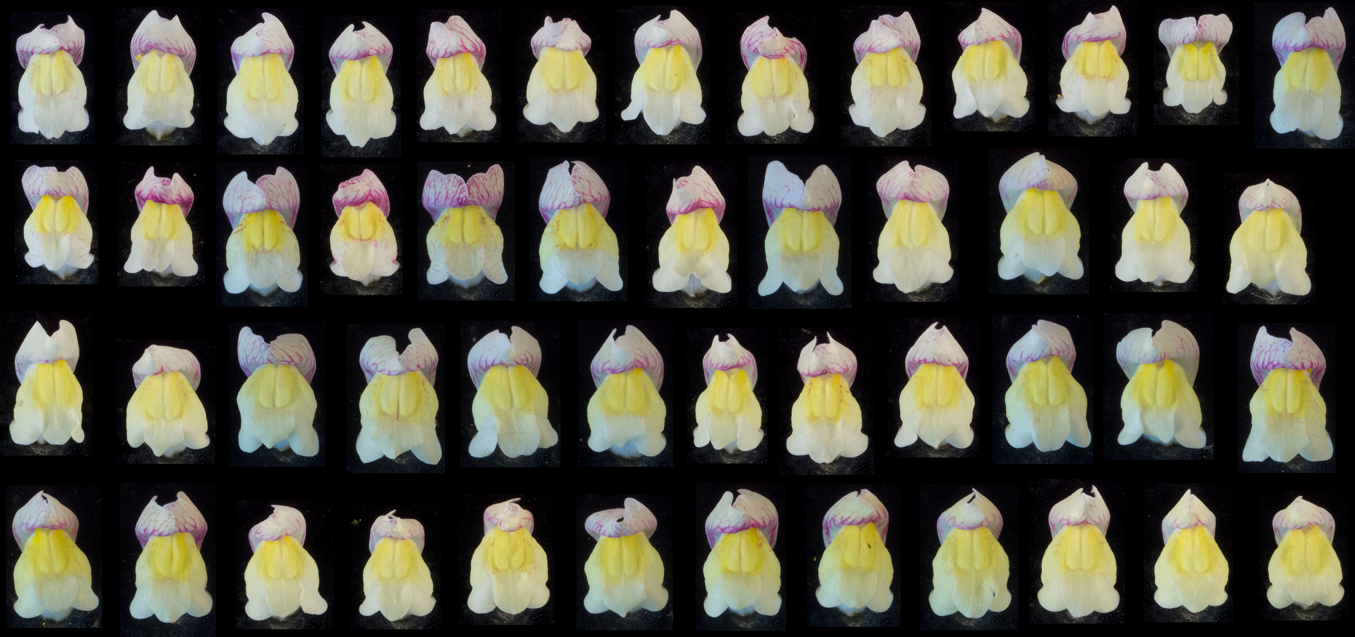

### **Supplementary Figure 20: Yellow colour ranking analysis of an F_4_ family segregating for island 2 SNPs (Replicate 3)**

Replicate ranking carried out by Tingting Li.

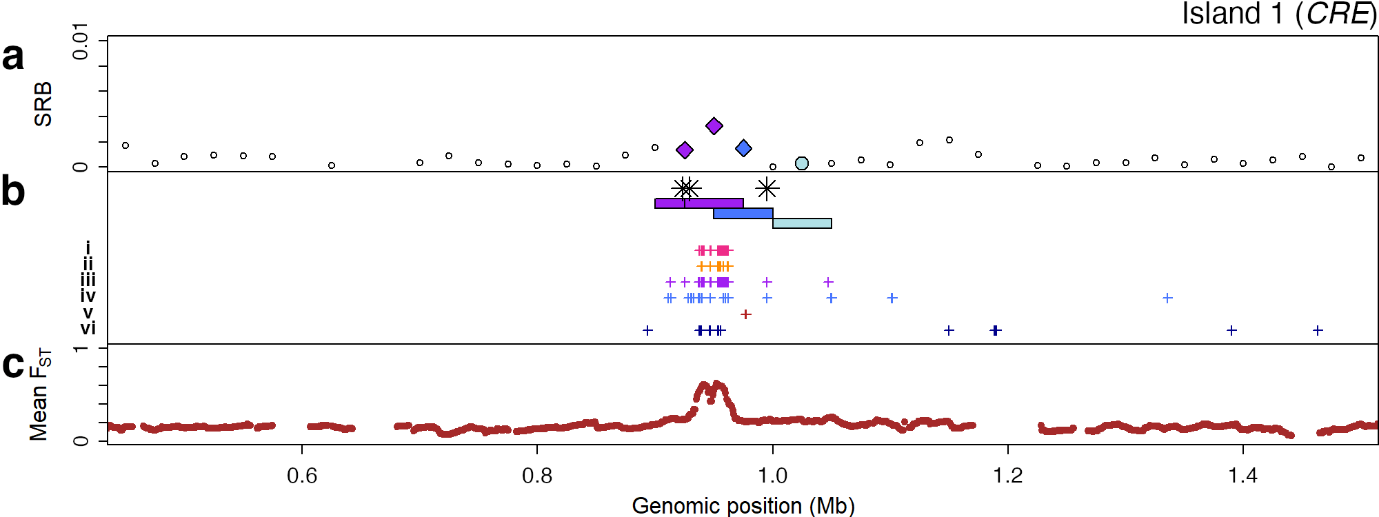

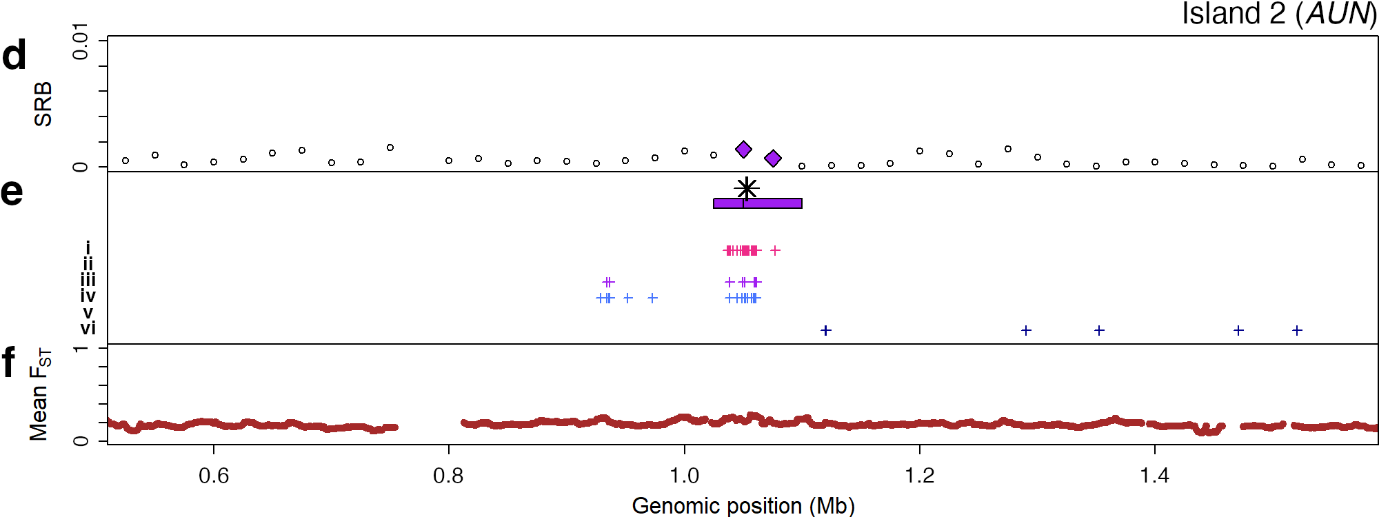

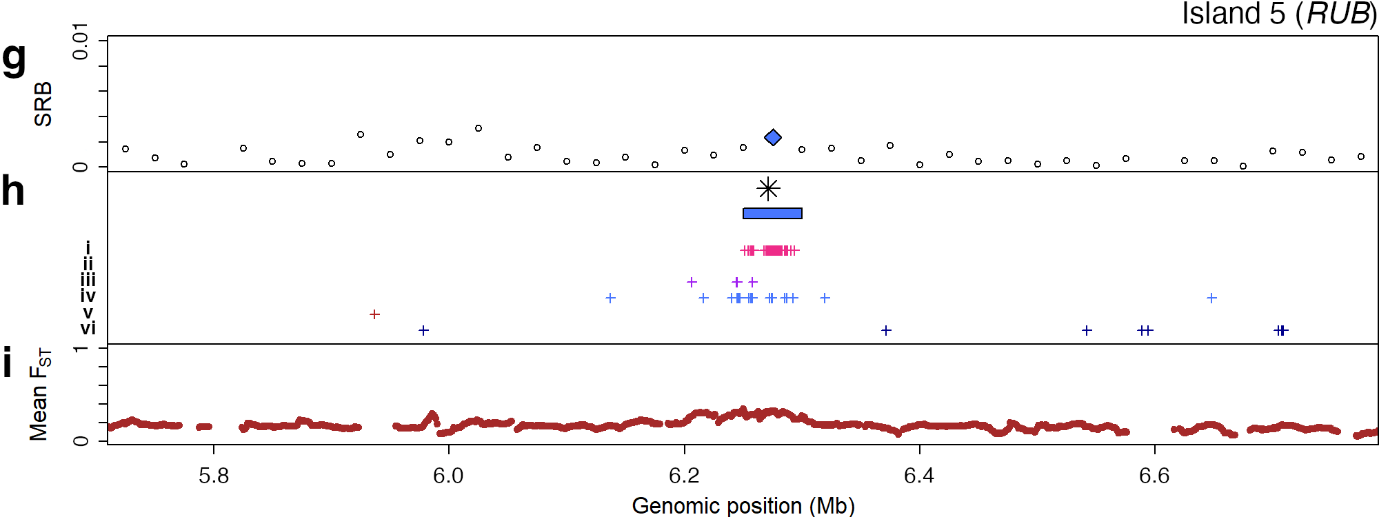

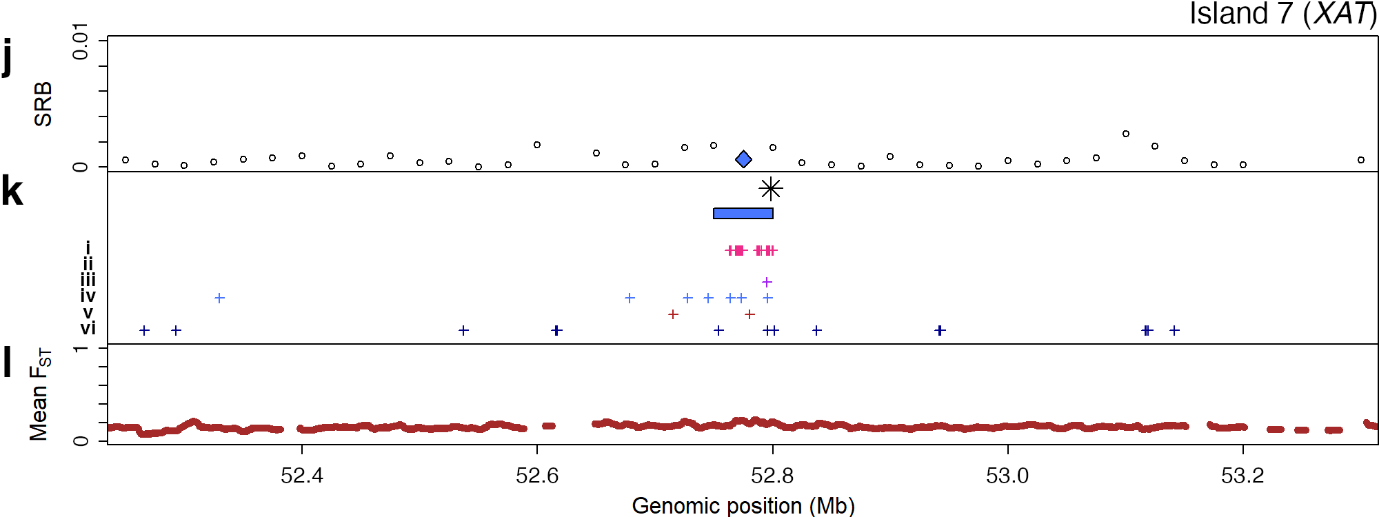

### **Supplementary Figure 21: 1 Mb regions of partition islands 1, 2, 5 and 7**

**a, d, g, j** SRB for 50 kb window trees across three partition islands. Coloured diamonds indicate trees giving the A partition (purple), the A’ partition (dark blue), or the A’’ partition (light blue) in > 50% of bootstrap replicates (Main text Fig. 3g). **b, e, h, k** 50 kb windows (rectangles, colour-coded as above) and SNPs below showing (i) The top 100 island SNPs showing the highest mean frequency difference between pseudomajus (A1) and striatum (A2) populations (magenta crosses) (ii) Those with mean allele frequency differences >= 0.8 between A1 and A2 populations (orange crosses). (iii) Partition-A SNP_XY_ trees (purple crosses). (iv) Partition-A’ SNP_XY_ trees (lighter blue crosses). (v) SNPs showing a mean read depth difference > 0.95, depleted in pseudomajus. (vi) SNPs showing a mean read depth difference > 95%, depleted in striatum. The locations of candidate flower colour loci are indicated by asterisks. **c, f, i, l** Mean F_ST_-A, calculated from averages of π_w_, pairwise d_XY_ and pairwise π_t_, across all A1 and A2 populations, averaged in 10 kb windows with 9 kb overlaps.

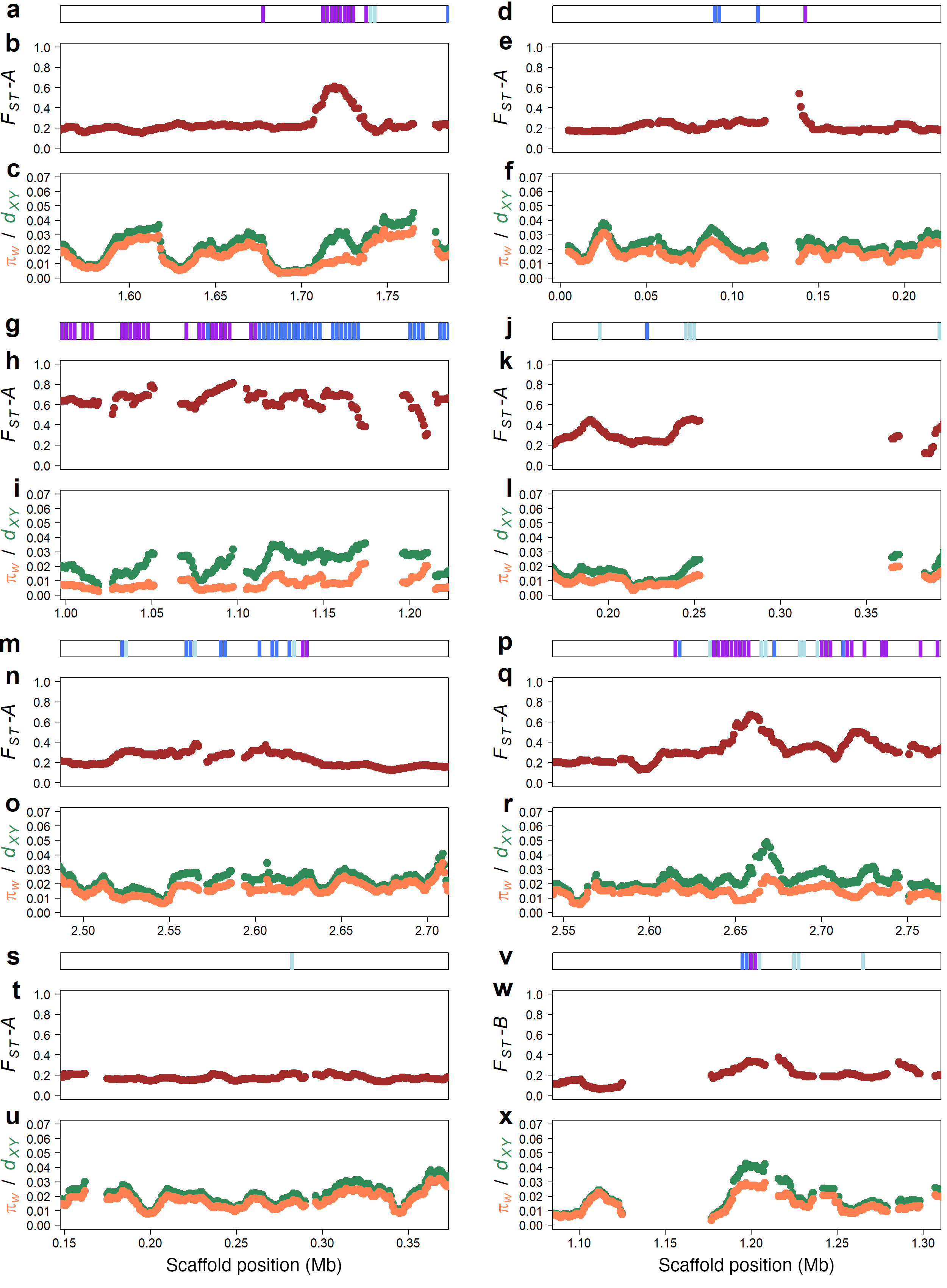

### **Supplementary Figure 22: Partition island genome scans for tree partition, F_ST_, π_w_ and d_XY_ across partition islands in *A. m. m. var. pseudomajus***

**a, d, g, j, m, p, s** Positions of 5 kb windows showing the A partition (purple), the A’ partition (one population misgrouped compared to the A partition, dark blue), or the A’’ partition (two populations misgrouped compared to the A partition, light blue) in partition islands 1-7, mapped to an *A. m. m. var. pseudomajus* genome assembly, which was used to enable detection of indel sequences which may not be present within the *A. majus* reference genome (such as *SCL8-like*). **b, e, h, k, n, q, t** Mean *F_ST_* for pairwise comparisons of all Group A1 and A2 populations, averaged in 10 kb windows with 9 kb overlaps across partition islands. **c, f, I, l, o, r, u** Mean *d_XY_* (green) and *π_w_* (orange) for Group A1 and A2 populations across partition islands. **v, w, x** As (a), (b), (c), but calculating statistics between B1 and B2 for partition island 8.

### **Supplementary Table 1: Details of sampled populations**

| **Supplementary Table 1: Details of sampled populations.** | | | | | |
| --- | --- | --- | --- | --- | --- |
| **Population identifier** | **Variety** | **Number of individuals sampled** | **Latitude** | **Longitude** | **Altitude (m)** |
| UNA | pseudomajus | 60 | 42.763361 | 1.772739 | 678 |
| BED | pseudomajus | 44 | 42.869186 | 1.568953 | 618 |
| LU | striatum | 47 | 42.968486 | 2.260464 | 227 |
| AXA | striatum | 32 | 42.798143 | 2.2231055 | 445 |
| MIJ | striatum | 44 | 42.725164 | 2.039864 | 1,325 |
| MON | striatum | 50 | 42.507878 | 2.122297 | 1,578 |
| PER | pseudomajus | 43 | 42.467675 | 2.8552415 | 296 |
| BOU | striatum | 20 | 42.643378 | 2.58705 | 209 |
| VIL | striatum | 50 | 42.587006 | 2.367453 | 441 |
| ARS | pseudomajus | 41 | 42.3895975 | 2.4876195 | 1,052 |
| THU | striatum | 47 | 42.644139 | 2.721694 | 123 |
| BAN | pseudomajus | 35 | 42.489458 | 3.124183 | 32 |
| ARL | pseudomajus | 21 | 42.4479485 | 2.6084845 | 329 |
| CIN | pseudomajus | 58 | 43.311569 | 1.533579 | 197 |
| YP1 | striatum | 50 | 42.326943 | 2.052929 | 1,343 |
| YP4 | striatum | 52 | 42.359921 | 1.926958 | 1,446 |
| MP4 | pseudomajus | 50 | 42.322234 | 2.091375 | 1,161 |
| MP11 | pseudomajus | 50 | 42.331038 | 2.170284 | 1,031 |
| M-AUT | pseudomajus | 1 | 43.342746 | 1.492514 | 220 |
| Z-ALE | striatum | 1 | 42.99 | 2.25 | 361 |
| Z-NDM | pseudomajus | 1 | 42.516039 | 2.976307 | 439 |
| Z-VCO | striatum | 1 | 42.59 | 2.37 | 427 |
| Ventolà | pseudomajus | 1 | 42.321803 | 2.137230 | 1,465 |
| Pont Napoléon | *Antirrhinum sempervirens* | 41 | 42.86 | -0.05 | 2,339 |

### **Supplementary Table 2: Names and genomic coordinates of partition islands in the *Antirrhinum majus* reference genome**

| **Supplementary Table 2: Names and genomic coordinates of partition islands.** | | | |
| --- | --- | --- | --- |
| **Island number** | **Island name** | **Coordinates** | **Partition** |
| 1 | CREMOSA (*CRE*) | Chr1:900,000-1,000,000 | A |
| 2 | AURINA (*AUN*) | Chr2:1,025,000-1,100,000 | A |
| 3 | FLAVIA (*FLA*) | Chr2:52,650,000-54,100,000 | A |
| 4 | SULFUREA (*SULF*) | Chr4:38,050,000-38,425,000 | A |
| 5 | RUBIA (*RUB*) | Chr5:6,250,000-6,300,000 | A |
| 6 | ROSEA / ELUTA (*ROS* *EL*) | Chr6:52,775,000-53,150,000 | A |
| 7 | XANTHIA (*XAT)* | Chr8:52,750,000-52,800,000 | A |
| 8 | ALTA1 (*ALT1*) | Chr6:42,025,000-42,100,000 | B |

### **Supplementary Table 3: SRB values and tree partitions given by 50 kb windows in partition islands**

| **Supplementary Table 3: SRB values and tree partitions given by 50 kb windows in partition islands.** | | | | | |
| --- | --- | --- | --- | --- | --- |
| **Chromosome** | **Partition island** | **Window start (Mb)** | **Window end (Mb)** | **SRB** | **Tree partition** |
| Chr1 | 1 | 0.9 | 0.95 | 0.00136 | A |
| Chr1 | 1 | 0.925 | 0.975 | 0.00327 | A |
| Chr1 | 1 | 0.95 | 1 | 0.00150 | A' |
| Chr2 | 2 | 1.025 | 1.075 | 0.00140 | A |
| Chr2 | 2 | 1.05 | 1.1 | 0.00069 | A |
| Chr2 | 3 | 46.525 | 46.575 | 0.00291 | A'' |
| Chr2 | 3 | 46.55 | 46.6 | 0.00126 | A'' |
| Chr2 | 3 | 52.65 | 52.7 | 0.00113 | A' |
| Chr2 | 3 | 52.825 | 52.875 | 0.00345 | A' |
| Chr2 | 3 | 52.85 | 52.9 | 0.00229 | A' |
| Chr2 | 3 | 52.875 | 52.925 | 0.00322 | A' |
| Chr2 | 3 | 52.9 | 52.95 | 0.00569 | A' |
| Chr2 | 3 | 52.925 | 52.975 | 0.00708 | A' |
| Chr2 | 3 | 52.95 | 53 | 0.00552 | A' |
| Chr2 | 3 | 52.975 | 53.025 | 0.00413 | A' |
| Chr2 | 3 | 53 | 53.05 | 0.00523 | A' |
| Chr2 | 3 | 53.025 | 53.075 | 0.00643 | A' |
| Chr2 | 3 | 53.05 | 53.1 | 0.00361 | A' |
| Chr2 | 3 | 53.1 | 53.15 | 0.00080 | A'' |
| Chr2 | 3 | 53.125 | 53.175 | 0.00226 | A' |
| Chr2 | 3 | 53.15 | 53.2 | 0.00515 | A' |
| Chr2 | 3 | 53.175 | 53.225 | 0.00636 | A' |
| Chr2 | 3 | 53.2 | 53.25 | 0.00475 | A' |
| Chr2 | 3 | 53.225 | 53.275 | 0.00252 | A'' |
| Chr2 | 3 | 53.25 | 53.3 | 0.00193 | A'' |
| Chr2 | 3 | 53.275 | 53.325 | 0.00504 | A' |
| Chr2 | 3 | 53.3 | 53.35 | 0.00791 | A' |
| Chr2 | 3 | 53.325 | 53.375 | 0.00621 | A' |
| Chr2 | 3 | 53.35 | 53.4 | 0.00432 | A' |
| Chr2 | 3 | 53.375 | 53.425 | 0.00607 | A' |
| Chr2 | 3 | 53.4 | 53.45 | 0.00812 | A' |
| Chr2 | 3 | 53.425 | 53.475 | 0.00739 | A' |
| Chr2 | 3 | 53.45 | 53.5 | 0.00695 | A |
| Chr2 | 3 | 53.475 | 53.525 | 0.00646 | A |
| Chr2 | 3 | 53.5 | 53.55 | 0.00495 | A |
| Chr2 | 3 | 53.525 | 53.575 | 0.00361 | A |
| Chr2 | 3 | 53.55 | 53.6 | 0.00434 | A' |
| Chr2 | 3 | 53.575 | 53.625 | 0.00386 | A' |
| Chr2 | 3 | 53.6 | 53.65 | 0.00326 | A' |
| Chr2 | 3 | 53.625 | 53.675 | 0.00377 | A |
| Chr2 | 3 | 53.65 | 53.7 | 0.00514 | A |
| Chr2 | 3 | 53.675 | 53.725 | 0.00450 | A |
| Chr2 | 3 | 53.7 | 53.75 | 0.00335 | A |
| Chr2 | 3 | 53.725 | 53.775 | 0.00284 | A |
| Chr2 | 3 | 53.75 | 53.8 | 0.00282 | A |
| Chr2 | 3 | 53.775 | 53.825 | 0.00155 | A |
| Chr2 | 3 | 53.825 | 53.875 | 0.00327 | A'' |
| Chr2 | 3 | 53.85 | 53.9 | 0.00316 | A'' |
| Chr2 | 3 | 53.875 | 53.925 | 0.00173 | A'' |
| Chr2 | 3 | 53.9 | 53.95 | 0.00346 | A' |
| Chr2 | 3 | 53.925 | 53.975 | 0.00247 | A' |
| Chr2 | 3 | 53.95 | 54 | 0.00256 | A' |
| Chr2 | 3 | 53.975 | 54.025 | 0.00307 | A' |
| Chr2 | 3 | 54 | 54.05 | 0.00297 | A |
| Chr2 | 3 | 54.025 | 54.075 | 0.00290 | A |
| Chr2 | 3 | 54.05 | 54.1 | 0.00213 | A |
| Chr2 | 3 | 54.225 | 54.275 | 0.00041 | A' |
| Chr2 | 3 | 54.25 | 54.3 | 0.00035 | A' |
| Chr4 | 4 | 38.05 | 38.1 | 0.00300 | A |
| Chr4 | 4 | 38.075 | 38.125 | 0.00546 | A' |
| Chr4 | 4 | 38.1 | 38.15 | 0.00371 | A'' |
| Chr4 | 4 | 38.125 | 38.175 | 0.00427 | A'' |
| Chr4 | 4 | 38.35 | 38.4 | 0.00112 | A'' |
| Chr4 | 4 | 38.375 | 38.425 | 0.00048 | A'' |
| Chr4 | 4 | 38.4 | 38.45 | 0.00095 | A'' |
| Chr4 | 4 | 38.425 | 38.475 | 0.00124 | A'' |
| Chr5 | 5 | 6.25 | 6.3 | 0.00237 | A' |
| Chr6 | 6 | 52.775 | 52.825 | 0.00078 | A' |
| Chr6 | 6 | 52.8 | 52.85 | 0.00062 | A'' |
| Chr6 | 6 | 52.825 | 52.875 | 0.00087 | A |
| Chr6 | 6 | 52.85 | 52.9 | 0.00383 | A |
| Chr6 | 6 | 52.875 | 52.925 | 0.00400 | A |
| Chr6 | 6 | 52.9 | 52.95 | 0.00244 | A |
| Chr6 | 6 | 52.925 | 52.975 | 0.00059 | A |
| Chr6 | 6 | 52.975 | 53.025 | 0.00105 | A |
| Chr6 | 6 | 53 | 53.05 | 0.00187 | A |
| Chr6 | 6 | 53.025 | 53.075 | 0.00418 | A |
| Chr6 | 6 | 53.05 | 53.1 | 0.00265 | A |
| Chr6 | 6 | 53.075 | 53.125 | 0.00076 | A |
| Chr6 | 6 | 53.1 | 53.15 | 0.00047 | A |
| Chr8 | 7 | 52.75 | 52.8 | 0.00062 | A' |
| Chr6 | 8 | 42.025 | 42.075 | 0.00262 | B |
| Chr6 | 8 | 42.05 | 42.1 | 0.00407 | B |

### **Supplementary Table 4: Gene density across the genome, compared to partition islands**

| **Supplementary Table 4: Gene density across the genome, compared to partition islands.** | | | |
| --- | --- | --- | --- |
| **Locus** | **Sequence length** | **Predicted coding sequences** | **Coding sequences / 1 Mb** |
| Chromosome 1 | 71,919,034 | 7,005 | 97.40119702 |
| Chromosome 2 | 77,118,269 | 6,545 | 84.86964353 |
| Chromosome 3 | 65,231,163 | 5,286 | 81.03488819 |
| Chromosome 4 | 54,887,108 | 4,871 | 88.74579437 |
| Chromosome 5 | 71,106,538 | 5,569 | 78.31909915 |
| Chromosome 6 | 55,699,338 | 4,623 | 82.99919112 |
| Chromosome 7 | 55,564,713 | 4,617 | 83.09230356 |
| Chromosome 8 | 57,431,585 | 4,551 | 79.24211042 |
| Whole genome | 508,957,748 | 43,067 | 84.61802609 |
| Partition island 1 | 100,000 | 9 | 90 |
| Partition island 2 | 75,000 | 13 | 173.3333333 |
| Partition island 3 | 1,450,000 | 146 | 100.6896552 |
| Partition island 4 | 375,000 | 32 | 85.33333333 |
| Partition island 5 | 50,000 | 6 | 120 |
| Partition island 6 | 75,000 | 9 | 120 |
| Partition island 7 | 375,000 | 43 | 114.6666667 |
| Partition island 8 | 50,000 | 5 | 100 |
| All islands | 2,550,000 | 263 | 103.1372549 |

### **Supplementary Table 5: eggnog-mapper functional predictions for coding sequences within partition islands in the A. majus reference genome.**

| **Supplementary Table 5: *eggnog-mapper* functional predictions for coding sequences within partition islands in the *A. majus* reference genome.** | | | | |
| --- | --- | --- | --- | --- |
| **CDS ID** | **Partition Island** | **e-Value** | **Score** | **Description** |
| AnM01G00094.01 | 1 | 4.17E-201 | 586 | Modified RING finger domain |
| AnM01G00094.02 | 1 | 1.09E-204 | 595 | Modified RING finger domain |
| AnM01G00095.01 | 1 | 3.65E-74 | 229 | Domain of unknown function (DUF4228) |
| AnM01G00096.01 | 1 | 0 | 1800 | Phosphatidylinositol-4-phosphate 5-Kinase |
| AnM01G00097.01 | 1 | 3.22E-191 | 538 | Nicotinate-nucleotide pyrophosphorylase carboxylating |
| AnM01G00098.01 | 1 | 1.41E-164 | 471 | Belongs to the class I-like SAM-binding methyltransferase superfamily. Cation-independent O- methyltransferase family |
| AnM01G00099.01 | 1 | 0 | 1145 | Epidermal growth factor-like domain. |
| AnM01G00100.01 | 1 |  |  |  |
| AnM01G00101.01 | 1 | 0 | 2131 | DNA-directed RNA polymerase |
| AnM01G00102.01 | 1 | 4.21E-14 | 70.9 | leucine-rich repeat extensin-like protein |
| AnM01G07125.01 | 2 | 1.60E-27 | 108 | Plant invertase/pectin methylesterase inhibitor |
| AnM01G07126.01 | 2 | 1.51E-229 | 649 | Belongs to the protein kinase superfamily. Ser Thr protein kinase family |
| AnM01G07127.01 | 2 | 2.50E-167 | 479 | Toprim domain |
| AnM01G07127.02 | 2 | 9.23E-165 | 473 | Toprim domain |
| AnM01G07128.01 | 2 | 2.02E-237 | 667 | CRAL/TRIO, N-terminal domain |
| AnM01G07129.01 | 2 |  |  |  |
| AnM01G07130.01 | 2 | 5.92E-239 | 677 | Polyphenol oxidase, chloroplastic-like |
| AnM01G07131.01 | 2 | 2.53E-238 | 673 | Belongs to the oxygen-dependent FAD-linked oxidoreductase family |
| AnM01G07132.01 | 2 | 3.76E-100 | 310 | Belongs to the oxygen-dependent FAD-linked oxidoreductase family |
| AnM01G07133.01 | 2 | 1.74E-237 | 667 | Belongs to the oxygen-dependent FAD-linked oxidoreductase family |
| AnM01G07134.01 | 2 | 8.86E-301 | 830 | Belongs to the oxygen-dependent FAD-linked oxidoreductase family |
| AnM01G07135.01 | 2 | 4.18E-284 | 788 | Belongs to the oxygen-dependent FAD-linked oxidoreductase family |
| AnM01G07136.01 | 2 | 1.15E-289 | 802 | Belongs to the oxygen-dependent FAD-linked oxidoreductase family |
| AnM01G07137.01 | 2 | 8.49E-236 | 665 | flavin adenine dinucleotide binding |
| AnM01G10994.01 | 3 | 2.64E-61 | 194 | methyl-CpG-binding domain-containing protein |
| AnM01G10994.02 | 3 | 3.74E-61 | 194 | methyl-CpG-binding domain-containing protein |
| AnM01G10995.01 | 3 | 0 | 3693 | Piezo non-specific cation channel, R-Ras-binding domain |
| AnM01G10996.01 | 3 | 1.11E-97 | 290 | - |
| AnM01G10997.01 | 3 |  |  |  |
| AnM01G10998.01 | 3 | 2.69E-59 | 188 | Auxin binding protein |
| AnM01G10999.01 | 3 |  |  |  |
| AnM01G11000.01 | 3 | 5.52E-205 | 572 | Belongs to the glycosyltransferase 8 family |
| AnM01G11001.01 | 3 |  |  |  |
| AnM01G11002.01 | 3 | 3.56E-236 | 666 | oxidoreductase, 2OG-Fe(II) oxygenase family protein |
| AnM01G11003.01 | 3 |  |  |  |
| AnM01G11004.01 | 3 |  |  |  |
| AnM01G11005.01 | 3 |  |  |  |
| AnM01G11006.01 | 3 | 1.85E-183 | 520 | Belongs to the 'GDSL' lipolytic enzyme family |
| AnM01G11007.01 | 3 |  |  |  |
| AnM01G11008.01 | 3 | 1.87E-176 | 504 | Belongs to the 'GDSL' lipolytic enzyme family |
| AnM01G11009.01 | 3 | 6.65E-50 | 189 | acid phosphatase activity |
| AnM01G11010.01 | 3 | 9.12E-291 | 797 | This protein promotes the GTP-dependent binding of aminoacyl-tRNA to the A-site of ribosomes during protein biosynthesis |
| AnM01G11011.01 | 3 | 2.73E-94 | 295 | - |
| AnM01G11012.01 | 3 | 2.14E-59 | 203 | - |
| AnM01G11013.01 | 3 | 1.06E-62 | 191 | Component of LSM protein complexes, which are involved in RNA processing |
| AnM01G11014.01 | 3 | 6.78E-32 | 121 | Natural resistance-associated macrophage protein |
| AnM01G11015.01 | 3 | 1.15E-148 | 421 | Dienelactone hydrolase family |
| AnM01G11016.01 | 3 | 5.09E-130 | 417 | cysteine-rich RLK (RECEPTOR-like protein kinase) 8 |
| AnM01G11017.01 | 3 | 7.25E-29 | 115 | gag-polypeptide of LTR copia-type |
| AnM01G11018.01 | 3 | 4.99E-51 | 179 | Ribonuclease H protein |
| AnM01G11019.01 | 3 |  |  |  |
| AnM01G11020.01 | 3 |  |  |  |
| AnM01G11021.01 | 3 |  |  |  |
| AnM01G11022.01 | 3 | 0 | 2013 | mediator of RNA polymerase II transcription subunit |
| AnM01G11022.02 | 3 | 0 | 2109 | mediator of RNA polymerase II transcription subunit |
| AnM01G11023.01 | 3 | 0 | 905 | Cytosolic domain of 10TM putative phosphate transporter |
| AnM01G11024.01 | 3 |  |  |  |
| AnM01G11025.01 | 3 | 5.13E-123 | 395 | strictosidine synthase activity |
| AnM01G11026.01 | 3 | 0 | 1309 | RPR |
| AnM01G11027.01 | 3 | 3.02E-101 | 295 | Belongs to the glutathione peroxidase family |
| AnM01G11028.01 | 3 | 5.71E-257 | 715 | Domain of unknown function DUF21 |
| AnM01G11029.01 | 3 | 4.27E-21 | 95.1 | Belongs to the GRAS family |
| AnM01G11030.01 | 3 | 8.31E-45 | 163 | Belongs to the cyclin family |
| AnM01G11030.02 | 3 | 3.26E-48 | 172 | Belongs to the cyclin family |
| AnM01G11031.01 | 3 | 2.23E-17 | 84.3 | Putative S-adenosyl-L-methionine-dependent methyltransferase |
| AnM01G11032.01 | 3 |  |  |  |
| AnM01G11033.01 | 3 | 9.25E-44 | 149 | Plant lipid transfer protein / seed storage protein / trypsin-alpha amylase inhibitor domain family |
| AnM01G11034.01 | 3 | 6.81E-136 | 394 | Adenylate isopentenyltransferase |
| AnM01G11035.01 | 3 | 1.46E-57 | 193 | transcription factor |
| AnM01G11035.02 | 3 | 4.55E-59 | 197 | transcription factor |
| AnM01G11036.01 | 3 | 5.91E-161 | 456 | Phenazine biosynthesis-like domain-containing protein 1 |
| AnM01G11037.01 | 3 | 1.71E-142 | 419 | Phenazine biosynthesis-like domain-containing protein 1 |
| AnM01G11038.01 | 3 | 9.98E-246 | 682 | Belongs to the peptidase A1 family |
| AnM01G11039.01 | 3 | 8.62E-240 | 667 | Belongs to the peptidase A1 family |
| AnM01G11040.01 | 3 | 7.37E-33 | 119 | Salt stress response/antifungal |
| AnM01G11041.01 | 3 | 3.14E-11 | 71.6 | iq-domain |
| AnM01G11042.01 | 3 | 1.17E-63 | 197 | Salt stress response/antifungal |
| AnM01G11043.01 | 3 |  |  |  |
| AnM01G11044.01 | 3 | 6.85E-226 | 650 | basic helix-loop-helix protein |
| AnM01G11045.01 | 3 | 0 | 1431 | Helical and beta-bridge domain |
| AnM01G11046.01 | 3 | 2.77E-121 | 367 | Transferase family |
| AnM01G11047.01 | 3 |  |  |  |
| AnM01G11048.01 | 3 | 1.62E-139 | 411 | Transferase family |
| AnM01G11049.01 | 3 |  |  |  |
| AnM01G11050.01 | 3 | 4.80E-200 | 570 | Transferase family |
| AnM01G11051.01 | 3 | 2.73E-15 | 77 | mitochondrial protein |
| AnM01G11052.01 | 3 | 3.74E-17 | 85.5 | mitochondrial protein |
| AnM01G11053.01 | 3 |  |  |  |
| AnM01G11054.01 | 3 |  |  |  |
| AnM01G11055.01 | 3 | 9.37E-10 | 60.8 | transposition, RNA-mediated |
| AnM01G11056.01 | 3 | 1.97E-22 | 104 | Uncharacterized protein K02A2.6-like |
| AnM01G11057.01 | 3 | 1.16E-49 | 181 | Protein of unknown function (DUF 659) |
| AnM01G11058.01 | 3 | 4.89E-36 | 147 | Protein of unknown function (DUF 659) |
| AnM01G11059.01 | 3 |  |  |  |
| AnM01G11060.01 | 3 |  |  |  |
| AnM01G11061.01 | 3 | 2.61E-43 | 152 | - |
| AnM01G11062.01 | 3 | 0 | 1085 | Mitochondrial Rho GTPase |
| AnM01G11063.01 | 3 | 3.00E-45 | 160 | Fantastic Four meristem regulator |
| AnM01G11064.01 | 3 | 2.24E-08 | 61.6 | f-box protein |
| AnM01G11065.01 | 3 |  |  |  |
| AnM01G11066.01 | 3 | 4.50E-23 | 103 | isoform X1 |
| AnM01G11066.02 | 3 | 4.77E-23 | 103 | isoform X1 |
| AnM01G11067.01 | 3 |  |  |  |
| AnM01G11068.01 | 3 | 4.74E-191 | 546 | Belongs to the UDP-glycosyltransferase family |
| AnM01G11069.01 | 3 | 6.34E-11 | 68.9 | isoform X1 |
| AnM01G11070.01 | 3 | 1.54E-124 | 368 | - |
| AnM01G11071.01 | 3 | 2.85E-225 | 628 | Chloroplast stem-loop binding protein of 41 kDa a, chloroplastic |
| AnM01G11072.01 | 3 | 1.23E-190 | 552 | RAN GTPase-activating protein |
| AnM01G11073.01 | 3 |  |  |  |
| AnM01G11074.01 | 3 | 1.68E-71 | 218 | DnaJ molecular chaperone homology domain |
| AnM01G11075.01 | 3 | 1.34E-213 | 595 | Ferrous iron transport protein B |
| AnM01G11076.01 | 3 |  |  |  |
| AnM01G11077.01 | 3 | 3.49E-15 | 77.8 | transposition, RNA-mediated |
| AnM01G11078.01 | 3 | 4.75E-40 | 147 | Contains the following InterPro domains Glycoside hydrolase, family 18, catalytic domain (InterPro IPR001223), Chitinase II (InterPro IPR011583), Glycoside hydrolase, catalytic core (InterPro IPR017853), Glycoside hydrolase, subgroup, catalytic core (InterPro IPR013781) |
| AnM01G11079.01 | 3 | 6.52E-53 | 184 | transposition, RNA-mediated |
| AnM01G11080.01 | 3 | 2.55E-73 | 244 | Uncharacterized protein K02A2.6-like |
| AnM01G11081.01 | 3 |  |  |  |
| AnM01G11082.01 | 3 |  |  |  |
| AnM01G11083.01 | 3 | 1.86E-235 | 652 | Lipolytic acyl hydrolase (LAH) |
| AnM01G11084.01 | 3 | 1.92E-61 | 196 | Plastocyanin-like domain |
| AnM01G11085.01 | 3 | 3.22E-131 | 380 | Belongs to the peroxidase family. Classical plant (class III) peroxidase subfamily |
| AnM01G11086.01 | 3 |  |  |  |
| AnM01G11087.01 | 3 |  |  |  |
| AnM01G11087.02 | 3 |  |  |  |
| AnM01G11088.01 | 3 |  |  |  |
| AnM01G11089.01 | 3 |  |  |  |
| AnM01G11090.01 | 3 | 7.46E-11 | 69.3 | transposition, RNA-mediated |
| AnM01G11091.01 | 3 | 6.66E-49 | 177 | Uncharacterized protein K02A2.6-like |
| AnM01G11092.01 | 3 | 1.13E-67 | 240 | transposition, RNA-mediated |
| AnM01G11093.01 | 3 |  |  |  |
| AnM01G11094.01 | 3 |  |  |  |
| AnM01G11095.01 | 3 |  |  |  |
| AnM01G11096.01 | 3 |  |  |  |
| AnM01G11097.01 | 3 |  |  |  |
| AnM01G11098.01 | 3 |  |  |  |
| AnM01G11099.01 | 3 | 0 | 1121 | Belongs to the terpene synthase family |
| AnM01G11100.01 | 3 |  |  |  |
| AnM01G11101.01 | 3 | 3.06E-34 | 130 | Uncharacterized protein K02A2.6-like |
| AnM01G11102.01 | 3 | 7.87E-77 | 263 | transposition, RNA-mediated |
| AnM01G11103.01 | 3 |  |  |  |
| AnM01G11104.01 | 3 | 1.94E-231 | 645 | Splicing Factor Motif, present in Prp18 and Pr04 |
| AnM01G11105.01 | 3 | 7.03E-67 | 208 | - |
| AnM01G11106.01 | 3 | 1.12E-123 | 364 | Four repeated domains in the Fasciclin I family of proteins, present in many other contexts. |
| AnM01G11107.01 | 3 | 1.85E-10 | 64.3 |  |
| AnM01G11108.01 | 3 |  |  |  |
| AnM01G11109.01 | 3 |  |  |  |
| AnM01G11110.01 | 3 |  |  |  |
| AnM01G11111.01 | 3 | 4.27E-33 | 130 | fasciclin-like arabinogalactan protein |
| AnM01G11112.01 | 3 | 3.10E-41 | 151 | gag-polypeptide of LTR copia-type |
| AnM01G11113.01 | 3 | 2.04E-33 | 130 | fasciclin-like arabinogalactan protein |
| AnM01G11114.01 | 3 | 1.86E-24 | 100 | Reverse transcriptase-like |
| AnM01G11115.01 | 3 | 1.16E-19 | 90.1 | Four repeated domains in the Fasciclin I family of proteins, present in many other contexts. |
| AnM01G11116.01 | 3 | 3.74E-20 | 91.3 | Ribonuclease H protein |
| AnM01G11117.01 | 3 |  |  |  |
| AnM01G11118.01 | 3 | 4.92E-66 | 221 | fasciclin-like arabinogalactan protein |
| AnM01G11119.01 | 3 | 5.68E-134 | 389 | Plant protein of unknown function (DUF868) |
| AnM01G11120.01 | 3 | 7.76E-66 | 204 | Belongs to the eukaryotic ribosomal protein eS24 family |
| AnM01G11121.01 | 3 | 4.02E-104 | 301 | Ribosomal protein S13/S18 |
| AnM01G11122.01 | 3 | 1.04E-38 | 149 | Ribonuclease H protein |
| AnM01G11123.01 | 3 | 6.52E-35 | 138 | Domain of unknown function (DUF4283) |
| AnM01G11124.01 | 3 | 6.04E-57 | 186 | transcription repressor |
| AnM01G11125.01 | 3 | 8.97E-39 | 146 | Domain of unknown function (DUF4216) |
| AnM01G11126.01 | 3 | 4.14E-118 | 345 | late embryogenesis abundant protein |
| AnM01G11127.01 | 3 |  |  |  |
| AnM01G11128.01 | 3 |  |  |  |
| AnM01G11129.01 | 3 | 2.92E-108 | 320 | late embryogenesis abundant protein |
| AnM01G11130.01 | 3 | 2.15E-122 | 355 | late embryogenesis abundant protein |
| AnM01G11131.01 | 3 | 3.19E-123 | 357 | late embryogenesis abundant protein |
| AnM01G11132.01 | 3 |  |  |  |
| AnM01G11133.01 | 3 |  |  |  |
| AnM01G11134.01 | 3 |  |  |  |
| AnM01G11135.01 | 3 | 5.00E-59 | 191 | Proline-rich nuclear receptor coactivator motif |
| AnM01G11136.01 | 3 |  |  |  |
| AnM01G11137.01 | 3 | 1.45E-68 | 220 | zinc-finger of the FCS-type, C2-C2 |
| AnM01G11138.01 | 3 |  |  |  |
| AnM01G11139.01 | 3 |  |  |  |
| AnM01G27070.01 | 4 | 1.31E-129 | 375 | PPPDE putative peptidase domain |
| AnM01G27071.01 | 4 | 8.78E-62 | 198 | - |
| AnM01G27072.01 | 4 |  |  |  |
| AnM01G27073.01 | 4 | 3.63E-168 | 476 | Belongs to the sterol desaturase family |
| AnM01G27074.01 | 4 | 2.98E-44 | 175 | Tetratricopeptide repeat |
| AnM01G27075.01 | 4 | 0 | 1220 | Heat shock protein |
| AnM01G27076.01 | 4 | 6.88E-102 | 297 | 40S ribosomal protein |
| AnM01G27077.01 | 4 | 1.91E-73 | 238 | WRKY transcription factor |
| AnM01G27078.01 | 4 | 7.87E-192 | 540 | SGS domain |
| AnM01G27079.01 | 4 | 9.17E-116 | 332 | Sybindin-like family |
| AnM01G27080.01 | 4 | 3.33E-75 | 238 | - |
| AnM01G27081.01 | 4 | 2.38E-33 | 126 | - |
| AnM01G27082.01 | 4 | 3.57E-10 | 69.3 | Uncharacterized protein K02A2.6-like |
| AnM01G27083.01 | 4 | 4.91E-67 | 238 | Plant mobile domain |
| AnM01G27084.01 | 4 | 1.54E-203 | 577 | Belongs to the UDP-glycosyltransferase family |
| AnM01G27085.01 | 4 | 9.55E-48 | 172 | gag-polypeptide of LTR copia-type |
| AnM01G27086.01 | 4 | 2.98E-57 | 183 | Pathogenesis-related protein Bet v I family |
| AnM01G27087.01 | 4 | 8.71E-201 | 570 | Belongs to the UDP-glycosyltransferase family |
| AnM01G27088.01 | 4 | 9.55E-48 | 172 | gag-polypeptide of LTR copia-type |
| AnM01G27089.01 | 4 | 3.47E-13 | 69.3 | Major allergen Pru ar 1-like |
| AnM01G27090.01 | 4 | 8.71E-201 | 570 | Belongs to the UDP-glycosyltransferase family |
| AnM01G27091.01 | 4 | 9.55E-48 | 172 | gag-polypeptide of LTR copia-type |
| AnM01G27092.01 | 4 | 2.98E-57 | 183 | Pathogenesis-related protein Bet v I family |
| AnM01G27093.01 | 4 | 6.57E-169 | 488 | Belongs to the UDP-glycosyltransferase family |
| AnM01G27094.01 | 4 | 7.25E-54 | 183 | Belongs to the UDP-glycosyltransferase family |
| AnM01G27095.01 | 4 | 1.08E-16 | 78.2 | Ribonuclease H protein |
| AnM01G27096.01 | 4 | 1.73E-11 | 72.8 | Ribonuclease H protein |
| AnM01G27097.01 | 4 |  |  |  |
| AnM01G27098.01 | 4 | 0 | 1321 | Bulb-type mannose-specific lectin |
| AnM01G27099.01 | 4 | 0 | 1415 | Timeless protein |
| AnM01G27100.01 | 4 | 0 | 972 | Belongs to the pyruvate kinase family |
| AnM01G27101.01 | 4 | 4.78E-79 | 270 | - |
| AnM01G19541.01 | 5 | 4.82E-161 | 483 | Protein KRI1 homolog |
| AnM01G19542.01 | 5 | 1.14E-42 | 142 | Gibberellin-regulated protein |
| AnM01G19543.01 | 5 | 3.63E-209 | 582 | Belongs to the iron ascorbate-dependent oxidoreductase family |
| AnM01G19544.01 | 5 | 0 | 875 | Transmembrane amino acid transporter protein |
| AnM01G19545.01 | 5 | 6.14E-174 | 488 | Thaumatin family |
| AnM01G19546.01 | 5 | 3.25E-229 | 645 | dnaJ homolog 1, mitochondrial-like |
| AnM01G33517.01 | 6 |  |  |  |
| AnM01G33518.01 | 6 |  |  |  |
| AnM01G33519.01 | 6 |  |  |  |
| AnM01G33520.01 | 6 | 1.52E-217 | 628 | Carbohydrate-binding protein of the ER |
| AnM01G33521.01 | 6 | 1.72E-49 | 157 | protein complex oligomerization |
| AnM01G33522.01 | 6 | 2.20E-36 | 143 | Oligopeptidase |
| AnM01G33523.01 | 6 |  |  |  |
| AnM01G33524.01 | 6 | 5.00E-36 | 125 | - |
| AnM01G33525.01 | 6 | 3.87E-92 | 279 | MADS-box transcription factor |
| AnM01G33525.02 | 6 | 6.97E-96 | 288 | MADS-box transcription factor |
| AnM01G33526.01 | 6 | 8.85E-56 | 178 | Uncharacterized protein At4g22758-like |
| AnM01G33527.01 | 6 | 8.73E-111 | 328 | Glycogen recognition site of AMP-activated protein kinase |
| AnM01G33528.01 | 6 | 1.66E-69 | 220 | Myb-like DNA-binding domain |
| AnM01G33529.01 | 6 | 7.32E-46 | 153 | Transcription factor |
| AnM01G33530.01 | 6 | 1.17E-130 | 410 | Belongs to the helicase family |
| AnM01G33531.01 | 6 | 2.10E-09 | 60.8 | Ribonuclease H protein |
| AnM01G33532.01 | 6 | 5.88E-94 | 320 | ribonuclease H protein |
| AnM01G33533.01 | 6 | 2.92E-67 | 215 | Myb-like DNA-binding domain |
| AnM01G33534.01 | 6 | 4.07E-25 | 100 | MLP-like protein |
| AnM01G33535.01 | 6 | 5.42E-17 | 80.1 | MLP-like protein |
| AnM01G33536.01 | 6 | 1.33E-25 | 107 | MLP-like protein |
| AnM01G33537.01 | 6 |  |  |  |
| AnM01G33538.01 | 6 | 3.45E-139 | 399 | Eukaryotic translation initiation factor 2 subunit |
| AnM01G33539.01 | 6 | 3.80E-146 | 425 | GDSL-like Lipase/Acylhydrolase |
| AnM01G33540.01 | 6 | 3.12E-29 | 111 | MLP-like protein |
| AnM01G33541.01 | 6 | 1.19E-25 | 102 | EF-hand domain |
| AnM01G33542.01 | 6 | 1.02E-92 | 285 | helix loop helix domain |
| AnM01G33543.01 | 6 | 2.08E-144 | 421 | U-box domain-containing protein |
| AnM01G33544.01 | 6 | 0 | 1907 | Protease Do-like 7 |
| AnM01G33545.01 | 6 |  |  |  |
| AnM01G33546.01 | 6 | 3.24E-34 | 125 | - |
| AnM01G33547.01 | 6 | 1.21E-37 | 134 | Domain of unknown function (DUF4228) |
| AnM01G33548.01 | 6 | 3.76E-16 | 77.4 | - |
| AnM01G33549.01 | 6 |  |  |  |
| AnM01G33550.01 | 6 |  |  |  |
| AnM01G33551.01 | 6 | 2.85E-65 | 207 | SANT SWI3, ADA2, N-CoR and TFIIIB'' DNA-binding domains |
| AnM01G33552.01 | 6 | 3.47E-222 | 620 | AAR2 protein |
| AnM01G33553.01 | 6 | 9.98E-80 | 239 | Heavy metal-associated isoprenylated plant protein |
| AnM01G33554.01 | 6 | 0 | 1050 | high mobility group |
| AnM01G33555.01 | 6 | 6.43E-123 | 358 | Cofactor assembly of complex C |
| AnM01G33556.01 | 6 | 3.88E-45 | 168 | E3 ubiquitin-protein ligase that mediates ubiquitination and subsequent proteasomal degradation of target proteins. E3 ubiquitin ligases accept ubiquitin from an E2 ubiquitin- conjugating enzyme in the form of a thioester and then directly transfers the ubiquitin to targeted substrates |
| AnM01G33557.01 | 6 | 1.28E-19 | 95.5 | regulation of N-terminal protein palmitoylation |
| AnM01G33558.01 | 6 |  |  |  |
| AnM01G33559.01 | 6 | 1.39E-71 | 234 | E3 ubiquitin-protein ligase that mediates ubiquitination and subsequent proteasomal degradation of target proteins. E3 ubiquitin ligases accept ubiquitin from an E2 ubiquitin- conjugating enzyme in the form of a thioester and then directly transfers the ubiquitin to targeted substrates |
| AnM01G42377.01 | 7 | 0 | 889 | starch synthase |
| AnM01G42378.01 | 7 | 3.91E-306 | 850 | YT521-B-like domain |
| AnM01G42379.01 | 7 | 9.01E-34 | 120 | Cytochrome c oxidase subunit VII |
| AnM01G42380.01 | 7 | 2.79E-93 | 295 | Squamosa promoter-binding-like protein |
| AnM01G42381.01 | 7 | 7.75E-209 | 589 | CemA family |
| AnM01G42382.01 | 7 | 7.20E-220 | 620 | Sugar (and other) transporter |
| AnM01G42383.01 | 7 | 1.84E-251 | 701 | Sugar (and other) transporter |
| AnM01G32329.01 | 8 | 0 | 986 | Carbohydrate binding domain CBM49 |
| AnM01G32330.01 | 8 |  |  |  |
| AnM01G32331.01 | 8 | 3.40E-18 | 81.6 | protection from non-homologous end joining at telomere |
| AnM01G32332.01 | 8 | 0 | 936 | Belongs to the aldehyde dehydrogenase family |
| AnM01G32333.01 | 8 |  |  |  |
| AnM01G32334.01 | 8 | 5.06E-115 | 358 | Transposase family tnp2 |
| AnM01G32335.01 | 8 | 8.98E-42 | 144 | TdcA1-ORF2 protein |
| AnM01G32336.01 | 8 |  |  |  |
| AnM01G32337.01 | 8 | 1.59E-216 | 613 | Belongs to the cytochrome P450 family |

### **Supplementary Table 6: *eggnog-mapper* functional predictions for differentially expressed coding sequences within partition islands in the *A. m. m.* var*. pseudomajus* assembly**

| **Supplementary Table 6: *eggnog-mapper* functional predictions for differentially expressed coding sequences within partition islands in the *A. m. m.* var*. pseudomajus* assembly.** | | | | |
| --- | --- | --- | --- | --- |
| **CDS ID** | **Partition Island** | **e-Value** | **Score** | **Description** |
| A.pmajus012565.01 | 1 | 1.80E-18 | 82.8 | leucine-rich repeat extensin-like protein |
| A.pmajus012569.01 | 1 | 1.08E-96 | 292 | Belongs to the class I-like SAM-binding methyltransferase superfamily. Cation-independent O- methyltransferase family |
| A.pmajus012570.01 | 1 | 1.97E-52 | 176 | Belongs to the class I-like SAM-binding methyltransferase superfamily. Cation-independent O- methyltransferase family |
| A.pmajus012571.01 | 1 | 1.15E-163 | 468 | Belongs to the class I-like SAM-binding methyltransferase superfamily. Cation-independent O- methyltransferase family |
| A.pmajus012572.01 | 1 | 2.10E-192 | 539 | Nicotinate-nucleotide pyrophosphorylase carboxylating |
| A.pmajus034314.01 | 1 | 1.20E-57 | 194 | Transferase family |
| A.pmajus007962.01 | 2 | 4.44E-241 | 682 | Polyphenol oxidase, chloroplastic-like |
| A.pmajus007963.01 | 2 | 2.54E-250 | 707 | Polyphenol oxidase, chloroplastic-like |
| A.pmajus007964.01 | 2 | 4.06E-233 | 660 | Belongs to the oxygen-dependent FAD-linked oxidoreductase family |
| A.pmajus007969.01 | 2 | 6.95E-291 | 805 | Belongs to the oxygen-dependent FAD-linked oxidoreductase family |
| A.pmajus024142.01 | 2 | 3.33E-286 | 794 | Belongs to the oxygen-dependent FAD-linked oxidoreductase family |
| A.pmajus001506.01 | 3 | 2.76E-11 | 70.5 | zinc-binding in reverse transcriptase |
| A.pmajus000172.01 | 3 | 4.35E-118 | 351 | Belongs to the class I-like SAM-binding methyltransferase superfamily. Cation-independent O- methyltransferase family |
| A.pmajus000510.01 | 3 | 0 | 1292 | Prolyl oligopeptidase, N-terminal beta-propeller domain |
| A.pmajus006541.01 | 3 |  |  |  |
| A.pmajus012921.01 | 3 | 1.08E-59 | 194 | Catalyzes xyloglucan endohydrolysis (XEH) and or endotransglycosylation (XET). Cleaves and religates xyloglucan polymers, an essential constituent of the primary cell wall, and thereby participates in cell wall construction of growing tissues |
| A.pmajus009207.01 | 3 | 3.25E-130 | 382 | Zinc finger, C3HC4 type (RING finger) |
| A.pmajus018062.01 | 3 | 1.22E-50 | 171 | synthase |
| A.pmajus030889.01 | 3 |  |  |  |
| A.pmajus023541.01 | 3 | 0 | 1725 | Belongs to the glycosyltransferase 2 family. Plant cellulose synthase subfamily |
| A.pmajus026155.01 | 3 |  |  |  |
| A.pmajus013053.01 | 3 | 0 | 1608 | Belongs to the glycosyl hydrolase 31 family |
| A.pmajus031326.01 | 3 | 6.64E-108 | 319 | late embryogenesis abundant protein |
| A.pmajus031337.01 | 3 | 1.88E-133 | 389 | Plant protein of unknown function (DUF868) |
| A.pmajus031369.01 | 3 | 2.64E-235 | 651 | Lipolytic acyl hydrolase (LAH) |
| A.pmajus031377.01 | 3 | 3.46E-225 | 628 | Chloroplast stem-loop binding protein of 41 kDa a, chloroplastic |
| A.pmajus031378.01 | 3 | 3.37E-119 | 354 | - |
| A.pmajus031382.01 | 3 | 6.18E-194 | 553 | Belongs to the UDP-glycosyltransferase family |
| A.pmajus031383.01 | 3 | 5.59E-23 | 103 | isoform X1 |
| A.pmajus031383.02 | 3 | 5.83E-23 | 103 | isoform X1 |
| A.pmajus031388.01 | 3 | 1.01E-45 | 162 | Fantastic Four meristem regulator |
| A.pmajus038185.01 | 3 | 1.02E-74 | 253 | transposition, RNA-mediated |
| A.pmajus034231.01 | 3 | 4.39E-14 | 79.3 | zinc-binding in reverse transcriptase |
| A.pmajus036168.01 | 3 | 7.58E-238 | 670 | oxidoreductase, 2OG-Fe(II) oxygenase family protein |
| A.pmajus040272.01 | 3 | 6.30E-215 | 612 | OTU domain-containing protein |
| A.pmajus040272.02 | 3 | 2.47E-215 | 613 | OTU domain-containing protein |
| A.pmajus041562.01 | 3 | 1.85E-229 | 659 | basic helix-loop-helix protein |
| A.pmajus041571.01 | 3 | 1.69E-160 | 455 | Phenazine biosynthesis-like domain-containing protein 1 |
| A.pmajus031349.01 | 3 |  |  |  |
| A.pmajus025506.01 | 3 | 1.24E-199 | 569 | Belongs to the peptidase S10 family |
| A.pmajus005557.01 | 3 | 7.75E-33 | 137 | Retrotransposon gag protein |
| A.pmajus015427.01 | 3 |  |  |  |
| A.pmajus001398.01 | 4 | 5.15E-65 | 218 | gag-polypeptide of LTR copia-type |
| A.pmajus002062.01 | 4 | 0 | 1231 | Heat shock protein |
| A.pmajus002063.01 | 4 | 8.51E-98 | 288 | Plectin/S10 domain |
| A.pmajus002090.01 | 4 | 6.56E-241 | 672 | Belongs to the UDP-glycosyltransferase family |
| A.pmajus010107.01 | 4 | 1.69E-66 | 221 | gag-polypeptide of LTR copia-type |
| A.pmajus009421.01 | 4 | 7.29E-53 | 172 | Belongs to the BetVI family |
| A.pmajus009425.01 | 4 | 7.29E-53 | 172 | Belongs to the BetVI family |
| A.pmajus018675.01 | 4 | 0 | 972 | Belongs to the pyruvate kinase family |
| A.pmajus018677.01 | 4 | 0 | 1324 | Bulb-type mannose-specific lectin |
| A.pmajus018690.01 | 4 | 8.71E-201 | 570 | Belongs to the UDP-glycosyltransferase family |
| A.pmajus018691.01 | 4 | 9.17E-116 | 332 | Sybindin-like family |
| A.pmajus018692.01 | 4 | 1.27E-211 | 590 | SGS domain |
| A.pmajus018695.01 | 4 | 0 | 1221 | Heat shock protein |
| A.pmajus028563.01 | 4 | 2.97E-131 | 381 | Mediator of RNA polymerase II transcription subunit |
| A.pmajus034712.01 | 4 | 0 | 1100 | STAS domain |
| A.pmajus034712.02 | 4 | 1.09E-98 | 308 | STAS domain |
| A.pmajus032591.01 | 4 | 5.34E-66 | 219 | gag-polypeptide of LTR copia-type |
| A.pmajus006943.01 | 5 | 5.15E-209 | 581 | Belongs to the iron ascorbate-dependent oxidoreductase family |
| A.pmajus030349.01 | 5 | 4.84E-100 | 297 | PLATZ transcription factor |
| A.pmajus004987.01 | 6 | 9.98E-80 | 239 | Heavy metal-associated isoprenylated plant protein |
| A.pmajus004989.01 | 6 | 4.16E-65 | 207 | SANT SWI3, ADA2, N-CoR and TFIIIB'' DNA-binding domains |
| A.pmajus004999.01 | 6 | 2.16E-25 | 100 | MLP-like protein |
| A.pmajus005002.01 | 6 |  |  |  |
| A.pmajus005009.01 | 6 |  |  |  |
| A.pmajus005010.01 | 6 | 1.66E-69 | 220 | Myb-like DNA-binding domain |
| A.pmajus029557.01 | 6 | 2.04E-102 | 320 | Belongs to the protein kinase superfamily |
| A.pmajus027617.01 | 6 | 0 | 1017 | Sterol 3-beta-glucosyltransferase UGT80A2-like |
| A.pmajus045006.01 | 6 |  |  |  |
| A.pmajus033889.01 | 7 | 1.06E-250 | 699 | Sugar (and other) transporter |

### **Supplementary Table 7: Linear model fit**

Coefficients for a linear model fit to the data of Marin et al. 2020. With altitude and its interaction with variety included as explanatory variables, variety alone does not significantly affect plant height, node or branch number.

| **Model formula** | **Response variable** | **Explanatory variable** | **Estimate** | **Std. Error** | **t value** | **Pr(>\|t\|)** |
| --- | --- | --- | --- | --- | --- | --- |
| LONG.TOT ~ COULEUR * ALT | Plant height | (Intercept) | 49.556712 | 1.751577 | 28.293 | <2.00E-16 |
|  |  | Variety | -3.044713 | 2.044504 | -1.489 | 0.137 |
|  |  | Altitude | -0.014228 | 0.002179 | -6.529 | 2.11E-10 |
|  |  | Variety:Altitude | 0.01518 | 0.002754 | 5.512 | 6.53E-08 |
| NOEUDS.FLO ~ COULEUR * ALT | # Nodes | (Intercept) | 13.41973 | 0.4306761 | 31.16 | <2.00E-16 |
|  |  | Variety | -0.165907 | 0.5075849 | -0.327 | 0.744 |
|  |  | Altitude | -0.0022256 | 0.0004977 | -4.472 | 9.19E-06 |
|  |  | Variety:Altitude | 0.0032675 | 0.0006367 | 5.132 | 3.81E-07 |
| RAMEAU.FLO ~ COULEUR * ALT | # Branches | (Intercept) | 16.9694208 | 0.808935 | 20.977 | <2.00E-16 |
|  |  | Variety | 0.9799639 | 0.953392 | 1.028 | 0.304403 |
|  |  | Altitude | -0.0030948 | 0.0009348 | -3.311 | 0.000984 |
|  |  | Variety:Altitude | 0.0023691 | 0.0011958 | 1.981 | 0.048004 |

### **Supplementary Table 8: Linear mixed model fit with population, family, and block effects**

Coefficients for a linear mixed model fit to the data of Marin et al. 2020. With population, family and container included as random effects according to the study's randomised block design, only altitude has a significant effect on plant height and branch number. The p-values in the last column were computed using Satterthwaites's method as implemented in R package "lmerTest".

| **Model formula** | **Response variable** | **Explanatory variable** | **Estimate** | **Std. Error** | **t value** | **Pr(>\|t\|)** |
| --- | --- | --- | --- | --- | --- | --- |
| LONG.TOT ~ COULEUR * ALT + (1 \| POP/FAM) + (1 \| BAC) | Plant height | (Intercept) | 45.135663 | 4.471108 | 9.552925 | 10.095 |
|  |  | Variety | 1.437692 | 5.348412 | 8.521022 | 0.269 |
|  |  | Altitude | -0.011513 | 0.004606 | 10.061138 | -2.499 |
|  |  | Variety:Altitude | 0.01051 | 0.006441 | 9.130928 | 1.632 |
| NOEUDS.FLO ~ COULEUR * ALT + (1 \| POP/FAM) + (1 \| BAC) | # Nodes | (Intercept) | 13.370747 | 1.132696 | 9.183197 | 11.804 |
|  |  | Variety | -0.033412 | 1.37616 | 8.711413 | -0.024 |
|  |  | Altitude | -0.002176 | 0.001174 | 9.893357 | -1.853 |
|  |  | Variety:Altitude | 0.003321 | 0.001645 | 9.03951 | 2.019 |
| RAMEAU.FLO ~ COULEUR * ALT + (1 \| POP/FAM) + (1 \| BAC) | # Branches | (Intercept) | 17.047105 | 1.424443 | 12.570752 | 11.968 |
|  |  | Variety | 0.720441 | 1.625528 | 9.370513 | 0.443 |
|  |  | Altitude | -0.003311 | 0.001435 | 12.078246 | -2.308 |
|  |  | Variety:Altitude | 0.002766 | 0.001962 | 10.105916 | 1.41 |

### **Supplementary Table 9: Linear mixed model excluding high altitude populations**

See Supplementary table 8, except populations with altitude > 1350m were excluded from the analysis.

| **Model formula** | **Response variable** | **Explanatory variable** | **Estimate** | **Std. Error** | **t value** | **Pr(>\|t\|)** |
| --- | --- | --- | --- | --- | --- | --- |
| LONG.TOT ~ COULEUR * ALT + (1 \| POP/FAM) + (1 \| BAC) | Plant height | (Intercept) | 44.200416 | 4.893524 | 8.735397 | 9.032 |
|  |  | Variety | 2.396778 | 5.772317 | 7.882842 | 0.415 |
|  |  | Altitude | -0.008757 | 0.006562 | 10.030797 | -1.335 |
|  |  | Variety:Altitude | 0.007775 | 0.008055 | 9.06336 | 0.965 |
| NOEUDS.FLO ~ COULEUR * ALT + (1 \| POP/FAM) + (1 \| BAC) | # Nodes | (Intercept) | 13.333735 | 1.260938 | 8.335218 | 10.574 |
|  |  | Variety | 0.001215 | 1.506439 | 7.905157 | 0.001 |
|  |  | Altitude | -0.002086 | 0.001704 | 9.867148 | -1.224 |
|  |  | Variety:Altitude | 0.003242 | 0.002092 | 8.906929 | 1.55 |
| RAMEAU.FLO ~ COULEUR * ALT + (1 \| POP/FAM) + (1 \| BAC) | # Branches | (Intercept) | 16.885113 | 1.572691 | 11.291779 | 10.736 |
|  |  | Variety | 0.882141 | 1.780694 | 8.707959 | 0.495 |
|  |  | Altitude | -0.002804 | 0.002119 | 12.985545 | -1.323 |
|  |  | Variety:Altitude | 0.002266 | 0.002543 | 10.86105 | 0.891 |

### **Supplementary Table 10: KASP / AFLP markers for islands 7 and 8**

| ***LOCUS*** | **Marker** | **Marker** | **Oligo** | **Oligo sequence** | **Oligo Position (*A. majus* reference genome)** |
| --- | --- | --- | --- | --- | --- |
|  | **Type** | **Name** | **Name** |  |  |
| ***island 7*** | KASP | Set 103 | do.752 | CTTCAAGGTCGTGGAGTGTTACG | Chr8:51,096,829-51,096,851 |
| ***(XAT)*** |  |  | do.753 | CTTCAAGGTCGTGGAGTGTTACA | Chr8:51,096,829-51,096,851 |
|  |  |  | do.754 | GTTCTGGTGCAAGGGAGGTAAGC | Chr8:51,096,777-51,096,799 |
| ***island 7*** | KASP | Set 110 | do.773 | ATTCTGGGCAGACACAATTGGG | Chr8:54,736,070-54,736,091 |
| ***(XAT)*** |  |  | do.774 | ATTCTGGGCAGACACAATTGGA | Chr8:54,736,070-54,736,091 |
|  |  |  | do.775 | TACTCCAGACCACTTCTTGAGAC | Chr8:54,736,028-54,736,050 |
| ***island 8*** | AFLP | Set 99* | do.636 | ACGTGTTCTCTTCTCTGCTCATGG | Chr6:40,704,294-40,704,317 |
| ***(ALTA1)*** |  |  | do.637 | CCGTGTCCATAATGAACGGTTGG | Chr6:40,704,710-40,704,688 |
| ***island 8*** | KASP | Set 92 | do.707 | CTCTACTTGCAGCCAGTTACT | Chr6:42,060,425-42,060,445 |
| ***(ALTA1)*** |  |  | do.708 | CTCTACTTGCAGCCAGTTACA | Chr6:42,060.425-42,060,445 |
|  |  |  | do.709 | CGTCCTCAAACACAACAATTGG | Chr6:42,060,488-42,060,467 |

*AFLP marker set 99 initially showed an excess of upper band (U) homozygotes. Individuals initially genotyped as U homozygotes were additionally genotyped with KASP marker set 92, which resolved U homozygotes into U/U and heterozygote classes.
